# Immunopeptidomics for Dummies: Detailed Experimental Protocols and Rapid, User-Friendly Visualization of MHC I and II Ligand Datasets with MhcVizPipe

**DOI:** 10.1101/2020.11.02.360958

**Authors:** Kevin A. Kovalchik, Laura Wessling, Frederic Saab, Qing Ma, Jérôme Despault, Peter Kubiniok, David Hamelin, Pouya Faridi, Chen Li, Anthony Purcell, Marco Tognetti, Lukas Reiter, Roland Bruderer, Joël Lanoix, Éric Bonneil, Mathieu Courcelles, Pierre Thibault, Etienne Caron, Isabelle Sirois

## Abstract

Immunopeptidomics refers to the science of investigating the composition and dynamics of peptides presented by major histocompatibility complex (MHC) class I and class II molecules using mass spectrometry (MS). Here, we aim to provide a technical report to any non-expert in the field wishing to establish and/or optimize an immunopeptidomic workflow with relatively limited computational knowledge and resources. To this end, we thoroughly describe step-by-step instructions to isolate MHC class I and II-associated peptides from various biological sources, including mouse and human biospecimens. Most notably, we created MhcVizPipe (MVP) (https://github.com/CaronLab/MhcVizPipe), a new and easy-to-use open-source software tool to rapidly assess the quality and the specific enrichment of immunopeptidomic datasets upon the establishment of new workflows. In fact, MVP enables intuitive visualization of multiple immunopeptidomic datasets upon testing sample preparation protocols and new antibodies for the isolation of MHC class I and II peptides. In addition, MVP enables the identification of unexpected binding motifs and facilitates the analysis of non-canonical MHC peptides. We anticipate that the experimental and bioinformatic resources provided herein will represent a great starting point for any non-expert and will therefore foster the accessibility and expansion of the field to ultimately boost its maturity and impact.

## INTRODUCTION

The importance of MS-based immunopeptidomics for the discovery of T cell targets against cancer and, more recently, pandemic pathogens, has attracted the interest of researchers from a wide range of scientific disciplines (1, 2). Despite the fact that the immunopeptidomics field has been historically limited to a handful of research groups, its constant gains in popularity may serve as an indicator of its trajectory toward large-scale deployment in the near future. In fact, the growing interest for immunopeptidomics was accelerated by the unquestionable contribution to the 2018 Nobel prize winners James P. Allison and Tasuku Honjo for their work on cancer immunotherapy as well as by the development of next-generation MS technologies and software tools for peptide identification and quantification (3–7). Furthermore, the development of MHC peptide databases [e.g. Immune Epitope Database (IEDB), HLA Ligand Atlas, and SysteMHC Atlas] has greatly contributed to the acceleration of data sharing and reflects the collaborative nature of the field (8–11). Akin to genomics, we anticipate that the immunopeptidomics field will keep growing in response to consistent community efforts and will reach its highest impact in biomedical research upon global accessibility and usage of mature protocols, tools, and technologies (12).

Several protocols and reviews in MS-based immunopeptidomics were recently published and they represent a reasonable starting point for any non-expert in the field aiming to conduct an initial immunopeptidomic experiment (2, 13–19). In addition to reviewing the literature, non-experts are encouraged to perform the following quality control experiments to validate the performance of their immunopeptidomic workflows: 1) grow standard cell lines (~10^9^ cells) expressing a high level of MHC class I and II molecules to use as positive controls [e.g. JY (20), B-LCL (21)], 2) apply the conventional immunoaffinity purification method, 3) measure the sampled peptides by conventional shotgun proteomics and 4) assess the quality and the specificity of the immunopeptidomic data by determining the proportion of peptides predicted to bind to the HLA haplotypes expressed by the cell lines. To this end, prediction algorithms such as NetMHCpan (22, 23) and MHCflurry (24, 25) are recommended. Other tools such as GibbsCluster (26) and MoDec (27) are also freely available and are useful to identify MHC binding motifs from peptide lists. Once the experimental workflow is validated based on these quality control steps, the users can move on and test samples of interest such as neoplastic tissues, cell lines, etc. When moving from standard cell lines to samples of interest, assessing the quality of the generated immunopeptidomic data is critical for the success of any study. However, the bioinformatic tools described above can process only one sample at a time and some were not even initially designed for immunopeptidomics applications. Alternatively, specialized immunopeptidomics software tools (e.g. MHCquant and MAPDP) were created for the analysis of canonical and non-canonical MHC-associated peptides from raw MS data but may remain relatively challenging to install and operate for non-experts with limited computational background and resources (28, 29). Hence, the development of an easy-to-install and easy-to-operate software tools for the rapid assessment of the quality and the specificity of immunopeptidomic datasets are needed and will find utility for any non-expert wishing to embark effectively into the immunopeptidomics’ journey!

In this technical report, we present MhcVizPipe (MVP), a new open-source and freely available software tool intended to be effortlessly used by any laboratory for rapid visualization of multiple immunopeptidomic datasets. Here, we describe how to install and run MVP (Figure 1 and **Supplemental data S26** and https://github.com/CaronLab/MhcVizPipe), the content of an MVP report (Figure 2 and **Supplemental data S1-S3**), as well as specific applications (Figures 3–4 and **Supplemental data S5**-S20 and **Table S1-S4**). To help non-expert laboratories to establish effective immunopeptidomic workflows, we also thoroughly reviewed and tested the most common immunoaffinity purification protocols (13–19). As a result, we provide herein a detailed protocol describing step-by-step instructions for any beginner in the field. The detailed protocol describe the most critical steps and reagents to consider while testing and optimizing the immunoaffinity purification procedure for the isolation of MHC class I and class II-associated peptides, from both mouse and human biological materials (Supplemental data S21-**S24**). Finally, we present applications of MVP to assess the specificity and sensitivity of the tested sample preparation protocols and to explore immunopeptidomics datasets for the analysis of both canonical and non-canonical MHCI-associated peptides.

**Figure 1.**
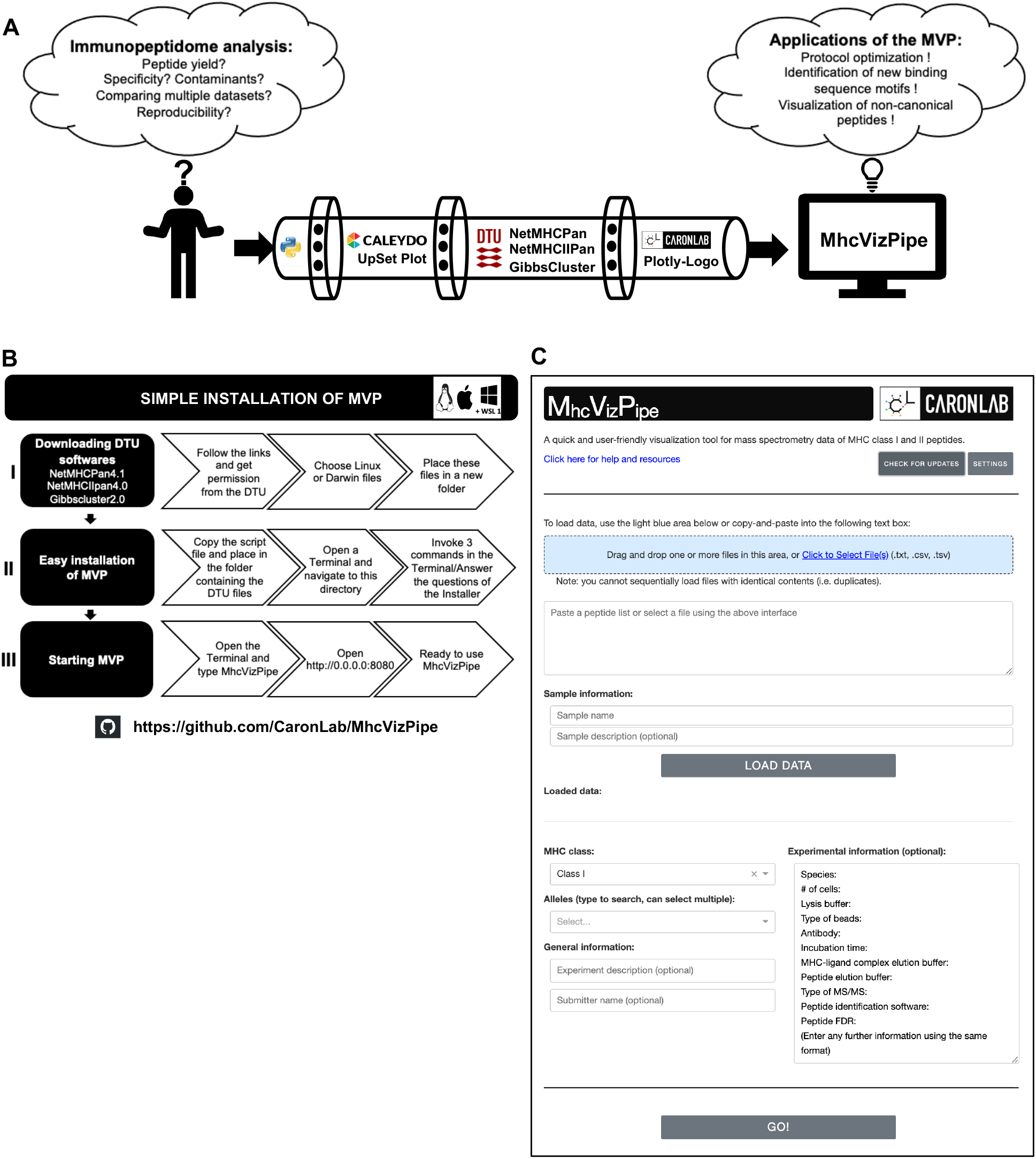
Overview and installation of the MVP pipeline. **A**. Schematic of the MVP pipeline content. The pipeline was created with the Python language and connects 4 algorithms available from the internet (i.e. UpSet Plot, NetMHCPan, NetMHCIIPan and GibbsCluster) and our in-house Plotly-Logo algorithm. **B.** Installation steps of MVP. The user can download the simple installation from http://github.com/CaronLab/MhcVizPipe on both Linux, Mac operating systems and on a Windows machine enabling a Linux subsystem (WSL 1). The installation requires 3 small blocks of operation: block I: downloading third-party softwares, block II: Easy installation of MVP and block III: Starting MVP. **C.** Overview of the MVP graphic user interface.

**Figure 2.**
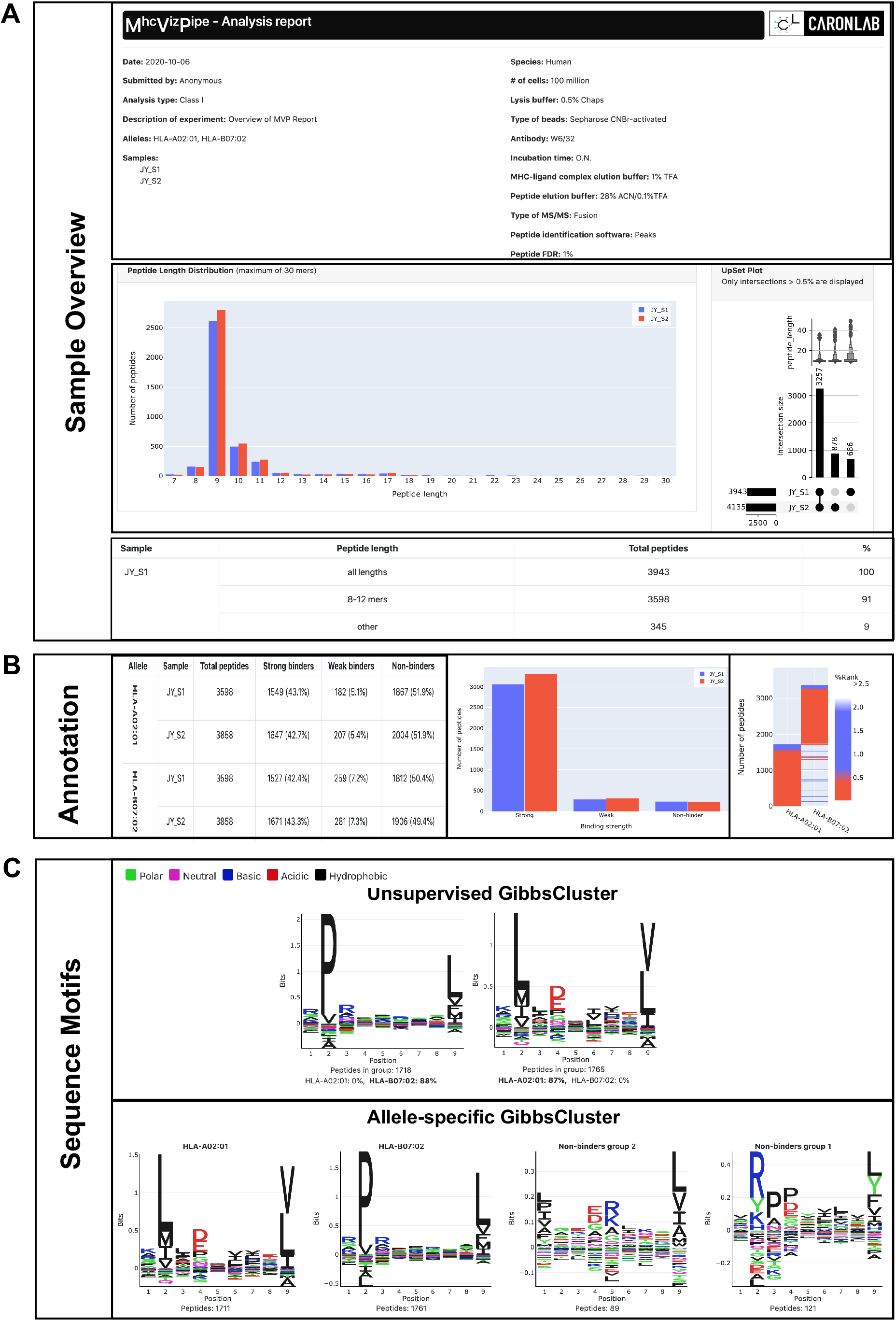
Overview of the MVP report. Two replicates of MHC class I immunopeptidomes isolated from 100 million JY cells were used to present the HTML report laid out in three sections: **A.** Visual the of the Sample Overview section from the MVP report. The heading of the report includes experimental and samples details. A table with peptide counts is shown below the peptide length distribution graph (left) and the UpSet Plot (right). **B.** Visual of the Annotation results section obtained from NetMHCpan (I or II). Left: descriptive table of the proportion of total number, strong, weak and non-binding MHC peptides. Middle: Graph showing the number of peptides in function of binding affinity. Right: heatmap showing the number of peptides with binding affinity scores for each allele analyzed. **C.** Visual of the Sequence motifs section from the GibbsCluster analysis presented in two ways. Upper part: Unsupervised tab displays the most significant motif(s) represented by the subset of peptides (i.e. 8-12 and 9-22 amino acids for MHC class I and II, respectively) in the sample along with the % of peptides associated to each allele. Lower part: the allele-specific cluster tab displays peptide motif(s) generated from strong and weak MHC binders as well as motifs from peptides predicted to be non-assigned MHC binders.

**Figure 3.**
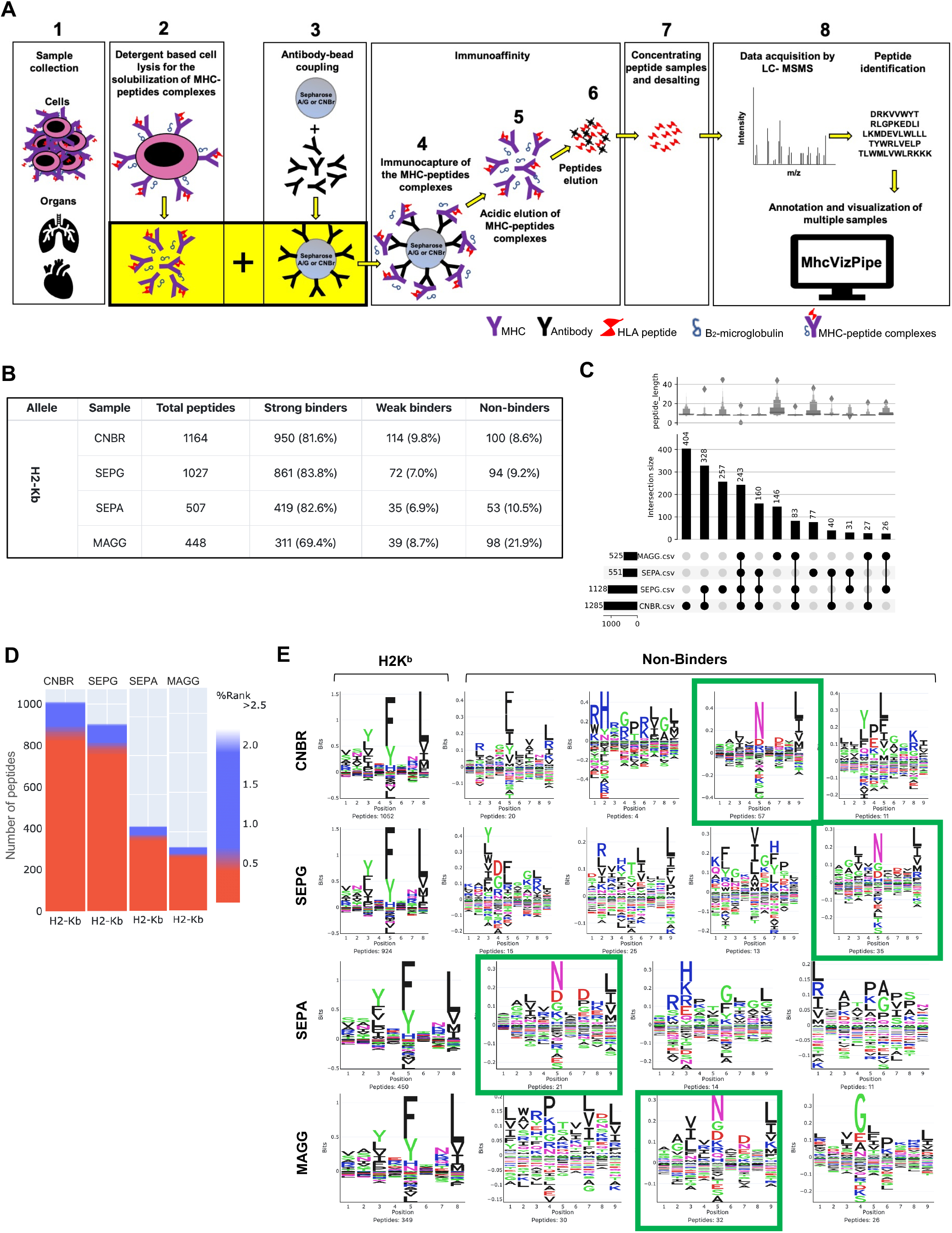
Examples of applications of MVP. **A.** Schematic of the complete procedure for the isolation of MHC peptides (1-7), data acquisition, peptide identification and peptide annotation and visualization using MVP (8). **B-D.** MVP was used to show the impact of using different beads to isolate MHC class I peptides from EL4 cells. **B.** Annotation table showing the total number of peptides and the proportion of peptides with different binding affinities. **C.** The UpSet plot showing the number of unique and shared MHC class I peptides isolated from EL4 mouse cancer cells using different types of Sepharose beads. **D.** Heatmaps representing the number of peptides in function of binding affinity scores of peptides analyzed in (B). **E.** MHC binding motifs of H2K^b^ MHC class I peptides visualized with the allele-specific GibbsCluster tab revealed the presence of other MHC binding motifs (H2D^b^, green boxes) in the analyzed samples.

**Figure 4.**
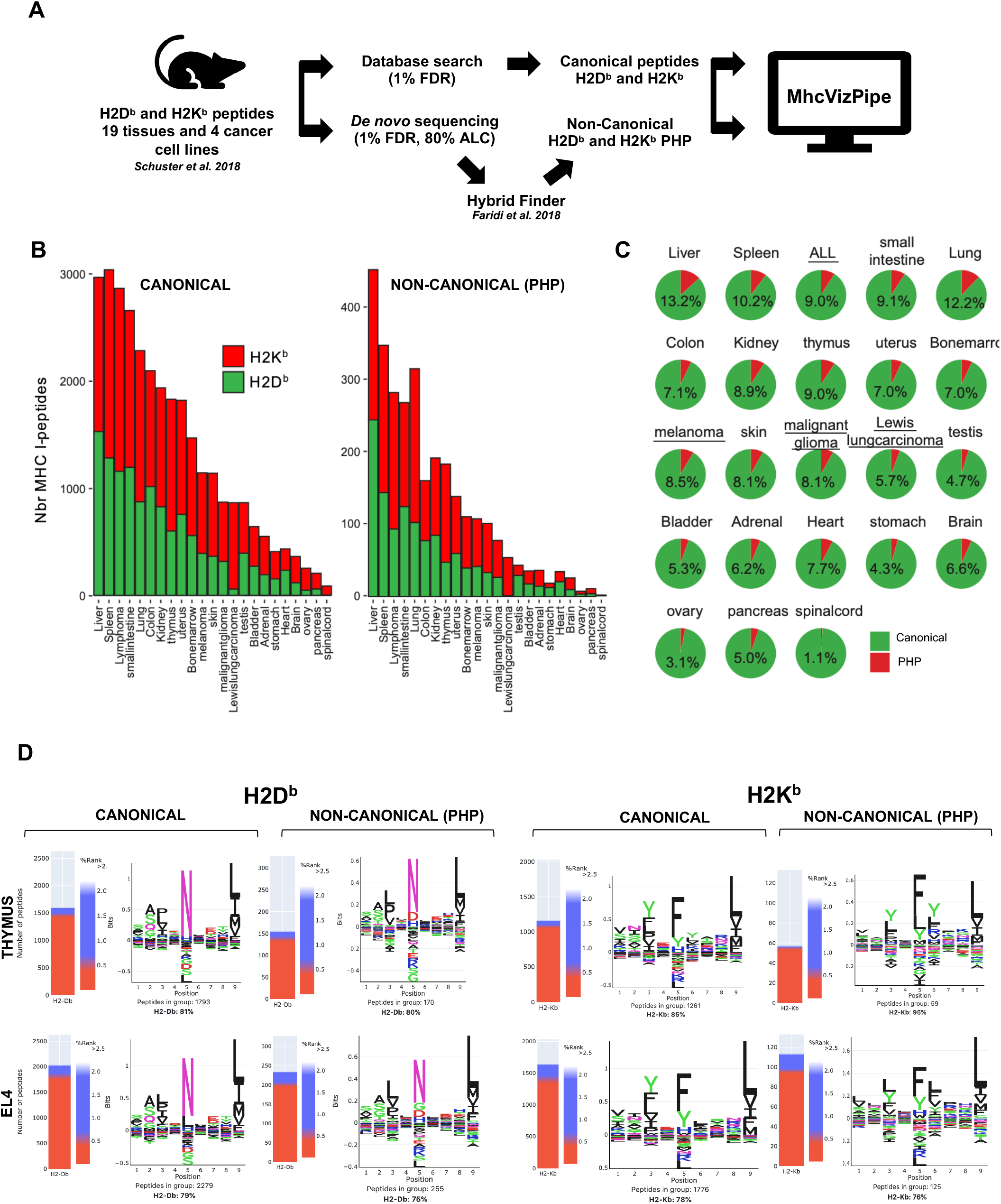
Visualization of non-canonical peptides with MVP. **A.** Schematic for the identification of canonical and non-canonical peptides (PHP) from H2K^b^ and H2D^b^ immunopeptidomes isolated from 19 mouse tissues and 4 cancer cell lines. Each dataset was analyzed using database search and *de novo* sequencing approaches with PEAKS X. To identify non-canonical PHP, peptides identified uniquely from the *de novo* approach were analyzed using the Hybrid Finder software. All generated data were visualized with MVP. **B.** Graphical distribution of the number of canonical and non-canonical PHP associated to H2K^b^ and H2D^b^ across different murine primary tissues and tumor cell lines. **C.** Pie chart distribution (%) of canonical peptides (green) and non-canonical PHP (red). Tumor cell lines are underlined. **D.** Binding affinity heatmaps and peptide binding motifs of canonical peptides and non-canonical PHP associated to H2D^b^ and H2K^b^. Thymus and EL4 are shown as representative examples.

## EXPERIMENTAL PROCEDURES

### Cell culture, cell viability, and reagents

A20 (BALB/c B cell lymphoma line derived from a spontaneous reticulum cell neoplasm found in an old BALB/cAnN mouse), EL4 (mouse thymic lymphoma cell line), JY (human lymphoblastoid B-cell line) were obtained from ATCC and B-LCL (lymphoblastoid B-cell from obtained from(21). Cell culture media: EL4 cell line was cultured in X-vivo 15 (04-744Q, Lonza) media supplemented with 25% DMEM (319-005-CL, Multicell), 10% FBS (98150, Multicell) and 1% pen/strep (450-201-EL, Multicell). The JY cell line was cultured in DMEM supplemented with 10% FBS and 1% pen/strep. The A20 cell line was cultured in RPMI 1640 1X (350-000-CL, Multicell) supplemented with 10% FBS, 1% pen/strep, 1% sodium pyruvate 100mM (100X)(600-110-EL, Multicell), 1% Hepes 1M solution (330-050-EL), 1% D-glucose (0188-500G, Amresco) and 0,1% Beta-mercaptoethanol (190242, MP Biomedicals). Refer to the **Supplemental Data S22a** for more details about cell subculturing. Cell viability was measured by an Invitrogen countess II automated cell counter (AMQAX1000) using 10 μl of culture media mixed with 10μl of 0.4% trypan blue (EB7-001, NanoEntek). The final cell concentration was measured using the mean of both counting chambers of the cell slide. The following antibodies were used for the Immunopurification (all from BioXcell): Mouse M5/114 Anti-MHC class II (BE00108), Mouse Y3 anti-H2Kb (BE0172), Human W6/32 Anti-HLA A, B, C (BE0079). Polyprep chromatography column (#7311553, Bio-Rad), Protein G Sepharose 4 fast flow (#17-0618-01, Fisher), CNBr activated sepharose 4B (#45000066, Fisher), Dynamag-2 (#12321D, Invitrogen), 1.5 ml Protein LoBind from Eppendorf, (022431081, Fisher), 2 ml Protein LoBind from Eppendorf (# 02243100, Fisher), 2.0 ml Costar microfuge tubes, 2 ml (3213, Corning) 15 ml Falcon tubes (352096, Corning), Solid phase extraction disk, ultramicrospin column C18 (The nest group, #SEMSS18V, 5-200 ul), Low retention tips: EPLORET reload tip 10, 200, 1000 ul (#2717349, #2717351, #2717352, Fisher). Acetonitrile (#A9964), trifluoroacetic acid (TFA, #AA446305Y), formic acid (#AC147930010), chaps (#22020110GM), PBS (Buph, phosphate buffer saline packs, #28372), ammonium bicarbonate (#A643-500), sodium deoxycholate (#89905), octyl-beta-d glucopyranoside (#AAL1325903) and Phenyl-methylsulfonyl fluoride (#AAB2214603) were purchased from Fisher. Combined inhibitor EDTA-free (#A32961, Bio-Rad).

### Experimental Design and Statistical Rationale

This technical report describes a new bioinformatic pipeline to visualize immunopeptidomics datasets. Therefore, the data shown in the results sections and the supplementary data files are indented to be used as examples of application of the MVP rather than biological findings. Therefore, the notion of biological and technical replicates is not applicable in this context.

### Cell lysis

Cell pellets were gently thawed and resuspended with 0.5 ml of cold PBS. The total volume (cells in PBS) was measured and the equivalent volume of 2x cell lysis buffer was added. Two types of lysis buffer were used: 1% chaps in PBS (2x) or NaODEOXY Lysis buffer as described (14): 0.5% sodium deoxycholate, 0.4 mM iodoacetamide, 2 mM EDTA, 1 pellet/5 ml of combined inhibitor EDTA-free, 2 mM Phenyl-methylsulfonyl fluoride, 2% octyl-beta-d glucopyranoside in PBS. Cell lysates were incubated and rotated for 1 hour at 4 degrees Celsius using the Revolver apparatus (20 RPM). Cleared cell lysates were harvested following 20 minutes by centrifugation at 17927 RCF (13000 RPM) at 4 degrees Celsius (fixed rotor). Supernatants were transferred into new 2.0 ml tubes and kept on ice.

### Bead coupling and immunopurification

Bead coupling and immunopurification protocols are described in full length in the **Supplemental data S22-S24**. Briefly, Sepharose-A, -G, or CNBr activated beads or magnetic beads were coupled with 1 or 2 mg of antibody. Sepharose antibody-coupled beads were incubated with the cell lysate overnight in a Bio-Rad column at 4 degrees Celsius using a slow rotating wheel Revolver apparatus (22 RPM). The next day, Bio-Rad columns were installed on a rack, and the bottom cap was removed to empty the unbound cell lysate. Beads were washed sequentially with 10 ml of buffer A (150 mM NaCl and 20 mM Tris–HCl pH 8), 10 ml of buffer B (400 mM NaCl and 20 mM Tris–HCl pH 8), 10 ml of buffer A and 10 ml of buffer C (20 mM Tris–HCl pH 8.). MHC-ligand complexes were eluted from the beads with 2 × 300 ul of 1% TFA (or as indicated) and kept on ice. MHC-ligands were eluted using C18 columns pre-conditioned with 200 ul of 80% acetonitrile/0.1%TFA and spin at 1545 RCF (3700 RPM) in a fixed rotor. The flowthrough was aspirated and 200 ul of 0.1% TFA was added to the column and spun again. 3 × 200 ul of the MHC-ligand complexes solution were loaded into the column followed by a wash (200 ul) with 0.1% TFA and spin. The stage tips were transferred onto a 2.0 ml Eppendorf tube and peptides were eluted with 3 × 200 ul of 28%ACN-0.1%TFA. The flowthrough containing the eluted peptides was stored at −20 degrees Celsius for MS analysis. For the protocol with the magnetic beads, refer to **Supplemental data S23f-g**.

### Mass spectrometric MSMS analysis and peptide identification

Samples were solubilized in 5% ACN-4% formic acid (FA). Peptides were loaded and separated on a home-made reversed-phase column (150-μm i.d. by 250 mm length, C18 Jupiter 3 μm C18 300 Å) with a gradient from 10 to 30% ACN-0.2% FA and a 600-nl/min flow rate on an Easy nLC-1000 connected to an Orbitrap Fusion (Thermo Fisher Scientific, San Jose, CA). Each full MS spectrum was acquired at a resolution of 120,000 an AGC of 5E5 and an injection time of 50 ms. It was followed by tandem-MS (MS-MS) spectra acquisition on the most abundant multiply charged precursor ions for a maximum of 3s. Tandem-MS experiments were performed using higher energy collisional dissociation (HCD) at a collision energy of 32%, a resolution of 15000 an AGC of 2E4, and an injection time of 1000ms. Data files were processed using PEAKS X (Bioinformatics Solutions, Waterloo, ON) using the mouse database UniProtKB/Swiss-Prot (2019_09) and human database UniProtKB/Swiss-Prot (2019_11). ‘Unspecified enzyme digestion’ was selected for the enzyme parameter and mass tolerances on precursor and fragment ions were 10 ppm and 0.01 Da, respectively. Variable modifications were deamidation (NQ) and oxidation (M). All other search parameters were the default values. Final peptides lists were filtered with a false discovery rate (FDR) of 1 % using the Peaks software. The identification of non-canonical peptides from *de novo* sequencing results (using ALC of 80%) was done using the Hybrid Finder pipeline, as described (30).

### Workflow and Installation of MhcVizPipe

MVP was built in the Python programming language. It connects the bioinformatics tools NetMHCpan (v4.0 or 4.1), NetMHCIIpan (v4.0) and GibbsCluster (v2.0) with an intuitive graphical user interface and generates organized and easy-to-understand reports in a portable HTML format. The computational workflow of the pipeline is described in detail in the **Supplemental Data S25**. Briefly, peptide lists are first stripped of peptides containing chemical modifications. The complete lists are then used for generating the length histogram and UpSet plot in the “Sample Overview” section of the report. The lists are then subset to peptides 8-12 or 9-22 mers in length for class I or class II peptides, respectively. Binding predictions for these subsets are made using NetMHCpan4.0 or 4.1 or NetMHCIIpan4.0 and are used to generate the “Binding Results” and “Annotation Heatmaps” sections in the report. Peptide grouping and alignment is performed by GibbsCluster twice, once using the complete subset of peptides (yielding the results in the “Unsupervised GibbsCluster” tab) and then again for the sets of peptides predicted to bind the different alleles (yielding the results in the “Allele-Specific GibbsCluster” tab). Peptide motifs for the GibbsCluster are generated using our in-house Plotly-Logo program (https://github.com/kevinkovalchik/Plotly-Logo). MVP is freely available at https://github.com/CaronLab/MhcVizPipe and can be downloaded and installed on Linux (e.g. Ubuntu), Mac and Windows operating systems (Installation on Windows requires the Windows Subsystem for Linux). A detailed explanation of the installation process and requirements can be found on the MVP GitHub wiki: https://github.com/CaronLab/MhcVizPipe/wiki and **Supplemental Data S26.** MVP installation was tested on the following systems: Ubuntu 18.04 and 20.04, MacOS 10.13 HighSierra, MacOS 10.14.16 Mojave and Windows 10 using the Windows Subsystem for Linux, WSL 1.

## RESULTS

### Overview and installation of the MhcVizPipe graphic user interface

The foremost goal behind the making of MVP is to provide a rapid and effective tool for the assessment of the quality and the specific enrichment of MHC-peptides datasets. In order to assess and compare simultaneously multiple MHC class I and II datasets, several measurable parameters specific to immunopeptidomic studies should include: 1) the number of peptides identified, 2) the proportion of peptides with the expected peptide length, 3) the predicted HLA binding affinity score according to the HLA subtype of the sample analyzed, and 4) the peptide binding motif. To achieve this, we created MVP, a bioinformatic analysis and visualization pipeline built-in the Python programming language, that is combining different software tools including Upset plot, NetMHCpan, NetMHCIIpan and Gibbclusters (23, 31–33) (Figure 1A).

MVP is freely available at https://github.com/CaronLab/MhcVizPipe and can be downloaded and installed on Linux (e.g. Ubuntu), Mac, and with the newest updates of Windows 10, it is easy to operate on a Windows machine while enabling a Linux subsystem (Figure 1B and **Supplemental Data S26**). A detailed explanation of the installation process can be found on the MVP GitHub wiki: https://github.com/CaronLab/MhcVizPipe/wiki. Briefly, two installation options are available: 1) a “simple” automated option for absolute beginners (Figure 1B), and 2) a step-by-step option intended for situations in which the simple option fails (e.g. due to issues encountered in an untested OS) or when the user wishes to use an existing Python environment or existing installations of NetMHCpan, NetMHCIIpan and GibbsCluster. To ease the installation process for those with limited technical experience, the automated installation of MVP includes a standalone distribution of Python 3.7.9. MVP runs as a graphical user interface (GUI) in any web browser (tested in Firefox, Safari, Chrome, Chromium and Edge) (Figure 1C). Because MVP is run as a web server, it can be accessed by any computer on the local network, facilitating use by multiple users and even by users with Windows computers.

The proper installation of MVP can be tested using peptide lists available on the GitHub page. Peptide lists can be uploaded in different formats (.csv, .tsv or .txt) or copy-pasted into the GUI. Any file of the mentioned formats may be uploaded provided they have columns headers. If a multi-column file is opened (e.g. database search results in .csv format), the user is asked by MVP to select the header of the column which contains the peptide sequences. There is no limit to the number of files which can be analyzed at one time, though processing time increases with the number of files. Once the peptide lists have been uploaded, the species, type of MHC class (I or II), and alleles corresponding to the sample(s) are specified. Many technical details related to the samples can also be specified and will appear in the final report, which is generated on a portable HTML format and can be viewed in the browser or saved to the computer. A resource tab is also available containing details related to MVP as well as useful information and resources.

### Content of the MhcVizPipe report

The HTML report generated from the MVP graphic user interface contains three main sections: 1) Sample Overview, 2) Annotation Results and 3) Sequence Motifs. The Sample Overview (Figure 2A) section regroups a peptide length distribution graph for a rapid assessment of the quality of the immunopeptidome since a specific preparation will lead to a specific length distribution. The descriptive peptide count table indicates the number of peptides corresponding to the expected length related to the MHC class selected. Finally, an “UpSet” plot (31) shows the number of unique and shared peptides between multiple samples. Also, the reproducibility among replicates and the presence/absence of contaminant peptides can be easily estimated from the presented graph.

The second section of the report focuses on annotation of predicted MHC binding affinity values for individual peptides based on the provided MHC alleles annotation and could serve as an index of sample specificity (Figure 2B). For this section, specific peptides lengths were considered since MHC class I and II ligands display a specific range of amino acid length based on both their different mechanisms of production and the structural properties of the MHCs (34, 35). In line with this, peptides for the annotation section are subset to those between 8- and 12-amino acid in length for class I and between 9- and 22 for class II.

According to the class of the peptides to be analyzed, peptide lists are processed by NetMHCpan (v4.0 or 4.1) for class I peptides, or NetMHCIIpan4.0 for class II peptides. The annotation predictors use the eluted ligand (EL) scores generated by NetMHCpan to define strong binders (rank <= 0.5), weak binders (2.0 <= rank >= 0.5) or non-binders (rank > 2.0) (see Supplemental data S19–S20 for details about EL and why it was chosen over binding affinity (BA)). The report for this annotation section contains a peptide count table for each allele, a binding affinity graph distribution, and the corresponding heatmap showing the proportion of strong, weak, and non-MHC binders in the measured samples (Figure 2B). This information is quite useful to assess the specificity of the peptides identified; it is expected that a sample with a larger proportion of strong binders to the HLA subtypes associated to the sample would be a good indicator of sample specificity.

The third part of the report focuses on the identification of sequence motifs that are enriched in the analyzed samples using GibbsCluster and our in-house Plotly-Logo algorithm developed to facilitate MVP usage. The sequence motifs from the GibbsCluster analysis are presented in two ways: 1) the unsupervised tab displays the most significant motif(s) represented by the subset of peptides (i.e. 8-12 and 9-22 amino acids for MHC class I and II, respectively) in the sample along with the % of peptides associated to each allele, and 2) the allele-specific cluster tab displays peptide motif(s) generated from strong and weak MHC binders as well as motifs from peptides predicted to be non-assigned MHC binders (Figure 2C).

An interesting use of these sequence motifs results is the rapid recognition of classical or even undescribed binding motifs in the analyzed samples. For instance, the MHC class I H2K^b^ binding peptides are typically enriched with a phenylalanine (F) residue in the middle of the binding motif whereas H2D^b^ peptides are enriched with an asparagine (N), as shown in **Supplemental Data S11** and **S13**. These binding motifs are hallmarks of H2^b^ class I mouse peptides and can be straightforwardly seen in the MVP report under both Unsupervised and Allele-specific GibbsCluster tabs. Of note, one great advantage of the allele-specific cluster tab is the possibility to observe new groups of peptides with a significant binding motif that are either unassigned to any allele yet or indicative of contaminant peptides. To demonstrate the utility of MVP for non-expert, representative datasets of human class I and II peptides were used to generate several HTML reports as examples (**Supplemental data S1-S3** and **Table S1**). In summary, MVP generates in a few clicks a complete report with an overview of the sample composition as well as specific characteristics of MHC peptide ligands identified by MS.

### Applications of the MVP for protocol optimization and the assessment of data quality and specificity of mouse and human immunopeptidomes

MVP finds great utility in immunopeptidomics for the testing and optimization of sample preparation protocols. As frequently requested by non-experts, we provide in **Supplemental data (S22-S24)** a detailed protocol for the isolation of MHC class I and II peptides. The overall sample preparation and immunoaffinity procedure consist of eight main steps (Figure 3A, Table 1 and **Supplemental data S22-S24**). Importantly, the antibody used has a critical impact on the feasibility of a study and the quality of the immunopeptidomic data generated. While the W6/32 antibody is widely used in the field, most of the other antibodies documented for the immunoaffinity purification of MHC-peptide complexes have not been widely adopted by the community.

**Table 1.**
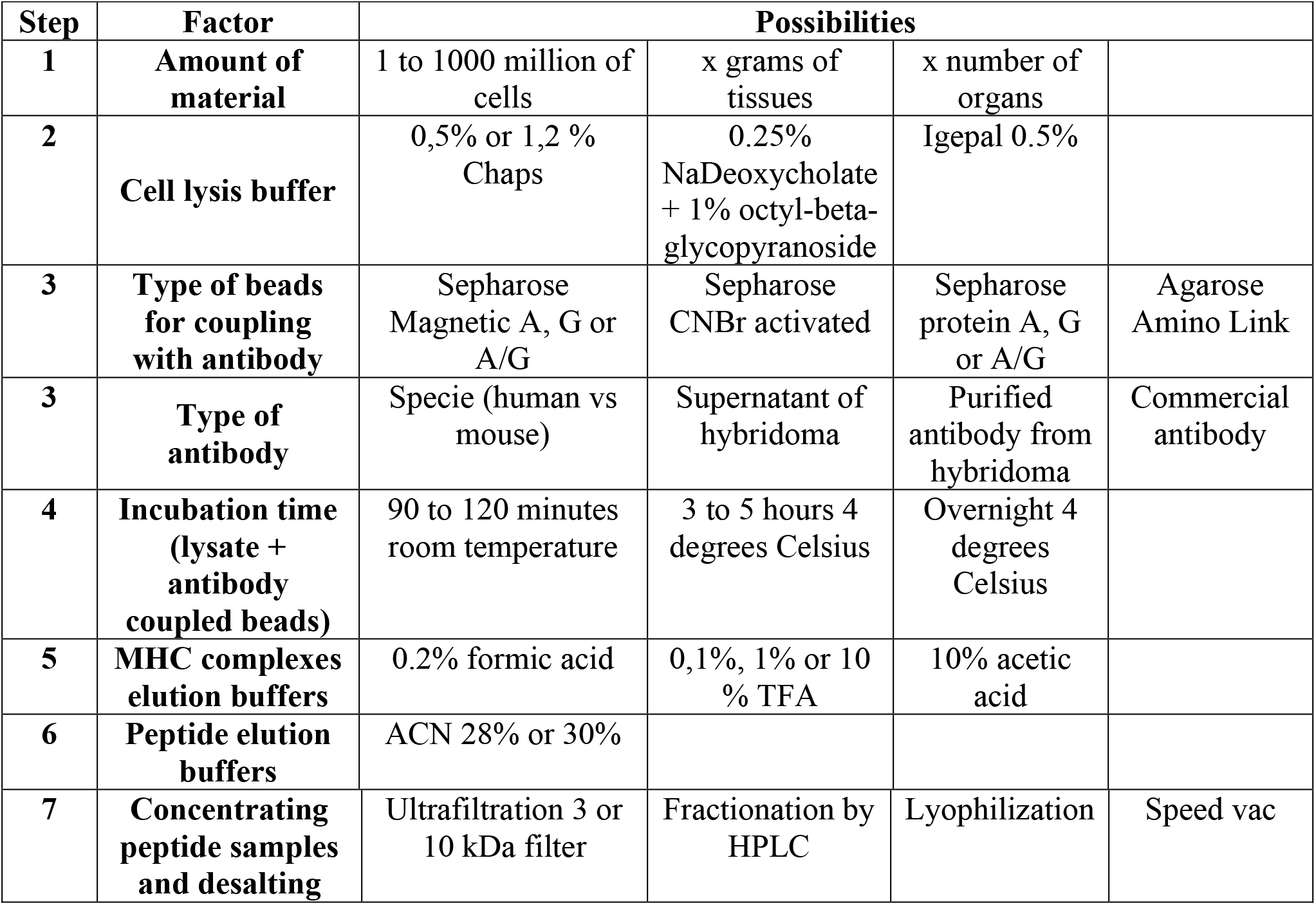
Factors influencing sensitivity and specificity of immunopeptidomes isolated by immunoprecipitation. Table showing every step and some associated variables related to the purification of immunopeptidomes reviewed from the literature (13, 14, 16, 54, 64, 65).

| Step | Factor | Possibilities |  |  |  |
| --- | --- | --- | --- | --- | --- |
| 1 | Amount of material | 1 to 1000 million of cells | x grams of tissues | x number of organs |  |
| 2 | Cell lysis buffer | 0,5% or 1,2 % Chaps | 0.25% NaDeoxycholate + 1% octyl-beta-glycopyranoside | Igepal 0.5% |  |
| 3 | Type of beads for coupling with antibody | Sepharose Magnetic A, G or A/G | Sepharose CNBr activated | Sepharose protein A, G or A/G | Agarose Amino Link |
| 3 | Type of antibody | Specie (human vs mouse) | Supernatant of hybridoma | Purified antibody from hybridoma | Commercial antibody |
| 4 | Incubation time (lysate + antibody coupled beads) | 90 to 120 minutes room temperature | 3 to 5 hours 4 degrees Celsius | Overnight 4 degrees Celsius |  |
| 5 | MHC complexes elution buffers | 0.2% formic acid | 0,1%, 1% or 10 % TFA | 10% acetic acid |  |
| 6 | Peptide elution buffers | ACN 28% or 30% |  |  |  |
| 7 | Concentrating peptide samples and desalting | Ultrafiltration 3 or 10 kDa filter | Fractionation by HPLC | Lyophilization | Speed vac |

To exemplify the usefulness of MVP for good protocol optimization, we applied MVP to report evidence that the W6/32 antibody performs equally well independently of the lysis buffers, the type of beads, and the incubation times tested (**Supplemental data S4, S6, S7**). In contrast, the performance of other commercially available antibodies for the isolation of mouse MHC class I molecules (Y3 antibody for H2K^b^) and MHC class II molecules (M5 antibody for H2-IAd and H2-IEd) were drastically affected by the type of beads and acidic buffers that were applied (Figure 3B-D and **Supplemental data S9-S10**). In fact, while the specificity (i.e. the proportion of strong H2K^b^ binders) of the datasets was relatively similar between the various types of beads tested, the sensitivity (i.e. the total number of peptides identified) was strongly affected. For instance, Sepharose-CNBR beads (CNBR) and Sepharose-G beads (SEPG) yielded 1164 (82% strong H2K^b^ binders) and 1027 (84% strong H2K^b^ binders) peptides, respectively, whereas Sepharose-A beads (SEPA) and Magnetic-G beads (MAGG) resulted in a ~2-fold decrease in peptide yield, i.e. 507 (83% strong H2K^b^ binders) and 448 (69% strong H2K^b^ binders) peptides, respectively (Figure 3B-C). Thus, all the beads tested were able to isolate peptides with the same specific dominant H2K^b^ binding motif from EL4 cells but in different proportions, with CNBR being the most efficient condition as shown using the allele-specific GibbsCluster tab (Figure 3D). Notably, the low number of peptides observed using MAGG (Figure 3C, D) correlated with the poor binding of Y3 antibody to MAGG compared to CBNR and SEPG, as visualized by Coomassie blue staining (Supplemental data S21). Together, these results indicate that the antibody-bead combination can considerably influence the overall performance of the immunoaffinity purification procedure. These results also highlight the utility of Coomassie blue staining during protocol optimization, when using new antibodies in particular (Supplemental Data S23c **and** e).

In addition, the MVP allele-specific GibbsCluster tab showed that the Y3 antibody documented to be specific against the classical H2K^b^ molecules also cross-reacted with H2D^b^, resulting in the identification of H2D^b^-associated peptides. This observation was made from the significant enrichment of peptides with a sequence motif sharing an asparagine (N), which corresponds to the anchor residue that is necessary to bind H2D^b^ (see green squares in Figure 3E). Finally, MVP was also useful to illustrate the isolation of a poor quality immunopeptidome as shown by the divergent peptide length distribution of the class II peptides eluted using different acidic conditions, i.e. 10% TFA versus three other conditions (1% TFA, acetic acid and formic acid) (**Supplementary data S10**). Taken together, these examples highlight that protocol optimization and data visualization using MVP could be done effortlessly to show the robustness of the W6/32 antibody and the cross-reactive nature of the Y3 antibody, which would have gone unnoticed without MVP. Hence, MVP considerably accelerate the analysis process for assessing the quality of immunopeptidomic datasets from any biological sources and permits a fast comparison of different protocols, thereby facilitating protocol optimization.

### Application of MVP for the analysis of non-canonical mouse immunopeptidomes

In the above section, MVP was applied for the analysis of canonical peptides presented by MHC molecules. Those canonical peptides derive from exome reading frames and are thought to be the most prominent population of peptides in the immunopeptidomic landscape of the cell (12). Beside canonical peptides, recent immunopeptidomics studies have reported the presentation of non-canonical MHCI-associated peptides, including those that derive from non-coding genomic regions, untranslated and out-of-frame coding regions, as well as post-translationally modified peptides (36–39). Among the latter, proteasome-catalyzed spliced peptides are particularly intriguing (39–41). In fact, their presence was reported in several studies and was shown to be targeted by T cells in both mouse and human disease models (42–45). However, their exact proportion remains debated as a result of the immaturity of the currently available methodologies in immunopeptidomics (46–51). Since most proteasome-catalyzed spliced peptides that have been systematically discovered by MS remain to be experimentally validated, we carefully referred them herein as ‘Potential Hybrid Peptides’ (PHP) (Figure 4).

Until now, the systematic investigation of the PHP immunopeptidome has mainly been performed in human cells. In fact, very little is known about those non-canonical peptides in other species (52, 53). Here, we sought to apply MVP in combination with PHP discovery-enabling computational tools to rapidly visualize the quality and proportion of PHP in 19 primary mouse tissues and four common tumor mouse cell lines: EL4 (lymphoma), B16F10 (melanoma), GL261 (malignant glioma) and LLC1 (LL/2) (Lewis lung carcinoma). More specifically, we used i) raw MS data available from the tissue-based atlas of the mouse MHC I immunopeptidome (54), ii) a recently described *de novo* sequencing-based computational workflow (Hybrid Finder) (49), and iii) our MVP tool for effective visualization of mouse PHP (Figure 4A). By using these various computational and data resources, we detected PHP in all the 19 murine tissues and the four murine tumor cell lines tested (Figure 4B). The average proportion of PHP was ~7% across all primary mouse tissues and cell lines tested (Figure 4C). Next, MVP was used to evaluate whether those PHP had similar MHC binding affinities and motifs compared to the canonical H2K^b^ and H2D^b^ peptides. Interestingly, the MVP plots revealed that PHP were preferably enriched with strong binders for H2D^b^ and H2K^b^ (Figure 4D). Accordingly, unsupervised clustering showed that PHP had dominant MHC peptide binding motifs in most murine tissues and cell lines tested (Figure 4D, **Supplemental data S11-S18** and **Tables S2-S4**). Further investigation is required to determine the exact origin of those PHP. Nonetheless, these results clearly highlight the suitability of MVP in assessing the MHC-specificity of non-canonical immunopeptidomic datasets generated by MS.

## DISCUSSION

Robust sample preparation protocols are frequently looked for by non-experts aiming at establishing an immunopeptidomic workflow. To support this need, several detailed protocols for the isolation of MHC-associated peptides have recently been reported (13–19). Nevertheless, those protocols include multiple steps and remain relatively challenging to establish in order to generate high-quality immunopeptidomic data. To support and facilitate the establishment and optimization of those protocols, we created MVP, a new and easy-to-use software tool to rapidly assess the data quality and specificity of any immunopeptidomic experiment. Furthermore, using the detailed sample preparation protocols provided will further help in the establishment of such a pipeline. We applied MVP to demonstrate the robustness of the W6/32 antibody as well as the necessity to optimize the immunoaffinity purification procedure for other antibodies. In addition, MVP enabled the exploration and visualization of non-canonical PHP presented by primary mouse tissues and cell lines. Thus, we anticipate that MVP will be widely used by non-experts and experts alike to assess the quality of immunopeptidomic data generated by MS for the analysis of both canonical and non-canonical MHC-associated peptides.

Sample preparation remains a major technical limitation that hampers the accessibility of immunopeptidomics to non-experts (1, 12). In this regard, leaders in the field have argued that significant investments for the development of new reagents, including new monoclonal antibodies, must be prioritized to make the isolation of MHC-associated peptides faster, cheaper, and more robust (1, 12). For instance, antibody engineering and application of robotics and digital microfluidics could remarkably increase the robustness and reproducibility of sample preparation procedures in immunopeptidomics (55–61). Achieving high robustness for the isolation of MHC-associated peptides could be transformative for the field and could drastically accelerate its accessibility, expansion, and maturity (12). Until now, the W6/32 antibody remains the only widely used and commercially available monoclonal antibody in the field. In addition, no protocol for MHC peptide isolation have been standardized across the community (1, 2, 12, 62). Thus, immunopeptidomics is of great scientific and clinical relevance but suffers from a lack of standardized protocols and highly efficient and accessible antibodies for the analysis of MHC class I and II immunopeptidomes in mouse, rat, monkey, and human. Tools such as MVP are instrumental in the development of new reagents, methods, and technologies. We anticipate MVP will contribute to unlocking the full potential of immunopeptidomics in biomedical research (12).

## Supporting information

All supplementary data and tables

## List of abbreviations

ACN: Acetonitrile
BA: Binding Affinity
EL: Elution binding affinity
FA: Formic acid
GUI: Graphic User Interface
HLA: Human Leukocyte Antigen
M: Oxidation
MHC: Major histocompatibility complex
MS: Mass spectrometry
MVP: MhcVizPipe
NQ: Deamidation
PHP: Potential Hybrid Peptides
TFA: Trifluoroacetic Acid

## Acknowledgement

IRIC proteomics facility is a Genomics Technology platform funded in part by the Canadian Government through Genome Canada.

## DATA AVAILABILITY

The mass spectrometry proteomics data have been deposited to the ProteomeXchange Consortium via the PRIDE(63) partner repository with the dataset identifier PXD022195.

## List and legends of Supplemental Data S1 to S26 and Supplemental Tables S1 to S4

### Supplemental Data

**Supplemental data S1-S2. Example of MVP report for MHC class I human dataset.** Complete MVP report of for human MHC class I analysis of PMBC samples from published datasets (DOI: https://doi.org/10.7554/eLife.07661.001, samples PMBC#1 and PBMC#2).

**Supplemental data S3. Example of MVP report for MHC class II human dataset.** Complete MVP report for human MHC class II analysis of JY human cell line from the published dataset (https://doi.org/10.1038/s41587-019-0289-6).

**Supplemental data S4 and S5. Complete MVP report to evaluate the effect of 2 lysis buffer using B-LCL and JY cancer cells.** 100 million of B-LCL (S6) and JY (S7) human cancer cells expressing a high level of MHC class I treated with 0.5 % chaps lysis buffer or NaODEOXY buffer prior to immunoprecipitation with the pan-anti-MHC class I W6/32 antibody using magnetic A beads and 0.2% formic acid for the elution of MHC-ligand complexes (See Supplementary Figure S23a-f). The 2 types of cell lysis buffer generated highly similar results in terms of peptide length, binding affinity scores, and binding motifs.

**Supplemental data S6. Complete MVP report to exemplify the effect of 3 different types of beads for the immunoprecipitation of JY immunopeptidomes.** 100 million of JY human cancer cells treated with 0.5 % chaps lysis buffer prior to immunoprecipitation with the pan-anti-MHC class I W6/32 antibody using 4 different types of beads including magnetic Sepharose protein A beads (MAGA), Sepharose CNBr-activated beads (CNBR), and Sepharose protein G beads (SEPG) and 0.2% formic acid for the elution of MHC-ligand complexes. All types of beads used gave good results in terms of binding affinities and binding motif, although MAGA allowed the identification of a few more peptides.

**Supplemental data S7. Complete MVP report to exemplify the effect of incubation time for the immunoprecipitation of JY immunopeptidomes.** Cell lysates of 100 million of JY human cancer cells were incubated 90 minutes at room temperature or overnight at 4 degrees Celsius prior to immunoprecipitation with the pan-anti-MHC class I W6/32 antibody using CNBr-activated beads and 1% TFA for the elution of MHC-ligand complexes. Both incubation time generated similar results in terms peptide length, binding affinity scores, and binding motifs. Three biological replicates for each condition are shown.

**Supplemental data S8. Complete MVP report to exemplify the effect of 4 different types of beads for the immunoprecipitation of EL4 immunopeptidomes.** 100 million of EL4 mouse cancer cells treated with 0.5 % chaps lysis buffer prior to immunoprecipitation with the anti-MHC class I H2Kb (Y3) antibody using 4 different types of beads including Sepharose CNBr-activated beads (CNBR), Sepharose protein G beads (SEPG), Sepharose protein A beads (SEPA) and magnetic Sepharose protein G beads (MAGG) and 1% TFA for the elution of MHC-ligand complexes. CNBR and SEPG beads were similar in terms of peptides number, peptide length, binding affinities scores, and binding motif and outperformed SEPA and MAGG beads.

**Supplemental data S9. Complete MVP report to exemplify the effect MHC-ligand complexe elution buffers for the immunoprecipitation of EL4 immunopeptidomes.** 100 million of EL4 mouse cancer cells treated with 0.5 % chaps lysis buffer prior to immunoprecipitation with the anti-MHC class I H2Kb (Y3) antibody using sepharose protein G beads and 4 different acidic elution buffers including 0.2% formic acid, 10% acetic acid, 1% TFA and 10% TFA. The four acidic elution buffers led to the isolation of peptides sharing the same expected H2Kb binding motif but 1% TFA elution buffer led to the identification of the highest number of strong H2Kb binder peptides.

**Supplemental data S10. Complete MVP report to exemplify the effect MHC-ligand complexes elution buffers for the immunoprecipitation of A20 immunopeptidomes.** 100 million of A20 mouse cancer cells treated with 0.5 % chaps lysis buffer prior to immunoprecipitation with the anti-MHC class II IA-IE (M5) antibody using sepharose protein G beads and 4 different acidic elution buffers including 0.2% formic acid, 10% acetic acid, 1% TFA and 10% TFA. 1% TFA elution buffer led to the identification of the highest number of peptides with the expected binding motif.

**Supplemental data S11. Complete MVP report of canonical H2D^b^ peptides from 19 mouse tissues.** Analysis was performed using data published from https://doi.org/10.7554/eLife.07661.001. Refer to Table S3 for peptide the list.

**Supplemental data S12. Complete MVP report of PHP H2D^b^ from 19 mouse tissues.** Analysis was performed using data published from https://doi.org/10.7554/eLife.07661.001 and reprocessed with the Hybrid Finder pipeline to identify non-canonical peptides. Refer to the Supplemental Table S4 for the peptide lists.

**Supplemental data S13. Complete MVP report of canonical H2K^b^ peptides from 19 mouse tissues.** Analysis was performed using data published from https://doi.org/10.7554/eLife.07661.001. Refer to the Supplemental Table S2 for the peptide lists.

**Supplemental data S14. Complete MVP report of PHP H2K^b^ from 19 mouse tissues.** Analysis was performed using data published from https://doi.org/10.7554/eLife.07661.001 and reprocessed with the Hybrid Finder pipeline to identify non-canonical peptides. Refer to the Supplemental Table S4 for the peptide lists.

**Supplemental data S15. Complete MVP report of canonical H2D^b^ peptides from mouse cancer cell lines.** Analysis was performed using data published from https://doi.org/10.7554/eLife.07661.001.

**Supplemental data S16. Complete MVP report of PHP H2D^b^ peptides from mouse cancer cell lines.** Analysis was performed using data published from https://doi.org/10.7554/eLife.07661.001.

**Supplemental data S17. Complete MVP report of canonical H2K^b^ peptides from mouse cancer cell lines.** Analysis was performed using data published from https://doi.org/10.7554/eLife.07661.001.

**Supplemental data S18. Complete MVP report of PHP H2K^b^ peptides from mouse cancer cell lines.** Analysis was performed using data published from https://doi.org/10.7554/eLife.07661.001.

**Supplemental data S19. Comparison of EL and BA scoring from NetMHCpan4.0 for class I peptides.**

**Supplemental data S20. Comparison of EL and BA scoring from NetMHCIIpan4.0 for class II peptides.**

**Supplemental Data S21. Binding of H2K^b^ antibody (Y3) on different types of beads**.

**Supplemental data S22a.** General comments for the isolation of MHC class I and II immunopeptidomes.

**Supplemental data S22b.** Quantification of the mean number of MHC molecules/cell using the QiFiKit.

**Supplemental data S23a.** General comments for beads coupling with the antibodies

**Supplemental data S23b.** Detailed protocol for coupling CNBr activated beads

**Supplementary data S23c.** Coomassie gel staining to track covalent binding efficiency of anti-HLA A, B, C (W6/32) to Sepharose CNBr-activated beads.

**Supplementary data S23d.** Detailed protocol for coupling Sepharose beads A or G

**Supplementary data S23e.** Coomassie gel staining to track binding efficiency of anti-HLA A, B, C (W6/32) to Sepharose protein G beads.

**Supplemental data S23f.** Detailed protocol for coupling Magnetic Sepharose beads A or G with W6/32 antibody

**Supplemental data S23g.** Detailed protocol for covalent binding of antibody to Magnetic Sepharose beads or Protein A or G Sepharose beads.

**Supplemental data S24a.** General comments for detailed protocols for immunoprecipitation of MHC Class I and II ligands

**Supplemental Figure S24b.** Detailed protocol for immunoprecipitation of MHC Class I and II ligands using Sepharose CNBr-activated or Sepharose A/G protein beads and Y3, W6, L243, and M5 antibodies

**Supplemental data S24c**. BCA quantification of proteins from cell lysate isolated from 100 million cells using 0.5% chaps buffer

**Supplemental data S24d.** Western blotting for tracking MHC-ligand complexes acidic elution from Sepharose protein G beads.

**Supplemental data S24e.** Detailed protocol for immunoprecipitation of MHC Class I using Magnetic Sepharose A/G protein beads

**Supplemental data S24f.** Western blotting for tracking MHC-ligand complexes acidic elution from magnetic protein A beads.

**Supplemental data S24g.** Visual of the Bio-Rad and C18 columns set up for the elution of MHC-peptides complexes peptides.

**Supplemental data S25.** Supplemental methods for the creation of the MVP

**Supplemental data S26.** MVP installation on Windows 10 using a Linux Subsystem WSL 1

### Supplemental tables

**Supplemental Table S1**: All datasets used in the paper

**Supplemental Table S2:** H2D^b^ Canonical peptide lists of 19 primary mouse tissues

**Supplemental Table S3:** H2K^b^ Canonical peptides lists of 19 primary mouse tissues

**Supplemental Table S4:** H2D^b^ and H2K^b^ PHP peptide lists of 19 primary mouse tissues

**Supplemental data S19.**
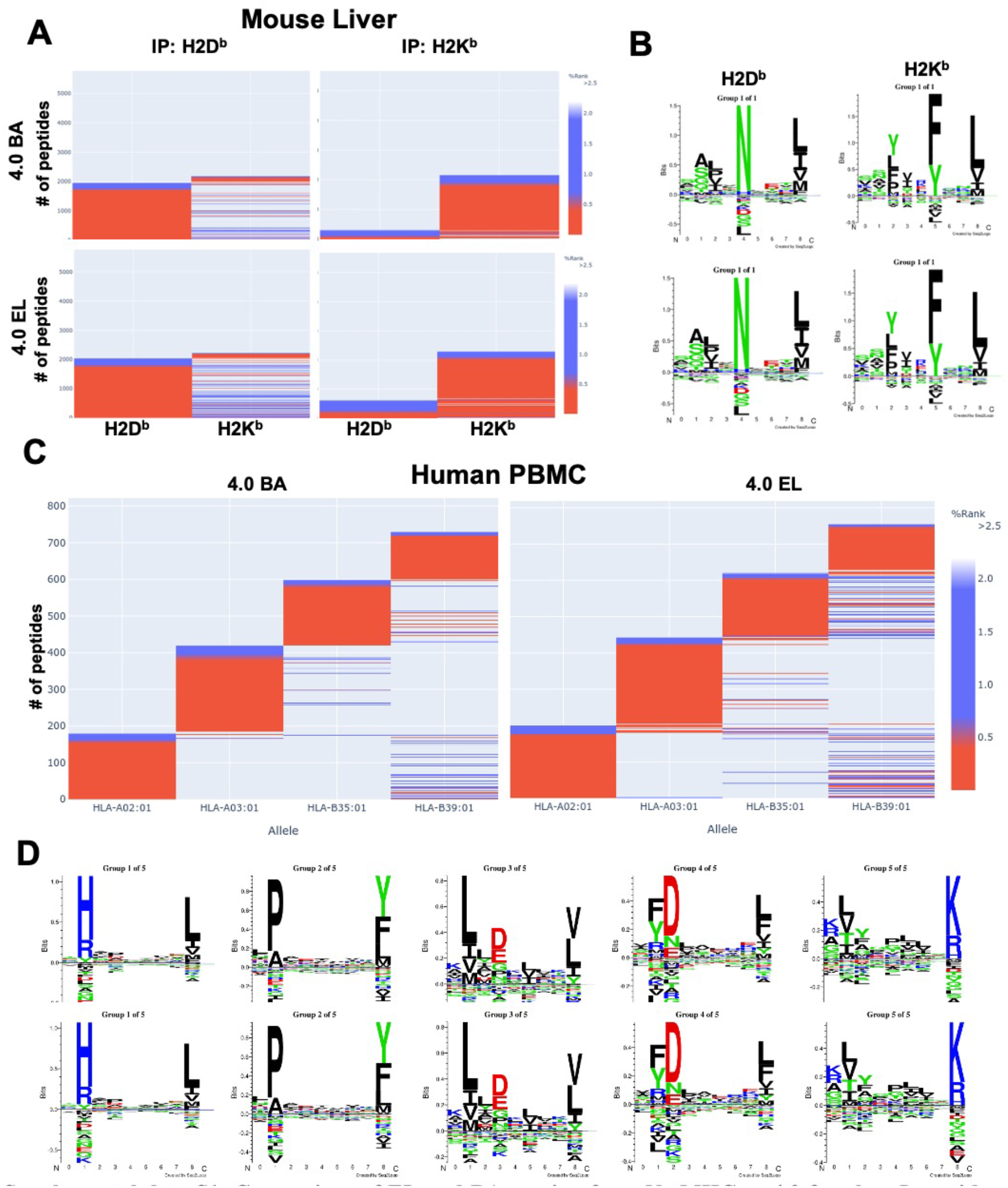
Comparison of EL and BA scoring from NetMHCpan4.0 for class I peptides. Heatmaps and sequence motifs of mouse liver (A & B) and human PBMC (C & D) samples show insignificant differences between the two scoring systems for class I peptides. Mouse data were used from https://doi.org/10.1038/sdata.2018.157 and human data were used from https://doi.org/10.7554/eLife.07661.001.

**Supplemental data S20.**
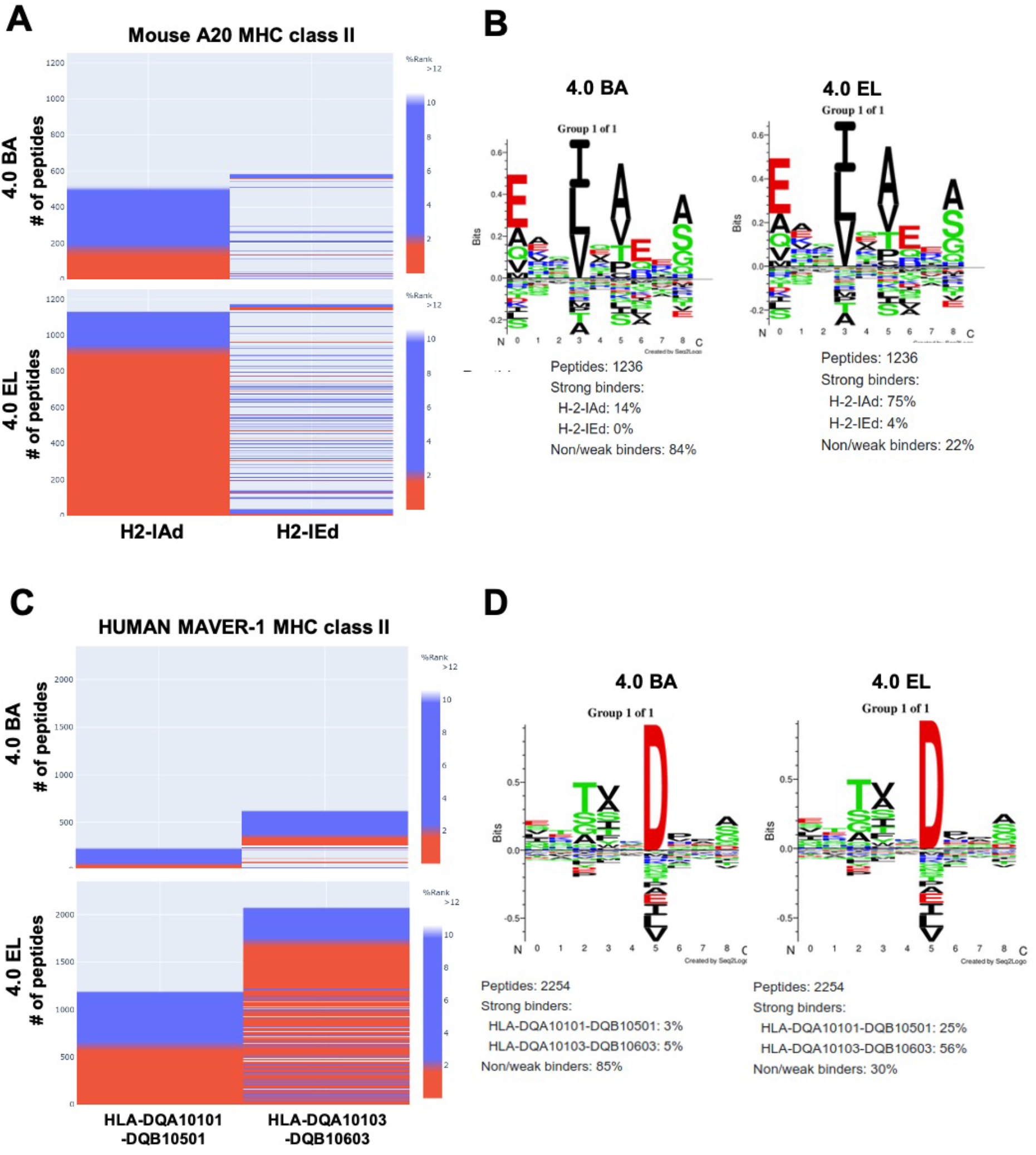
Comparison of EL and BA scoring from NetMHCIIpan4.0 for class II peptides. Heatmaps and sequence motifs mouse A20 cell line (A-B) and human MAVER-1 cell line (C-D) samples show a striking difference between the two scoring systems for class II peptides. The EL score increases the number of binders by approximately 4-fold over BA scoring. Published mouse data were used from https://doi.org/10.1002/eji.201545930 and human data were used from https://doi.org/10.1002/pmic.201700246.

#### Explanation for Supplemental data S19 and S20

In developing MhcVizPipe we had to decide whether to use NetMHC(II)pan’s eluted ligand (EL) or binding affinity (BA) scores. BA is a score reflecting the likelihood of observing a given peptide in a binding affinity study. EL is a score reflecting the likelihood of observing a given peptide in an eluted ligand experiment. A BA study directly measures the affinity between a MHC molecule and peptides. The advantage a BA study is its high specificity as experiments can be performed with purified MHC molecules. However, BA studies only measure binding affinity, a single part of the biological presentation of MHC-associated peptides (MAPs). An EL experiment is any experiment in which MHC-associated peptides are eluted from the cell surface and subsequently purified and measured qualitatively or quantitatively (e.g. by mass spectrometry). Examples of EL experiments are immunopurification and weak acid elution. The advantage of EL studies is that they directly represent the MAPs present on the cell surface, and thus incorporate the entire biological process of antigen presentation. The disadvantage is that it is difficult to be highly specific in these experiments because cells generally express more than one MHC allele and there is invariably contamination from non-MAP peptides. Because the advantages and disadvantages of BA and EL are inverse of each other, NetMHCpan and NetMHCIIpan incorporate data from both BA and EL experiments in their training sets. Though the scores are specific to the likelihood of observing a peptide in the respective type of experiment, both algorithms are trained on data from both BA and EL studies.

In our development of MhcVizPipe, we compared EL and BA scoring for both class I and class II peptides. It was expected that the EL scoring system would outperform BA as EL most closely represents the data being analyzed. Furthermore, EL scoring/rank have been recently shown to be more accurate, especially for MHC class II peptides annotation (Garde *et al.* Immunogenetics volume 71, pages445-454; 2019). Nevertheless, we analyzed a number of datasets using both systems as confirmation. As an example, **Figure S19** and **S20** show BA- and EL-identified binders for class I and class II peptides, respectively. For class I peptides (**Figure S19**), it is apparent that the choice of scoring does not significantly impact the allele-annotation of peptides. Both BA and EL results in very similar numbers of weak and strong binders for both mouse (**Figure S19A**) and human (**Figure S19C**) samples, with slightly higher numbers being observed for EL. For class II peptides, a very significant difference is observed for both mouse (**Figure S20A**) and human (**Figure S20C**) peptides. Using EL results in a nearly four-fold increase in annotated peptides compared to BA. Furthermore, the peptide groups identified by GibbsCluster (**Figures S20B & S20D**) contain significantly more EL-identified strong binders (**Figure S20D**) than BA-identified (**Figure S20B**). These findings in regards to class II peptides are in agreement with the conclusions of Garde *et al*. MHC class II peptide annotation has always been more challenging as compared to MHC class I peptides. This could be due to the different biology/nature of these peptides. Indeed, MHC class II peptides tend to be longer with intermittent conformation (Sadegh-Nasseri et al. Immunol Res 2010; 47:56-64) and the primary sequences of the peptides might not be sufficient to predict binding (like with BA). It is also possible that the tertiary/quaternary structure formed by the folding of the peptides within the MHC groove dominates the binding prediction. The increased performance obtained when using EL scoring, especially in relation to class II peptides, led us to choose it as the scoring system used by MhcVizPipe.

#### Supplemental Data S21

**Supplemental data S21.**
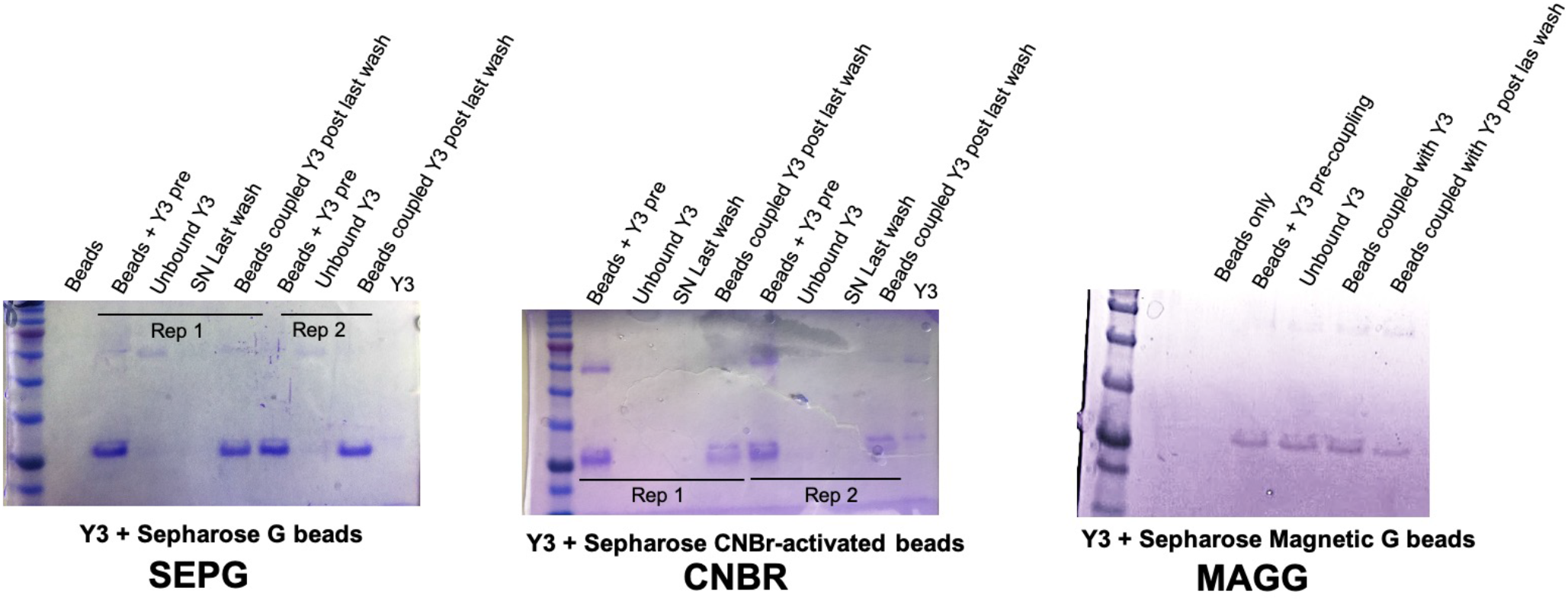
Binding of H2K^b^ antibody (Y3) on different types of beads. Coomassie gel staining to track binding efficiency of anti-H2K^b^ (Y3) to Sepharose protein G beads (SEPG) (left panel), Sepharose CBNR-activated beads (CNBR) (middle panel) and magnetic Sepharose G beads (MAGG) (right panel) reveals a poor binding of Y3 antibody on MAGG beads. Equivalent volumes of Beads only, Beads + Y3 pre-coupling, unbound antibody following coupling step, supernatant following last wash after coupling, aliquots were loaded onto 12% SDS-PAGE gel followed by Coomassie blue staining. Two replicates (Rep1 and Rep2) are shown Sepharose Protein G and CNBR-activated beads. A similar amount of staining between the beads + antibody pre-coupling vs the beads coupled with Y3 post last wash aliquots is indicative a successful coupling. Note that when the antibody is covalently bound to the CNBr beads specifically, the staining of the antibody on the Coomassie gel should be less intense as compared to the initial starting material containing the unbound antibody and the beads.

#### S22a. General comments regarding the isolation of MHC class I and II immunopeptidomes

##### What’s new with these protocols

The protocols we described here were adapted from established protocols^1–6^ and we additionally optimized them for the isolation of mouse immunopeptidomes since those for human are easier and more widespread in the community. Note that the optimized protocols for mouse samples work also very well for human samples. These protocols are intended to be ready-to-use in any laboratory and don’t require the additional step of purification of eluted peptides by fractionation, lyophilization and/or by ultrafiltration using 10-30 kDa filters as described by others^1–6^. However, if needed, this step could be considered among other factors to optimize the overall yield and numbers of peptide identified. On one hand, these protocols do not cover the isolation of antibodies from supernatant of hybridoma or the peptide acquisition by MS/MS and peptide identification. On the other hand, general comments and detailed step-by-step procedures—including steps to verify the beads-binding efficiency of the antibody and the elution efficiency of the MHC-ligand complexes from the beads—are described. In the protocols provided here, different steps in which aliquots can be taken for making quality control gels are indicated. An example of Coomassie blue staining gels as well as Western blots are provided here.

##### HLA/MHC genotyping and expression levels

The MHC class I and II subtype must be known for the successful isolation of class I and II ligands and can be done by high resolution HLA typing^7^. For various cell lines and mice, the documented HLA subtype can be found at https://web.expasy.org/cellosaurus. The level of HLA proteins expressed on the cell surface is also an important factor to consider. Evidently, low expression of these molecules hampers the number of peptides identified. The number of MHC molecules expressed on the cell surface can be quantified by flow cytometry using standard beads as described in the commercially available kit QIFIKIT (K007811-8, Dako Japan, Kyoto, Japan). For example, quantification of MHC class I H2K^b^ and H2D^b^ molecules for EL4 and HLA class I A, B and C molecules for JY cells are shown in data S22b. These cell lines are considered to express high levels of these molecules and are good positive controls for IP.

**Supplementary data S22b.**
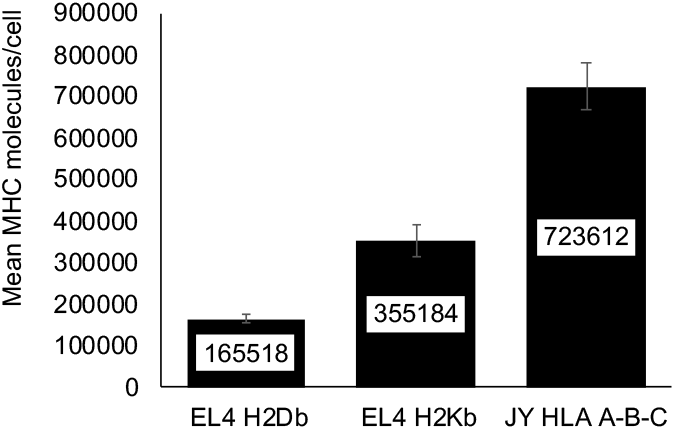
Quantification of the mean number of MHC molecules/cell using the QiFiKit. Mean number of MHC class I H2Db and H2Kb quantified from one million of EL4 cells of HLA A, B, C molecules from one million of JY cells. Mean and standard error of the mean are represented from 3 independent experiments.

##### Cell culture

We culture EL4, JY and A20 cells to generate pellets of 100 million cells as described below. JY, EL4 and A20 cell lines were cultured in T75 flat bottom flasks and kept straight in the incubator in a 25 ml of media. We observed that using bigger flasks (i.e. T175) reduced cell growth and cell viability leading to lower number of peptides identified. In order to get reproducible results, cells were used at low passage and cultured using an organized schedule (see below). Cells are always plated at the same cell density at day 1 and cell viability is constantly monitored. Cell viability is an important factor that could account for variability between samples and we strongly encourage to measure this factor when harvesting the cells. EL4, JY and A20 usually display 90% of cell viability at the moment of harvesting.

##### Complete procedure for harvesting 100 million cells pellets

All cell lines were purchased from ATCC. On day one of the cell culture schedule, the lymphoblastoid mouse cell line EL4 was seeded in two T75 cm^2^ Canted Neck Blue Vented Cap Tissue Culture Treated flasks (Ref number: 353136, Falcon) with 10×10^6^ cells in a volume of 25 ml (0,4×10^6^ cells/ml) of X-vivo 15 media (X-vivo 15 (04-744Q, Lonza) supplemented with 25% DMEM (319-005-CL, Multicell), 10% FBS (98150, Multicell) and 1% pen/strep (450-201-EL, Multicell). When using the human lymphoblastoid B-cell line JY, two T75 cm^2^ Canted Neck Blue Vented Cap Tissue Culture Treated flasks were seeded with 18.75×10^6^ cells in a volume of 25 ml (0,75×10^6^ cells/ml) of DMEM media (DMEM supplemented with 10% FBS and 1% pen/strep). The lymphoblastoid B-cell line A20 was seeded in two T75 cm^2^ Canted Neck Blue Vented Cap Tissue Culture Treated flask flasks with 7.5×10^6^ cells in a volume of 25 ml (0,3×10^6^ cells/ml) of RPMI 1640 1X (350-000-CL, Multicell) supplemented with 10% FBS, 1% pen/strep, 1% sodium pyruvate 100mM (100X)(600-110-EL, Multicell), 1% Hepes 1M solution (330-050-EL), 1% D-glucose (0188-500G, Amresco) and 0,1% Beta-mercaptoethanol (190242, MP Biomedicals). All cell lines were incubated at 37C° with 5% CO_2_ for a period of 48 hours, until they reached a concentration of approximately 2×10^6^ cells/ml. All three cell lines were harvested following the same protocol: 100×10^6^ cells were harvested and centrifuged at 433 rcf (1500 RPM, Swinging bucket rotor A-4-81, Eppendorf centrifuge 5810R) for a period of 5 minutes at room temperature in 50 ml polypropylene conical sterile centrifuge tubes (Ref: 352070, Falcon) The culture medium was removed by aspiration and the cell pellets were washed gently by pipetting up and down (without vortexing) with 5 ml of PBS (catalog number:311-010-CL). After a second round of centrifugation with the same parameters, PBS was removed and the cell pellets were transferred in a 15 ml high-clarity polypropylene conical sterile centrifuge tubes (Ref: 352096, Falcon) with 3ml of PBS. The tubes were centrifuged at 433 rcf (1500 RPM, Swinging bucket rotor A-4-81, Eppendorf centrifuge 5810R) for 5 minutes. After removing the PBS by aspiration, the cell pellets were stored at −80 degrees Celsius.

##### Cell pellets storage

In our hands, we observed that the fresher the cell pellet was, the more peptides we identified. The quality of cell pellets stored more than 6 months at −80 degrees Celsius was dramatically lowered.

##### Technical positive control

As mentioned in the manuscript, most of the published protocols refer to the isolation of immunopeptidomes from human tissues or cells and the protocols were optimized for the W6/32 antibody or for in house purified antibodies. Since the isolation of the immunopeptidomes require multiple steps, we strongly encourage the use of a technical positive control such as JY or B-LCL human cancer cells due to their high expression levels of MHC class I molecule. This cell line-antibody combinations work very well with most conditions/beads/elution buffers and up to 5000 peptides can easily be identified from 100 million cells. JY cells could also be used as a technical control for MHC Class II HLA-DR peptides using the L243 antibody from BioXcell. Up to 1000 peptides can be identified using this combination of cell line-antibody. It is possible to make a stock of JY or B-LCL cell pellets and keep them at −80 degrees Celsius and use them as a positive control for each run IP.

##### Timeline for the isolation of immunopeptidomes by IP

10-20 samples per experiment can be processed in parallel. One trick is to label all the required tubes (up to 9 tubes per sample) the day before the experiment.

Day 1:

- Prepare fresh lysis buffer and solutions for beads coupling
- Coupling the beads with the antibody
- Performing Coomassie blue gel to evaluate binding efficiency

Day 2

- Cell lysis of the cells/tissue to be analyzed
- Incubating cell lysate with the coupled beads overnight for 3 hours at 4 degrees Celsius

Day 2 (for 3 hours incubation) or Day 3 (for overnight incubation) *:

- Prepare fresh washing and acidic elution buffers
- Collecting and washing the beads with bounded MHC-ligand complexes
- Elute MHC-ligand complexes
- C18 cleanup and elution of peptides from C18 column
- Storage of the isolated peptides

Unspecific required time:

- Defining the HLA subtype of the cells of interest
- Evaluation of levels of MHC class I and II expression using, for example, the commercial kit QIFIKIT
- Generation of 100 million cells pellets by cell culture
- Data acquisition by MS/MS and peptide identification

*Note: It is possible to perform all the steps of Day 2 and Day 3 the same day if the incubation of the antibody-coupled beads with the cell lysate is done for 3 hours at 4 degrees Celsius.

#### S23a: General comments regarding beads coupling with the antibodies

##### Where to start

Sepharose CNBr-activated beads are a generally good starting point as we observed good performance with 4 commercially available antibodies (Y3, W6/32, L243 and M5). Besides their affordable price, the Sepharose CNBr-activated beads offer the advantage of covalent binding of the antibody to the beads and require only 4 hours of preparation. Protein A or G or A/G Sepharose 4 Fast Flow beads are also very easy to handle and affordable and they generate similar results as the Sepharose CNBr-activated beads. Here, the antibody is not covalently bound to the beads and therefore an additional crosslinking step with DMP is needed if the beads need to be re-used. In our hands, these 2 types of sepharose beads were binding efficiently to the anti-mouse MHC class I H2K^b^ (Y3), anti-Mouse MHC Class II IA/IE (M5/114), anti-Human MHC class I HLA A, B, C (W6/32) and anti-Human MHC class II HLA-DR (L243) antibodies. Of note, the Sepharose magnetic beads are also very easy to use but they remain more expensive and, in our hands, these beads are working very well only with the W6/32 antibody (refer to Supplementary Figure S10a-b).

The following factors could be evaluated to optimize the beads coupling procedure:

- The type of beads used to couple the antibody (we’ve tested Sepharose 4 fast flow A or G or A/G, Sepharose CNBr-activated beads, Magnetic Sepharose A or G or A/G beads)
- Amount of antibody and/or of the beads per IP – starting with 1 mg of antibody/IP
- Cross-linking or not the antibody to the beads

Here, we provide 3 optimized protocols for beads coupling:

- Supplemental data S22b-c: Sepharose CNBr-activated beads
- Supplemental data S22d-e: Sepharose protein A or G or A/G beads
- Supplemental data S22f: Sepharose protein A or G or A/G magnetic beads

The antibodies commercially available from Bioxcell (Cederlane) tested with these protocols are:

- Mouse M5/114 (BE0108) ‒ Anti-Mouse MHC Class II (IA/IE)
- Mouse Y3 (BE0172) ‒ Anti-Mouse MHC class I (H2K^b^)
- Mouse M1(BE077) ‒ Anti-Mouse MHC class I (H2K^b^ and H2D^b^)
- Human W6/32 (BE0079) ‒Anti-Human MHC class I (HLA A, B, C)
- Human L243 (BE0306) ‒ Anti-Human MHC class II (HLA-DR)

#### S23b – Detailed protocol for coupling Sepharose CNBr-activated beads

Note: Protocol adapted from Nelde al^1^.

##### Material

- Micro tube, 2 ml Protein LoBind (cat # 022431002, Eppendorf)
- CNBr activated sepharose 4B from GE Healthcare (Cat# 17-0430-01))
- Antibody of your choice (2 mg/sample). Note that 1 mg of antibody per sample could be enough for some antibodies. Some antibodies bind better than other and require smaller amount for beads coupling like the W6/32 antibody. In general, we use 80 mg of beads/2 mg antibody (sample).

##### Solutions

**Reactivating solution for Sepharose CNBr-activated Sepharose**: 1 mM HCl.

- 20ul HCl 6N in 120 ml H2O

**Antibody coupling buffer:** 0.5 M NaCl, 0.1 M NaHCO_3_, pH 8.3.

- 2.92 g NaCl
- 0.84 g NaHCO_3_
- Add 100 ml of water
- Mix and adjust pH with drops of NaOH 10 N

**Blocking solution for Sepharose CNBr-activated:** 0.2 M glycine.

- 0.15 g glycine
- Add 10 ml of H2O

###### Procedure for 1 sample

The following steps are performed at room temperature.

1. <u>Beads activation</u>

- Label 1× 15 ml tube
- Weight 0.04g (40 mg) of beads to each 15 ml tube
- Add 13.5 ml of 1 mM HCl
- Rotate slowly on a rotating wheel for 30 minutes
2. <u>Coupling: R.T.</u>

- Prepare tubes during the activation step. For example:

○ Tube 1 = Y3 = 307 ul Y3 + 193 ul coupling buffer
○ Tube 2 = M1 = 230 ul M1 + 270 ul coupling buffer
○ Tube 3 = M5 = 285 ul = 215 ul coupling buffer
○ Tube 4 = W6 = 294 ul W6 + 206 ul coupling buffer
○ Tube 5 = L243 = 247 ul of L243 + 243 ul of coupling buffer
- Spin down sepharose at 1000 RPM with for 2 minutes
- Remove SN
- Add 0.5 ml of coupling buffer to each tube
- Transfer the beads into tubes containing the antibody
- Slowly rotate for 120 min.
3. <u>Blocking</u>

- Spin beads at 113 rcf (1000 RPM, Fixed-angle rotor FA-45-48-11, Eppendorf centrifuge 5430R) for 3 minutes.
- OPTIONAL: Take a 18 ul aliquot of the supernatant (Unbound antibody)
- Aspirate supernatant
- Add 1.0 ml of 0.2 M glycine
- Rotate slowly on a rotating wheel for 60 min (22 RPM).
4. <u>Washing</u>

- Spin beads at 113 rcf (1000 RPM, Fixed-angle rotor FA-45-48-11, Eppendorf centrifuge 5430R) for 3 minutes
- Aspirate supernatant
- Add 1.0 ml PBS
- Spin
- Aspirate supernatant
- Repeat washing step
- OPTIONAL: Take a 18 ul aliquot of the supernatant LAST WASH
- Resuspend in 1 ml PBS and transfer in a new tube.
- OPTIONAL: Take a 18 ul Aliquot BEADS COUPLED
- Store the beads and aliquots at 4 degrees Celsius until use.

#### Tracking efficiency of beads coupling using Coomassie blue staining SDS-PAGE gels

In order to evaluate the coupling efficiency of the antibodies to the beads, aliquots are collected at key steps of the protocol. Note that when the antibody is covalently bound to the beads, the staining of the antibody should be less intense as compared to the initial starting material containing the unbound antibody and the beads. See S22c for a successful coupling.

**Supplementary Data S23c.**
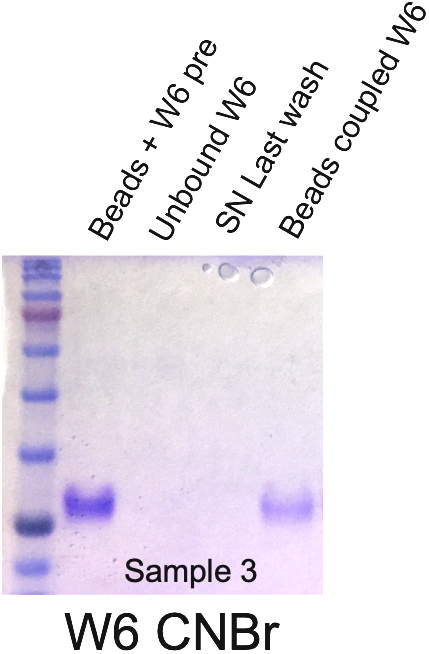
Coomassie gel staining to track covalent binding efficiency of anti-HLA A,B,C (W6/32) to Sepharose CNBr-activated beads. Equivalent volumes of aliquots from beads + uncoupled antibody pre-incubation (Lane 1), unbound antibody following coupling step (Lane 2), supernatant following last wash after coupling (Lane 3) and antibody covalently linked with the beads (Lane 4) were loaded onto 12% SDS-PAGE gel followed by Coomassie blue staining. These gels are representative of n = 5 independent experiments.

#### S23d – Detailed protocol for coupling Proteins A or G Sepharose beads

##### Material

- Micro tube, 2 ml Protein LoBind (cat # 022431002, Eppendorf)
- Protein G sepharose 4 fast flow – (cat # 17-0618-01, GE Healthcare)
- Protein A Sepharose 4B conjugate (cat # 10142, Invitrogen)
- Antibody of your choice (2 mg/sample). Note that 1 mg of antibody per sample could be enough for some antibodies. Some antibodies like the W6/32 antibody bind better than other and require smaller amount for beads coupling.

##### Procedure for 1 sample

Bring beads to room temperature before use

1. Vortex the vial containing beads to resuspend the slurry solution
2. Pipet 0.5 ml of slurry solution and transfer into 2.0 ml tube
3. Add 1 ml PBS/vortex
4. Spin beads at 113 rcf (1000 RPM, Fixed-angle rotor FA-45-48-11, Eppendorf centrifuge 5430R) for 3 minutes.
5. Aspirate supernatant
6. Add 1 ml PBS/vortex
7. OPTIONAL: Take a 18ul aliquot (beads)
8. Spin
9. Aspirate supernatant
10. Add 2.0 mg of antibody and fill to 1ml with PBS
11. Vortex
12. OPTIONAL: Take a 18ul aliquot (Beads + antibody pre-incubation)
13. Incubate at room temperature for 3 hours using slow rotating wheel (RPM 22)
14. Spin
15. OPTIONAL: Take a 18ul aliquot of the supernatant (unbound antibody)
16. Aspirate supernatant
17. Add 1 ml of PBS/vortex
18. Spin
19. OPTIONAL: Take a 18ul aliquot of the supernatant (last wash)
20. Aspirate supernatant
21. Add 1 ml of PBS and store at 4 degrees Celsius
22. OPTIONAL: Take a 18ul aliquot (beads coupled with antibody)
23. Store the beads at 4 degrees Celsius

Note: Coupled-beads and aliquots can be stored at 4 degrees Celsius until use.

#### Tracking efficiency of beads coupling using Coomassie blue staining SDS-PAGE gels

In order to evaluate the binding efficiency of antibodies coupled to the beads, aliquots are collected at key steps of the protocol. As shown in supplemental data S22e, efficient antibody binding with beads is achieved when minimal amount of unbound antibody is found in the supernatant following coupling and when the staining of the beads coupled with the antibody is almost similar to the starting material (Beads + antibody).

**Supplementary Data S23e.**
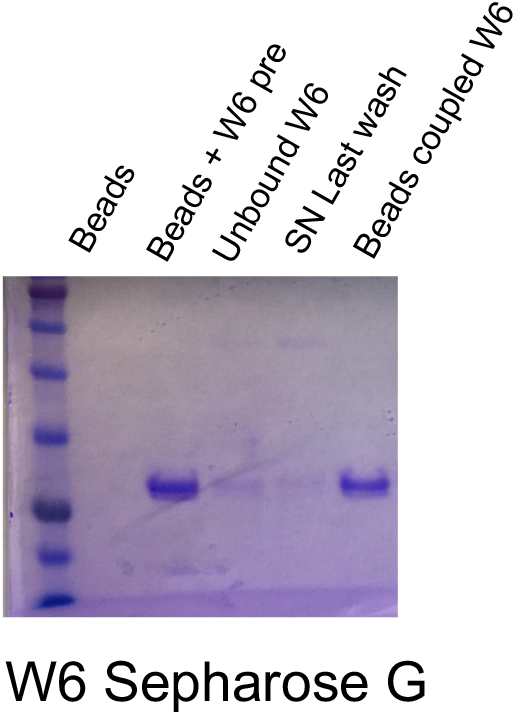
Coomassie gel staining to track binding efficiency of anti-HLA A,B,C (W6/32) to Sepharose protein G beads. Equivalent volumes of aliquots from beads only (Lane 1), from beads + uncoupled antibody pre-incubation (Lane 2), unbound antibody following coupling step (Lane 3), supernatant following last wash after coupling (Lane 4) and antibody linked with the beads (Lane 5) were loaded onto 12% SDS-PAGE gel followed by Coomassie blue staining. These gels are representative of n = 5 independent experiments.

#### S23f. Detailed protocol for coupling Magnetic Sepharose beads A or G with W6/32 antibody

##### Material

- Pure proteome Protein A or G magnetic beads (Cat # LSKMAGA10, LSKMAGG10, EMD Millipore)
- 1 mg of antibody per sample
- Note: It is possible to prepare a batch of magnetic beads.

##### Coupling of W6/32 antibody to Protein A magnetic beads

Preparation for 1 sample

1. Gently resuspend the magnetic bead slurry
2. Pipet 1 ml of resuspended magnetic bead slurry into 2 ml tube
3. Place tube on magnet and remove liquid
4. Add 1 ml of PBS
5. Place tubes of magnet and remove liquid
6. Repeat washes twice
7. Remove PBS
8. Add 1 mg of antibody
9. Add 1 ml of PBS
10. Incubate at RT for 60 minutes with rotating wheel (22 RPM)
11. Place tubes of magnet and remove liquid
12. Repeat washes twice (2 × 1 ml PBS)
13. Keep beads in 1 ml PBS and store at 4 degrees Celsius until use.

#### S23g. Detailed protocol for covalent coupling of antibody to Sepharose magnetic beads or Protein A or G Sepharose beads

Note: This step is optional and can be applied to non-magnetic sepharose beads also.

##### Material

- Triethanolamine (Cat# 90279-100 ml, Sigma)
- Dimethyl pimelimidate-2HCl (Cat# 21667, Pierce)

##### Solutions

200 mM triethanolamine pH 8.3

- 7.5 g of triethanolamine (Tare your glass bottle and add 7.5 g of Triethanolamine viscous solution)
- Add 150 ml water
- Adjust pH at 8.3 with HCl
- Fill up to 250 ml water

5mM DMP

- 143 mg DMP + 110 ml 200 mM triethanolamine pH 8.3

##### Procedure

1. Wash beads with 3 × 1 ml of 200 mM triethanolamine pH 8.3

a. Magnetic beads: place on magnet holder, remove liquid, add 1 ml of liquid, aspirate liquid
b. Sepharose beads: aspirate PBS, add 1 ml of liquid, spin at 113 rcf (1000 RPM), aspirate liquid
2. Add 1 ml of 5 mM DMP solution to the tube
3. Resuspend beads and transfer beads into 15 ml Falcon tube
4. Add 10 ml of 5 mM DMP solution into the 15 ml Falcon tubes
5. Incubate at RT for 60 minutes with slow rotation (22 RPM)
6. Add 1 × 550 ul of 1 M Tris-HCl pH7.6
7. Incubate at RT for 30 minutes with slow rotation (22 RPM)
8. Spin at 1000 rpm (200g) for 1 minute at 4^0^C
9. Remove supernatant
10. Add 1 ml of PBS
11. Transfer to 2 ml microfuge tubes

a. Magnetic beads:

i. Place tube of magnet and remove liquid, add 1 ml PBS
ii. Repeat washes twice (2 × 1 ml PBS)
b. Sepharose beads:

i. aspirate PBS, add 1 ml of liquid, spin at 113 rcf (1000 RPM), aspirate liquid
ii. Repeat washes twice (2 × 1 ml PBS)
12. Add 0.9 ml PBS to each tube
13. Store antibody-coupled beads 4 degrees Celsius until use.

#### S24a: General comments for detailed protocols for immunopurification of MHC Class I and II ligands

##### Where to start

When testing a new antibody, we suggest to start with the protocol described in S22-S23, i.e. with 100 million cells lysed with 0.5% chaps buffer, Sepharose CNBr-activated beads, 1% TFA acid elution buffer for eluting the MHC-peptides complexes from the beads and 28% ACN/0.1% TFA for peptide elution from the C18 columns. In our hands, this combination was the most effective in terms of number of peptides identified for the 4 antibodies and cell lines tested. All the possible variables (lysis buffers, type of beads, elution buffers) are described in the detailed protocols.

We observed that the following factors could impact the success of your peptide isolation by immunopurification:

- **Amount of starting material\***
- **Type of lysis buffer\*\***
- **The type of beads used to couple the antibody**
- Amount of antibody
- Cross-linking or not the antibody to the beads
- Time of incubation and incubation temperature of the cell lysate with the coupled-beads
- **Type of elution buffer used to eluate the MHC-peptides complexes from the beads**
- Type of elution buffer used to elute peptides from the C18 column.
- The use of low retention binding tips and HPLC MS/MS grade for water and solvents
- **Freshness of the prepared solutions**

Factors in bold represent those that -in our hands-account for big differences depending on the antibody used.

##### */**Amount of starting material and cell lysis

As shown in supplementary data S3-S4, the cell lysis buffer did not strongly impact the number of peptides identified by MS/MS for JY or B-LCL cells. However, it might have an impact on other cell lines/tissues and it could be considered during protocol optimization (especially for tissues). For instance, the OVCAR3 cell line cannot be solubilized using the chaps buffer and the NaDEOXY buffer is indicated. We estimated that 100 million of cells correspond to approximatively 6-10 mg of proteins/ when using 0.5% chaps lysis buffer (Supplementary data 23b).

**Supplementary data S24b.**
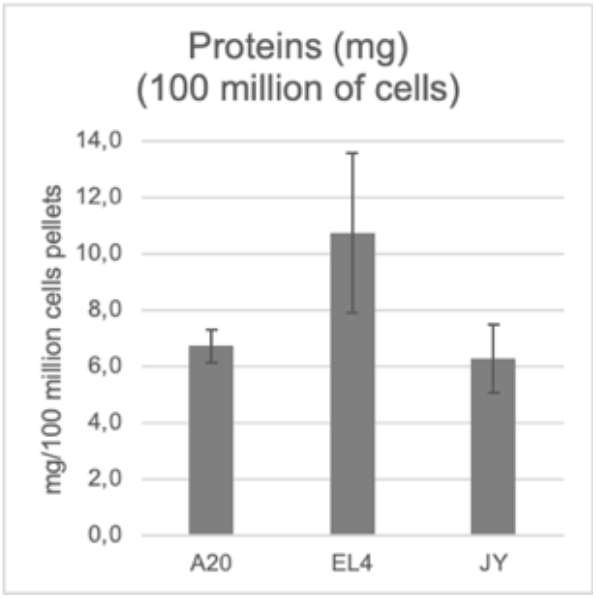
BCA quantification of proteins from cell lysate isolated from 100 million cells using 0.5% chaps buffer. Supernatants of cell lysates were quantified by BCA, n=3 independent experiments.

Here, we present 2 optimized protocols for immunopurification of MHC class I and II ligands from mouse and human cells:

- Supplemental data S23c-d: IP using Sepharose CNBr-activated and Sepharose 4 Fast Flow Protein G beads (or Protein A Sepharose 4B conjugate beads)
- Supplemental data S23e-f: IP using Sepharose protein A or G or A/G magnetic beads

#### S24c: Detailed protocol for immunopurification of MHC Class I and II ligands using Sepharose CNBr-activated or Sepharose A or G protein beads and Y3, W6, L243, M1 and M5 antibodies

##### Material

- 100 million cells
- Sepharose CNBR-activated beads or Protein G/A sepharose 4 fast flow coupled with your antibody of interest (see protocol S22-S23)
- Micro tube, 1.5 ml Protein LoBind (cat # 022431081, Eppendorf)
- Micro tube, 2 ml Protein LoBind (cat # 022431002, Eppendorf)
- 15 ml Falcon tubes, Corning, cat # 352096
- P-1000, P200, P20 and P10 tips (Eploret Reload Tip, cat # 02717349, 02717351, 02717341, 02717352, Eppendorf)
- Bio-Rad columns – Poly prep chromatography columns (cat # 731-1550, Bio-Rad)
- Solid phase extraction disk, ultramicrospin column C18 (Cat # SEMSS18V, The Nest group). Be careful, all the columns have the same product number but different volume capacity. Take the one with 5-200 ul.

##### Solutions- (Prepare the same day)

###### Cell lysis buffers(4 suggested 2x lysis buffers)

a. **Chaps buffer**

○ 1.0 % w/v CHAPS (**for 0.5% final**) = 0.1 g in 10 ml of PBS
○ 2.4% w/v CHAPS (**for 1.2% final**) = 0.24 g in 10 ml of PBS
○ Add 1 pellet of Invitrogen protein and proteases inhibitor/10 ml of buffer (Thermo Scientific, A32961)
○ Keep on ice
b. **Igepal 0.5%**

○ 1.0 % w/v Igepal (for 0.5% final) = 0.1 g in 10 ml of PBS +1 pellet of Invitrogen protein and proteases inhibitor/10 ml of buffer (Thermo Scientific, A32961)
c. **NaODEOXY Lysis buffer**: 0.5% sodium deoxycholate, 0.4 mM iodo-acetamide, 2 mM EDTA, 1 pellet/5 ml of combined inhibitor EDTA-free (Thermo Scientific, A32961), 2mM Phenyl-methylsulfonyl fluoride, 2% octyl-beta-d glucopyranoside in PBS

###### Elution of MHC-peptide complexes

- **Buffer C Tris 20 mM pH 8.0:** 157.6g/mol × 20 × 10-3 mol/1000 ml × 1000 ml = 3.14 g Tris HCl in 1L water and bring to pH 8.0
- **Buffer A 150 mM NaCl, 20 mM Tris, pH 8.0:** 58.44g/mol × 150 × 10-3 mol/1000ml × 600 ml = 5.26 g NaCl in 600 ml Buffer C
- **Buffer B 400 mM NaCl, 20 mM Tris, pH 8.0:** 58.44 g/mol × 400 × 10-3 mol/1000 ml × 100 ml = 4.68 g NaCl in 200 ml Buffer C
- **1% TFA:** 100 ul TFA + 9.9 ml water

###### Desalting and elution of the peptides from C18 column

- **0.1% TFA:** 10 ul 100% TFA in 10 ml water
- **80% ACN/0.1% TFA:** 8ml ACN + 10 ul 100% TFA+1.99ml water
- **28% ACN/0.1% TFA:** 2.8 ml ACN + 10 ul 100% TFA +7.2 ml water

###### Labeling of 2.0 ml low binding tubes for each sample

- 1× for the cell solubilization
- 1× for lysate flow through
- 1× for MHC-peptide complexes
- 1× for beads post elution
- 2 × C18 desalting
- 1× for eluted peptides

###### OPTIONAL: Aliquots for BCA and western blotting

- 1 aliquot for protein quantification
- Aliquot A = Coupled beads + lysate pre-incubation
- Aliquot B = unbound MHC-peptide complexes
- Aliquot C= MHC-peptide complexes

##### Cell solubilization

1. Thaw cell pellet in the tube by warming it in the palm of the hand

a. Add 450 ul of PBS to the pellet,
b. pipet up and down 2-3 times
c. Measure the total volume
d. transfer into a 2 ml tube
e. add 2× lysis buffer of the volume measured at step c.
2. Put tubes on a rotating wheel (22 RPM) for 1 hour in the cold room 4°C
3. Spin at 17927 rcf (13000 RPM, Fixed-angle rotor FA-45-48-11, Eppendorf centrifuge 5430R) for 20 minutes at 4°C
4. Recover supernatants (cleared lysate)
5. OPTIONAL: Measure the volume of the cell lysate and take an aliquot of 6 ul for BCA to evaluate protein concentration
6. Recover the prepared beads coupled with antibody of interest (from S23b or S23d)
7. Add cleared lysate to the beads
8. Incubate ON at 4 degrees Celsius on the rotator wheel (10 RPM)

##### Elution of MHC-peptide complexes

NOTE: It could be difficult to elute the liquid from the column just by removing the disposable bottom lid of the column. To increase the speed of liquid elution, cut the tip of the column to increase the size of the hole. Refer to Supplemental data S23g.

1. Cut the bottom of the Biorad column
2. Place onto the Biorad rack and place an empty container underneath to collect flow throughs
3. Wash Biorad column with 10 ml of buffer A

i. Let the buffer drains by gravity
4. Add the mixture of the beads + lysate into the column (the mixture incubated ON)
5. Let drain the lysate by gravity (or collect and store if desired)
6. Wash the beads by adding 10 mL of buffer A and let eluate by gravity the buffer
7. Repeat with 10 mL of buffer B
8. Repeat with 10 mL of buffer A,
9. Repeat with 10 mL of buffer C
10. After the last wash, MHC-peptide complexes are eluted 2× 300 ul of 1 % TFA:

i. Place the column on a new 2.0 ml tube
ii. Add 300 ul of 1% TFA
iii. Mix the beads with the 1% TFA by pipetting up and down 4-5 times directly in the column
iv. Let the 1% TFA solution eluate (about 3 minutes)
v. Repeat b-d

1. Combine flowthroughs = **MHC-peptide complexes + peptides**
11. OPTIONAL: Take an aliquot (Aliquot C, 24 ul) = MHC-peptide complexes
12. OPTIONAL: Following elution, beads are resuspended in 1 ml PBS and kept at 4 degrees:

i. Cap the bottom of the column
ii. Add 1 ml of PBS
iii. Collect beads and keep at 4 degrees

##### Desalting with C18 columns and elution of MHC Peptides

1. Install the column in 2 ml tubes with 2 anneals per column for a better fit in the 2.0 ml tube. These anneals are provided with the columns. Refer to Supplemental data S23g.
2. Add 200 ul of 80% ACN/0.1% TFA to the C18 column
3. Spin 1545 rcf (3700 RPM, Fixed-angle rotor FA-45-48-11, Eppendorf centrifuge 5430R) for 3min at 4°C
4. Aspirate flowthrough
5. Add 200 ul of 0.1% TFA
6. Spin - Aspirate flowthrough
7. Load sample: add 200 ul of sample (eluted MHC-peptide complexes, from step 9f)
8. Spin - Aspirate flowthrough
9. Repeat the loading of the sample 2× (the maximal loading volume of the column is 200 ul)
10. Spin – Aspirate flowthrough
11. Add 200 ul 0.1% TFA
12. Spin - Aspirate flowthrough

a. *Be sure there is no liquid left in the tip before elution. If so, spin again until no residual liquid is visible*
13. Transfer the C18 columns onto new 2.0 ml low bind Eppendorf tubes
14. Elute peptides with 28% ACN/0.1% TFA

a. Add 200 ul of 28% ACN/0.1% TFA
b. Spin
c. Transfer the eluate (peptides) in a new labeled tube
d. Repeat 2× and pool the eluate (peptides) (Final volume = 600 ul.)
15. Store the peptides at −20 degrees Celsius.

#### Tracking efficiency of MHC-peptides elution from the beads by western blotting

In order to evaluate the proportion of MHC-peptides complexes bounded to the beads before and after acidic elution with 1% TFA, it is possible to perform a western blot with aliquots taken at key steps during the protocol. Western blotting shown in Supplemental data 23d reveals as expected, the enrichment of MHC-peptide complexes following acidic elution with 1% TFA. The absence of signal in this fraction would indicate that the elution step was not successful.

**Supplemental Data S24d.**
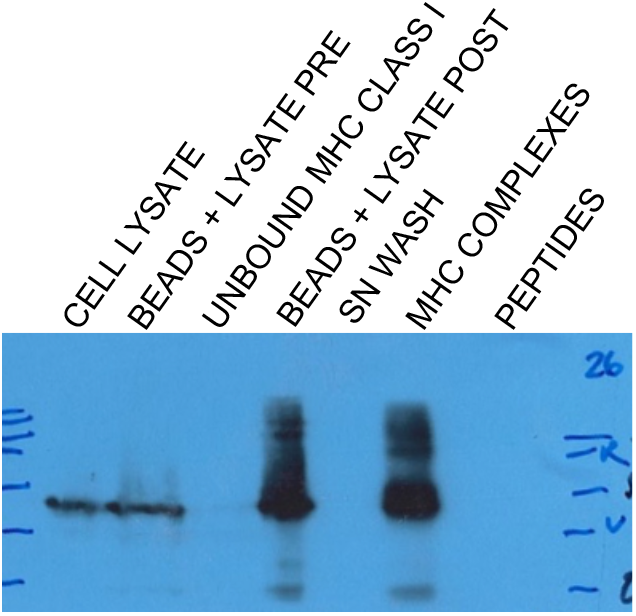
Western blotting for tracking MHC-ligand complexes following acidic elution. Aliquots taken from the steps indicated in the protocol S23b were loaded onto a 12% SDS-PAGE gel and transferred onto nitrocellulose membrane. The strong detection signal of the in the MHC-ligand complexes fraction using anti-HLA-A antibody (Abcam, #ab 52922, 1:1000) confirmed the enrichment of MHC-ligand complexes following immunocapture using Sepharose G beads.

#### S24e: Detailed protocol for immunopurification of MHC Class I using Magnetic Sepharose A/G protein beads

##### Material

- 100 million cells
- Magnetic sepharose beads coupled with your antibody of interest
- Micro tube, 1.5 ml Protein LoBind (cat # 022431081, Eppendorf)
- Micro tube, 2 ml Protein LoBind (cat # 022431002, Eppendorf)
- 15 ml Falcon tubes, Corning, cat # 352096
- P-1000 tips, P200, P20 and P10 (Eploret Reload Tip, cat # 02717349, 02717351, 02717341, 02717352, Eppendorf)
- Bio-Rad columns – Poly prep chromatography columns (cat # 731-1550, Bio-Rad)
- Solid phase extraction disk, ultramicrospin column C18 (Cat # SEMSS18V, The Nest group). Be careful, all the columns have the same product number but different volume capacity. Take the one with 5-200 ul.
- Spin × tube centrifuge tube filter 0.45 micron (Cat#8163, Costar)

##### Solutions- (Prepare the same day)

###### Cell lysis buffers

4 suggested 2× lysis buffers:

a. **Chaps buffer**:

a. 1.0 % w/v CHAPS (**for 0.5% final**) = 0.1 g in 10 ml of PBS
b. 2.4% w/v CHAPS (**for 1.2% final**) = 0.24 g in 10 ml of PBS
c. Add 1 pellet of Invitrogen protein and proteases inhibitor/10 ml of buffer (Thermo Scientific, A32961)
d. Keep on ice
b. **Igepal 0.5%**:

a. 1.0 % w/v Igepal (for 0.5% final) = 0.1 g in 10 ml of PBS +1 pellet of Invitrogen protein and proteases inhibitor/10 ml of buffer (Thermo Scientific, A32961)
c. **NaODEOXY Lysis buffer**: 0.5% sodium deoxycholate, 0.4 mM iodo-acetamide, 2 mM EDTA, 1 pellet/5 ml of combined inhibitor EDTA-free (Thermo Scientific, A32961), 2mM Phenyl-methylsulfonyl fluoride, 2% octyl-beta-d glucopyranoside in PBS

###### Elution of MHC-peptide complexes

- **Buffer C Tris 20 mM pH 8.0:** 157.6g/mol × 20 × 10-3 mol/1000 ml × 1000 ml = 3.14 g Tris HCl in 1L water and bring to pH 8.0
- **Buffer A 150 mM NaCl, 20 mM Tris, pH 8.0:** 58.44g/mol × 150 × 10-3 mol/1000ml × 600 ml = 5.26 g NaCl in 600 ml Buffer C
- **Buffer B 400 mM NaCl, 20 mM Tris, pH 8.0:** 58.44 g/mol × 400 × 10-3 mol/1000 ml × 100 ml = 4.68 g NaCl in 200 ml Buffer C
- **1% TFA:** 100 ul TFA + 9.9 ml water

###### Desalting and elution of the peptides from C18 column

- **0.1% TFA:** 10 ul 100% TFA in 10 ml water
- **80% ACN/0.1% TFA:** 8ml ACN + 10 ul 100% TFA+1.99ml water
- **28% ACN/0.1% TFA:** 2.8 ml ACN + 10 ul 100% TFA +7.2 ml water

###### Labeling of 2.0 ml low binding tubes for each sample

- 1× for the cell solubilization
- 1× for lysate flow through
- 1× for MHC-peptide complexes
- 1× for beads post elution
- 2 × C18 desalting
- 1× for eluted peptides

###### OPTIONAL: Aliquots for BCA and western blotting

- 1 aliquot for protein quantification
- Aliquot A = Coupled beads + lysate pre-incubation
- Aliquot B = unbound MHC-peptide complexes
- Aliquot C= MHC-peptide complexes

##### Cell solubilization

1. Thaw cell pellet in the tube by warming it in the palm of the hand

a. Add 450 ul of PBS to the pellet,
b. Pipet up and down 2-3 times
c. Measure the total volume
d. Transfer into a 2 ml tube
e. Add 2× lysis buffer of the volume measured at step c.
2. Put tubes a rotating wheel (rotation, 20 RPM) for 1 hour in the cold room
3. Spin at 17927 rcf (13000 RPM, Fixed-angle rotor FA-45-48-11, Eppendorf centrifuge 5430R) for 20 minutes at 4°C
4. Recover supernatants (cleared lysate)
5. OPTIONAL: Measure the volume of the cell lysate and take an aliquot of 6 ul for BCA to evaluate protein concentration

##### Elution of MHC-peptide complexes

1. Gently resuspend 1 ml of covalently bound (W6/32) antibody to Protein A magnetic beads from S22f
2. Wash the beads:

a. Place tube on magnet
b. Remove liquid
3. Add 1 ml PBS
4. Vortex 3 seconds
5. Repeat 2x
6. Place tubes on ice
7. Add cleared lysates
8. Vortex 3 seconds
9. OPTIONAL: Take a 24ul aliquot of the coupled beads + lysate pre-incubation ON
10. Incubate at 4°C for 3 hours with rotation (20 rpm)
11. Place on magnet holder
12. Remove unbound lysate
13. OPTIONAL: Take a 24ul aliquot of unbound MHC-peptide complexes
14. Wash beads 8 times (8 × 1 ml) with 1 ml PBS, vortex 3 seconds, place on magnet and remove liquid
15. Wash beads 1 ml with 0.1X PBS, vortex 3 seconds, place on magnet and remove liquid
16. Wash beads 1 ml with water, vortex 3 seconds, place on magnet and remove liquid
17. OPTIONAL: Take a 24ul aliquot of the supernatant of the last wash
18. Elute MHC-peptide complexes

a. Add 200 ul of 1% TFA to the tube
b. Vortex
c. Collect the liquid and transfer in a new 2.0 ml tube
d. Repeat 2× (total of 600 ul)
19. The MHC-peptide complexes eluate (600 ul) is further filtered using a Spin-X tube to remove remaining magnetic beads.
20. Transfer the 600ul MHC-peptide complexes into a Spin-X tube
21. Spin the Spin-x-tube at 1000 rcf (3000 rpm) for 2 minute/4°C
22. Discard filters and keep tubes containing eluted MHC-peptide complexes on ice
23. OPTIONAL: Take a 24ul aliquot of MHC-peptide complexes
24. OPTIONAL: Keep the beads for another round of IP

a. Wash beads with 2 × 1 ml PBS, vortex 3 seconds, place on magnet and remove supernatant
b. Resuspend beads with 1 × 900 ul PBS
c. store at 4°C

##### Desalting with C18 columns and elution of MHC Peptides

1. Install the column in 2 ml tubes with 2 anneals per column for a better fit in the 2.0 ml tube. These anneals are provided with the columns. Refer to Supplemental data S23g.
2. Add 200 ul of 80% ACN/0.1% TFA to the C18 column
3. Spin 1545 rcf (3700 RPM, Fixed-angle rotor FA-45-48-11, Eppendorf centrifuge 5430R) for 3min at 4°C
4. Aspirate flowthrough
5. Add 200 ul of 0.1% TFA
6. Spin - Aspirate flowthrough
7. Load sample: add 200 ul of sample (eluted MHC-peptide complexes, from step 22)
8. Spin - Aspirate flowthrough
9. Repeat the loading of the sample 2× (the maximal loading volume of the column is 200 ul)
10. Spin – Aspirate flowthrough
11. Add 200 ul 0.1% TFA
12. Spin - Aspirate flowthrough

a. *Be sure there is no liquid left in the tip before elution. If so, spin again until no residual liquid is visible*
13. Transfer the C18 columns onto new 2.0 ml low bind Eppendorf tubes
14. Elute peptides with 28% ACN/0.1% TFA

a. Add 200 ul of 28% ACN/0.1% TFA
b. Spin
c. Transfer the eluate (peptides) in a new labeled tube
d. Repeat 2× and pool the eluate (peptides) (Final volume = 600 ul.)
15. Store the peptides at −20 degrees Celsius.

#### Tracking efficiency of MHC-peptides elution from the beads by western blotting

In order to evaluate the proportion of MHC-peptides complexes bounded to the beads before and after acidic elution with 1% TFA, it is possible to perform western with aliquots taken at key steps during the protocol S23e. Western blotting shown in Supplemental data 23f reveals the enrichment of MHC-peptide complexes following acidic elution with 1% TFA. The absence of signal in this fraction would indicate that the elution step was not successful.

**Supplemental Data S24f.**
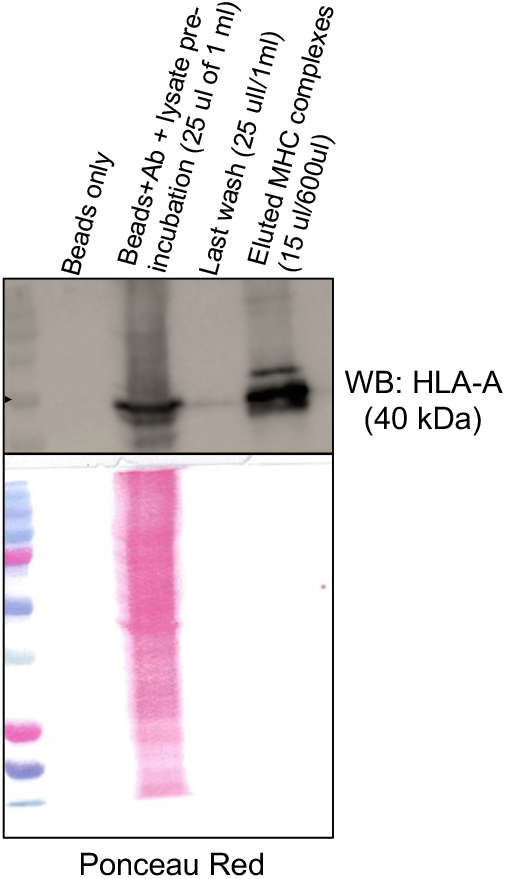
Western blotting for tracking MHC-ligand complexes following acidic elution from magnetic beads. Aliquots taken from the steps indicated in the protocol were loaded onto a 12% SDS-PAGE gel and transferred onto nitrocellulose membrane. The strong detection signal of the in the MHC-ligand complexes fraction using anti-HLA-A antibody (Abcam, #ab 52922, 1:1000) confirmed the enrichment of MHC-ligand complexes following immunocapture using Sepharose G beads. Ponceau red staining is shown as loading control.

#### Supplemental Data S24g

**Supplemental Data S24g.**
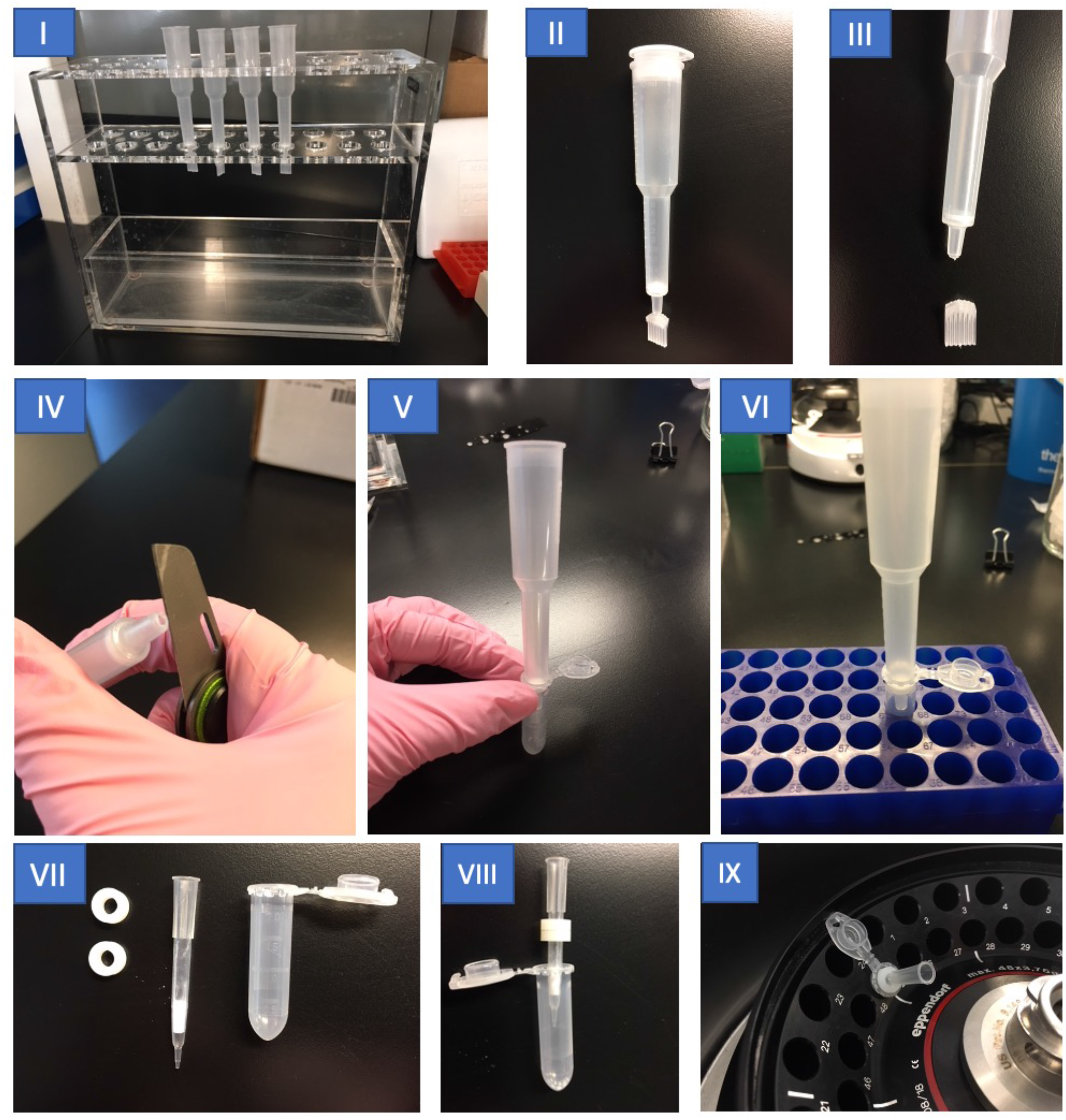
Visual of the Bio-Rad and C18 columns set up for the elution of MHC-peptides complexes peptides. I) Example of the Bio-Rad column holder. II-V) Cutting the bottom of the column after the incubation of the cell lysate with the antibody-coupled beads help to increase the speed of liquid flow by gravity. V-VI) For the elution of MHC-peptide complexes, the column is installed by standing on a 2.0 ml tube, which is placed on a microtube rack. VII-IX) C18 columns are installed on a 2.0 tube using 2 anneals provided with the columns. This set up helps to keep columns in the tube during centrifugation.

#### Supplemental Data S25. Creation of the MhcVizPipe

##### Creatin of the MhcVizPipe

MhcVizPipe (MVP) is a free and open-source bioinformatics and visualization pipeline developed in the Python programming language and is available at https://github.com/CaronLab/MhcVizPipe. It connects the bioinformatic tools NetMHCpan, NetMHCIIpan, and GibbsCluster with Python visualization libraries to create portable HTML reports for interpretation of immunopeptidomics mass spectrometry (MS) data. In addition to NetMHCpan, NetMHCIIpand, and GibbsCluster, it makes use of the following third-party Python libraries: Plotly, PlotlyDash, Dash Bootstrap Components, Pandas, Numpy, Dominate, UpSetPlot, Seaborn, Waitress and PlotlyLogo (developed in-house and describe below).

The raw results from NetMHCpan, NetMHCIIpan, and GibbsCluster are available in a directory used for intermediate results. A copy of the report itself is also kept here. The default location for this directory is /tmp/mhcvizpipe. This location can be customized by editing the config file using the “Settings” button in the GUI, or by directly editing the config file. The config file is named “.mhcvizpipe.config” and is stored in the user’s home directory. Note that the dot prefix in the filename makes it hidden on most systems, so accessing it might require changing your file visibility settings.

###### Installation

The installation of MVP is facilitated by a bash shell script which does the following:

1. Downloads a portable python distribution compatible with the user’s operating system.
2. Installs MhcVizPipe into the Python distribution using Pip.
3. Extracts the NetMHCpan, NetMHCIIpan, and GibbsCluster archives and places them in an appropriate location
4. Modifies the run scripts and generated temp directories for the programs in step 3 so they work properly in their new environment
5. Downloads the required data files for NetMHCpan and NetMHCIIpan and extracts them into the appropriate locations
6. Edits MVP’s config file so it knows where to find the NetMHCpan, netMHCIIpan, and GibbsCluster installations.
7. Adds MVP to users PATH as MhcVizPipe, which is a bash script that starts the GUI and is located in /usr/local/bin

###### Data loading

The interface of MVP is built using the PlotlyDash library and runs as a local web application (i.e. it runs in a web browser). Peptide data is accepted in text file format (e.g. TXT, CSV, and TSV files) or as lists copied and pasted into the graphical user interface.

###### Allele annotation

Subsets of the peptide lists are created as follows:

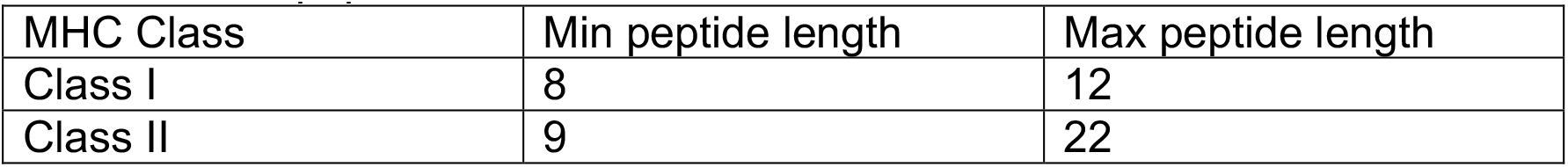

| MHC Class | Min peptide length | Max peptide length |
| --- | --- | --- |
| Class I | 8 | 12 |
| Class II | 9 | 22 |

These lists are analyzed using either NetMHCpan or NetMHCIIpan to yield binding predictions used to annotate the peptides as follows:

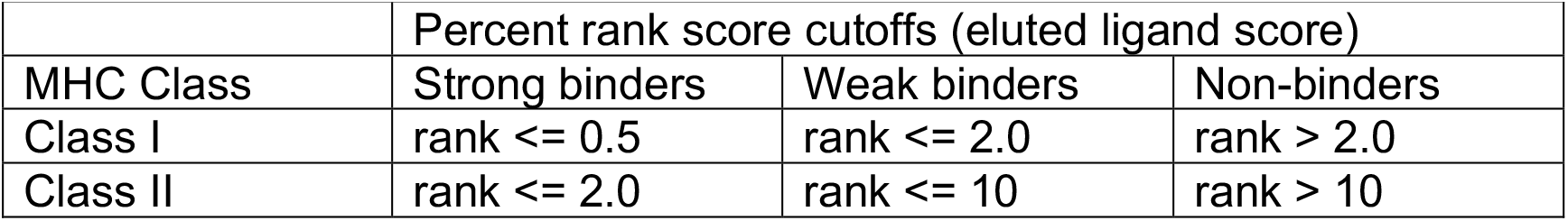

|  | Percent rank score cutoffs (eluted ligand score) |  |  |
| --- | --- | --- | --- |
| MHC Class | Strong binders | Weak binders | Non-binders |
| Class I | rank <= 0.5 | rank <= 2.0 | rank > 2.0 |
| Class II | rank <= 2.0 | rank <= 10 | rank > 10 |

The results are presented in tabular format as well as with a bar plot and heat map. The bar plot shows the number of peptides versus aggregate binding strength (e.g. a peptide that has a weak affinity for one allele and a strong affinity for another is counted as a strong binder). The heatmap is sorted by percent rank eluted ligand score from left-to-right (i.e. the peptides are ordered first by the left-most column, then the second-left-most, etc.). To prevent the convolution of the ordering, all values greater than 2.5 for class I or 12 for class II are set to 2.5 or 12, respectively. The colormap of heatmap is set such that red approximately represents strong binders and blue approximately represents weak binders.

###### GibbsCluster

MVP performs two GibbsCluster routines which we have termed “Unsupervised GibbsCluster” and “Allele-Specific GibbsCluster”. There is a tab for each of these in the “Sequence Motifs” section of the report.

The unsupervised GibbsCluster is a standard GibbsCluster run using all peptides in the subset described above. The following parameters are used, which are the recommended defaults for class I and I peptides, respectively:

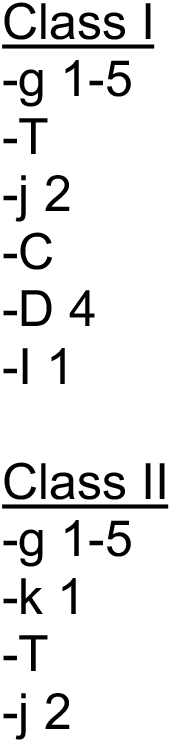

The grouping with the highest KLD score is presented in the report.

The “allele-specific GibbsCluster” is dependent upon the results of NetMHCpan or NetMHCIIpan. A subset of peptides is created for each allele, such that all the peptides in a subset are strong or weak binders for the respective allele. An additional subset is created in which all non-binders peptides for all the alleles are combined. Any subset containing less that 20 peptides is discarded. Each of the remaining subsets is run in GibbsCluster using the above parameters with -g set to 1, forcing GibbsCluster to look for only one peptide group. An exception is the subset of non-binders, in which -g is set to 1-5, allowing GibbsCluster to look for multiple groups in these unannotated peptides. As above, the grouping with the highest KLD score is presented in the report. The results are reportedin the “Allele-Specific GibbsCluster” tab, where we see one motif for each allele present in the sample, and up to 5 motifs for the non-binding peptides.

All the peptide groups shown in the “Sequence Motifs” section are visualized using the PlotlyLogo library as described below.

##### PlotlyLogo

PlotlyLogo is an open-source library for generating sequence logos from sequence alignment data and is available at https://github.com/kevinkovalchik/Plotly-Logo. It is developed in the Python programming language and requires the Python plotting library Plotly, hence the name PlotlyLogo. It is installable as ‘plotly-logo’ from from PyPI using Pip.

The algorithms in PlotlyLogo are based upon the methods described by Thomsen *et al.* for Seq2Logo [1] and in Immunological Bioinformatics [2] (Lund, Nielsen, Lundegaard, Kesmir and Brunak). In brief, sequence alignments are read from text-formatted files and probability matrices are generated using sequence weighting from Hobohm 1 clustering and pseudo count correction as described in the mentioned publication and book. Two types of sequence logos can be generated: Shannon and Kullback-Leibler. The default in MVP is Kullback-Leibler, but this can be changed in the program settings.

PlotlyLogo was designed for the specific purpose of generating amino acid sequence logos from GibbsCluster results as native Python objects using the Plotly plotting framework. As such, the version used in this publication (plotly-logo v0.0.2) does not include much of the functionality of more complete solutions such as Seq2Logo, but it has the advantage of utilizing a modern Python framework (compatible with Python3) and of generating figures as Python objects which can be directly used in other Python code. The choice to develop PlotlyLogo rather than using an existing Python solution such as LogoMaker [3] was influenced by the desire to generate live, interactive figures in a portable HTML format.

1. Martin Christen Frolund Thomsen; Morten Nielsen, Nucleic Acids Research 2012; 40 (W1): W281-W287.
2. Ole Lund, Morten Nielsen, Søren Brunak, Claus Lundegaard, Can Kesmir. MIT Press, 2005
3. Ammar Tareen, Justin B Kinney, Bioinformatics, Volume 36, Issue 7, 1 April 2020, Pages 2272-2274, https://doi.org/10.1093/bioinformatics/btz921

#### Supplemental Data S26

##### Enabling Windows Subsystem for Linux

1. Open *Turn Windows features on or off*

a. either by directly searching for it in the start menu
b. or going clicking in the start menu on

i. Open *Settings*
ii. Click on *Apps*
iii. Under *Related settings* section, click on *Programs and Features*
iv. On the left pane click *Turn Windows features on or off*
2. Check the *Windows Subsystem for Linux* option
3. Click *OK*
4. Click *Restart now*

##### Installing Linux distros using Microsoft Store

1. Open *Microsoft Store* app
2. Search for *Ubuntu*
3. Click the *Get* (or *Install*) button
4. Click the *Launch* button
5. Create a username for Ubuntu and press Enter
6. Specify a password for Ubuntu and press Enter
7. Repeat the password and press Enter to confirm

##### Installing MhcVizPipe

1. Open the Ubuntu app either from the *Microsoft Store* or from the start menu
2. Within the Ubuntu console navigate to the folder containing the MhcVizPipe_install.sh file along with the third party tools by invoking the following command *cd /mnt/XXX* where *XXX* stands for the Windows file path starting with the disk and finishing with the MhcVizPipe folder with forwardlashes and no colon
3. Invoke *chmod +x ./MhcVizPipe_install.sh*
4. Invoke *./MhcVizPipe_install.sh*
5. Answer *y* to all prompted questions
6. Wait for automated installation to finish

##### Using MhcVizPipe

1. Open the *Ubuntu* app either from the *Microsoft Store* or from the start menu
2. Invoke *MhcVizPipe*
3. Open a browser of your choice (e.g. *Chrome* or *Microsoft Edge*) and navigate to http://localhost:8080/

##### Accessing the temp folder of MhcVizPipe

1. Open the Windows *File Explorer*
2. In the path type *\\wsl$\Ubuntu\tmp\mhcvizpipe*
3. Press Enter

