## Supplementary material for "Immunopeptidomics for Dummies: Detailed Experimental Protocols and Rapid, User-Friendly Visualization of MHC I and II Ligand Datasets with MhcVizPipe": All supplementary data and tables: S1_MVP report for MHC class I human dataset_PBMC1 .htm

MhcVizPipe Report


# M

### hc

# V

### iz

# P

### ipe

### - Analysis report

---

**Date:** 2020-09-03

**Submitted by:** Anonymous

**Analysis type:** Class I

**Description of experiment:**

Example of MHC class I human data set

**Alleles:** HLA-A03:01, HLA-A24:02, HLA-B07:02, HLA-B51:01

**Samples:**

PBMC\_1: DOI: 10.7554/eLife.07661

---

### Sample Overview

**Peptide Length Distribution** (maximum of 30 mers)

| Sample | Peptide length | Total peptides | % |
| --- | --- | --- | --- |
| PBMC\_1 | all lengths | 1659 | 100 |
|  | 8-12 mers | 1659 | 100 |
|  | other | 0 | 0 |

---

### Annotation Results

NetMHCpan eluted ligand predictions made for all peptides between 8 & 12 mers, inclusive.
Percent rank cutoffs for strong and weak binders: 0.5 and 2.0.

| Allele | Sample | Total peptides | Strong binders | Weak binders |
| --- | --- | --- | --- | --- |
| HLA-A03:01 | PBMC\_1 | 1659 | 364 (21.9%) | 32 (1.9%) |
| HLA-A24:02 | PBMC\_1 | 1659 | 212 (12.8%) | 28 (1.7%) |
| HLA-B07:02 | PBMC\_1 | 1659 | 570 (34.4%) | 144 (8.7%) |
| HLA-B51:01 | PBMC\_1 | 1659 | 479 (28.9%) | 242 (14.6%) |

**Binding Affinities**

---

### Binding Heatmaps

---

### Sequence Motifs

Clustering performed with all peptides between 8 & 12 mers, inclusive.

Percentages represent the percentage of peptides in a given group predicted to strongly bind the indicated allele.

Polar

Neutral

Basic

Acidic

Hydrophobic

- Unsupervised GibbsCluster
- Allele-specific GibbsCluster

**PBMC\_1**

Peptides in group: 570

HLA-A03:01: 0%,

HLA-A24:02: 1%,

**HLA-B07:02: 83%,** 

HLA-B51:01: 19%

Peptides in group: 409

HLA-A03:01: 0%,

HLA-A24:02: 0%,

HLA-B07:02: 24%,

**HLA-B51:01: 90%**

Peptides in group: 411

**HLA-A03:01: 89%,** 

HLA-A24:02: 0%,

HLA-B07:02: 0%,

HLA-B51:01: 0%

Peptides in group: 238

HLA-A03:01: 0%,

**HLA-A24:02: 87%,** 

HLA-B07:02: 0%,

HLA-B51:01: 0%

**PBMC\_1 sequence motif(s)**

**HLA-A03:01**

Peptides: 395

**HLA-A24:02**

Peptides: 240

**HLA-B07:02**

Peptides: 710

**HLA-B51:01**

Peptides: 715

**Non-binders group 1**

Peptides: 55

**Non-binders group 2**

Peptides: 27
