## Supplementary material for "Immunopeptidomics for Dummies: Detailed Experimental Protocols and Rapid, User-Friendly Visualization of MHC I and II Ligand Datasets with MhcVizPipe": All supplementary data and tables: S2_MVP report for MHC class I human dataset_PBMC2 .htm

MhcVizPipe Report


# M

### hc

# V

### iz

# P

### ipe

### - Analysis report

---

**Date:** 2020-09-03

**Submitted by:** Anonymous

**Analysis type:** Class I

**Description of experiment:**

Example of Human Class I data set

**Alleles:** HLA-A02:01, HLA-A03:01, HLA-B35:01, HLA-B37:01

**Samples:**

PBMC\_2\_: DOI: 10.7554/eLife.07661

---

### Sample Overview

**Peptide Length Distribution** (maximum of 30 mers)

| Sample | Peptide length | Total peptides | % |
| --- | --- | --- | --- |
| PBMC\_2\_ | all lengths | 1364 | 100 |
|  | 8-12 mers | 1364 | 100 |
|  | other | 0 | 0 |

---

### Annotation Results

NetMHCpan eluted ligand predictions made for all peptides between 8 & 12 mers, inclusive.
Percent rank cutoffs for strong and weak binders: 0.5 and 2.0.

| Allele | Sample | Total peptides | Strong binders | Weak binders |
| --- | --- | --- | --- | --- |
| HLA-A02:01 | PBMC\_2\_ | 1364 | 267 (19.6%) | 40 (2.9%) |
| HLA-A03:01 | PBMC\_2\_ | 1364 | 356 (26.1%) | 55 (4.0%) |
| HLA-B35:01 | PBMC\_2\_ | 1364 | 240 (17.6%) | 65 (4.8%) |
| HLA-B37:01 | PBMC\_2\_ | 1364 | 13 (1.0%) | 225 (16.5%) |

**Binding Affinities**

---

### Binding Heatmaps

---

### Sequence Motifs

Clustering performed with all peptides between 8 & 12 mers, inclusive.

Percentages represent the percentage of peptides in a given group predicted to strongly bind the indicated allele.

Polar

Neutral

Basic

Acidic

Hydrophobic

- Unsupervised GibbsCluster
- Allele-specific GibbsCluster

**PBMC\_2\_**

Peptides in group: 302

**HLA-A02:01: 84%,** 

HLA-A03:01: 1%,

HLA-B35:01: 1%,

HLA-B37:01: 0%

Peptides in group: 334

HLA-A02:01: 1%,

HLA-A03:01: 0%,

HLA-B35:01: 1%,

**HLA-B37:01: 4%**

Peptides in group: 426

HLA-A02:01: 2%,

**HLA-A03:01: 82%,** 

HLA-B35:01: 2%,

HLA-B37:01: 0%

Peptides in group: 255

HLA-A02:01: 0%,

HLA-A03:01: 2%,

**HLA-B35:01: 88%,** 

HLA-B37:01: 0%

**PBMC\_2\_ sequence motif(s)**

**HLA-A02:01**

Peptides: 302

**HLA-A03:01**

Peptides: 406

**HLA-B35:01**

Peptides: 299

**HLA-B37:01**

Peptides: 236

**Non-binders group 1**

Peptides: 72

**Non-binders group 2**

Peptides: 163

**Non-binders group 3**

Peptides: 56
