## Supplementary material for "Immunopeptidomics for Dummies: Detailed Experimental Protocols and Rapid, User-Friendly Visualization of MHC I and II Ligand Datasets with MhcVizPipe": All supplementary data and tables: S3_MVP report for MHC class II human dataset_JY .htm

MhcVizPipe Report


# M

### hc

# V

### iz

# P

### ipe

### - Analysis report

---

**Date:** 2020-09-03

**Submitted by:** Anonymous

**Analysis type:** Class II

**Description of experiment:**

Example of human MCH class II dataset

**Alleles:** DRB1\_0404, DRB1\_1301

**Samples:**

JY\_HLA\_DR\_IP: Nature biotechnology, 37(11), 1283–1286. https://doi.org/10.1038/s41587-019-0289-6

---

### Sample Overview

**Peptide Length Distribution** (maximum of 30 mers)

| Sample | Peptide length | Total peptides | % |
| --- | --- | --- | --- |
| JY\_HLA\_DR\_IP | all lengths | 5464 | 100 |
|  | 9-22 mers | 5402 | 99 |
|  | other | 62 | 1 |

---

### Annotation Results

NetMHCIIpan eluted ligand predictions made for all peptides between 9 & 22 mers, inclusive.
Percent rank cutoffs for strong and weak binders: 2.0 and 10.0.

| Allele | Sample | Total peptides | Strong binders | Weak binders |
| --- | --- | --- | --- | --- |
| DRB1\_0404 | JY\_HLA\_DR\_IP | 5402 | 1711 (31.7%) | 921 (17.0%) |
| DRB1\_1301 | JY\_HLA\_DR\_IP | 5402 | 1158 (21.4%) | 1329 (24.6%) |

**Binding Affinities**

---

### Binding Heatmaps

---

### Sequence Motifs

Clustering performed with all peptides between 9 & 22 mers, inclusive.

Percentages represent the percentage of peptides in a given group predicted to strongly bind the indicated allele.

Polar

Neutral

Basic

Acidic

Hydrophobic

- Unsupervised GibbsCluster
- Allele-specific GibbsCluster

**JY\_HLA\_DR\_IP**

Peptides in group: 1335

DRB1\_0404: 33%,

**DRB1\_1301: 40%**

Peptides in group: 1096

**DRB1\_0404: 24%,** 

DRB1\_1301: 19%

Peptides in group: 1094

**DRB1\_0404: 22%,** 

DRB1\_1301: 21%

Peptides in group: 1704

**DRB1\_0404: 44%,** 

DRB1\_1301: 10%

**JY\_HLA\_DR\_IP sequence motif(s)**

**DRB1\_0404**

Peptides: 2563

**DRB1\_1301**

Peptides: 2407

**Non-binders group 1**

Peptides: 557

**Non-binders group 2**

Peptides: 623
