## Supplementary material for "Immunopeptidomics for Dummies: Detailed Experimental Protocols and Rapid, User-Friendly Visualization of MHC I and II Ligand Datasets with MhcVizPipe": All supplementary data and tables: S4_MVP report Effect of lysis buffer BLCL .htm

MhcVizPipe Report


# M

### hc

# V

### iz

# P

### ipe

### - Analysis report

---

**Date:** 2020-09-02

**Submitted by:** Anonymous

**Analysis type:** Class I

**Description of experiment:**

EFFEC OF LYSIS BUFFER

**Alleles:** HLA-B44:02, HLA-B08:01, HLA-A03:01, HLA-A29:01

**Samples:**

Chaps
NAODEOXY

**Species:** HUMAN

**# of cells:** 100 millions

**Lysis buffer:** Chaps vs NaDeoxycholate

**Type of beads:** Magnetic A

**Antibody:** W6/32

**Incubation time:** Overnight

**MHC-ligand complex elution buffer:** 0.2% formic acid

**Peptide elution buffer:** 28% ACN

**Type of MS/MS:** Fusion

**Peptide identification software:** Peaks

**Peptide FDR:** 1% FDR

---

### Sample Overview

**UpSet Plot**

Only intersections > 0.5% are displayed

**Peptide Length Distribution** (maximum of 30 mers)

| Sample | Peptide length | Total peptides | % |
| --- | --- | --- | --- |
| Chaps | all lengths | 8098 | 100 |
|  | 8-12 mers | 7662 | 95 |
|  | other | 436 | 5 |
| NAODEOXY | all lengths | 8321 | 100 |
|  | 8-12 mers | 7930 | 95 |
|  | other | 391 | 5 |

---

### Annotation Results

NetMHCpan eluted ligand predictions made for all peptides between 8 & 12 mers, inclusive.
Percent rank cutoffs for strong and weak binders: 0.5 and 2.0.

| Allele | Sample | Total peptides | Strong binders | Weak binders |
| --- | --- | --- | --- | --- |
| HLA-B44:02 | Chaps | 7662 | 3015 (39.4%) | 192 (2.5%) |
|  | NAODEOXY | 7930 | 3077 (38.8%) | 183 (2.3%) |
| HLA-B08:01 | Chaps | 7662 | 1733 (22.6%) | 277 (3.6%) |
|  | NAODEOXY | 7930 | 1865 (23.5%) | 302 (3.8%) |
| HLA-A03:01 | Chaps | 7662 | 1612 (21.0%) | 380 (5.0%) |
|  | NAODEOXY | 7930 | 1685 (21.2%) | 384 (4.8%) |
| HLA-A29:01 | Chaps | 7662 | 812 (10.6%) | 767 (10.0%) |
|  | NAODEOXY | 7930 | 806 (10.2%) | 779 (9.8%) |

**Binding Affinities**

---

### Binding Heatmaps

---

### Sequence Motifs

Clustering performed with all peptides between 8 & 12 mers, inclusive.

Percentages represent the percentage of peptides in a given group predicted to strongly bind the indicated allele.

Polar

Neutral

Basic

Acidic

Hydrophobic

- Unsupervised GibbsCluster
- Allele-specific GibbsCluster

**Chaps**

Peptides in group: 1991

HLA-B44:02: 0%,

**HLA-B08:01: 87%,** 

HLA-A03:01: 0%,

HLA-A29:01: 1%

Peptides in group: 2236

HLA-B44:02: 0%,

HLA-B08:01: 0%,

**HLA-A03:01: 71%,** 

HLA-A29:01: 28%

Peptides in group: 3219

**HLA-B44:02: 93%,** 

HLA-B08:01: 0%,

HLA-A03:01: 0%,

HLA-A29:01: 4%

**NAODEOXY**

Peptides in group: 2152

HLA-B44:02: 0%,

**HLA-B08:01: 86%,** 

HLA-A03:01: 0%,

HLA-A29:01: 1%

Peptides in group: 2324

HLA-B44:02: 0%,

HLA-B08:01: 0%,

**HLA-A03:01: 72%,** 

HLA-A29:01: 27%

Peptides in group: 3253

**HLA-B44:02: 94%,** 

HLA-B08:01: 0%,

HLA-A03:01: 0%,

HLA-A29:01: 4%

**Chaps sequence motif(s)**

**HLA-B44:02**

Peptides: 3187

**HLA-B08:01**

Peptides: 1977

**HLA-A03:01**

Peptides: 1970

**HLA-A29:01**

Peptides: 1539

**Non-binders group 1**

Peptides: 59

**Non-binders group 2**

Peptides: 64

**Non-binders group 3**

Peptides: 105

**Non-binders group 4**

Peptides: 33

**Non-binders group 5**

Peptides: 63

**NAODEOXY sequence motif(s)**

**HLA-B44:02**

Peptides: 3247

**HLA-B08:01**

Peptides: 2133

**HLA-A03:01**

Peptides: 2051

**HLA-A29:01**

Peptides: 1548

**Non-binders group 1**

Peptides: 68

**Non-binders group 2**

Peptides: 98

**Non-binders group 3**

Peptides: 72

**Non-binders group 4**

Peptides: 94
