## Supplementary material for "Immunopeptidomics for Dummies: Detailed Experimental Protocols and Rapid, User-Friendly Visualization of MHC I and II Ligand Datasets with MhcVizPipe": All supplementary data and tables: S5_MVP report Effect of lysis buffer JY .htm

MhcVizPipe Report


# M

### hc

# V

### iz

# P

### ipe

### - Analysis report

---

**Date:** 2020-09-02

**Submitted by:** Anonymous

**Analysis type:** Class I

**Description of experiment:**

Effect of lysis buffer J&

**Alleles:** HLA-A02:01, HLA-B07:02

**Samples:**

CHAPS
NADEOXY

**Species:** HUMAN - JY CELLS

**# of cells:** 100 MILLIONS

**Lysis buffer:** CHAPS VS NADEOXY

**Type of beads:** MAGNETIC A

**Antibody:** W6-32

**Incubation time:** OVERNIGHT

**MHC-ligand complex elution buffer:** 0.2% FORMIC ACID

**Peptide elution buffer:** ACN 28%

**Type of MS/MS:** FUSION

**Peptide identification software:** PEAKS

**Peptide FDR:** 1%

---

### Sample Overview

**UpSet Plot**

Only intersections > 0.5% are displayed

**Peptide Length Distribution** (maximum of 30 mers)

| Sample | Peptide length | Total peptides | % |
| --- | --- | --- | --- |
| CHAPS | all lengths | 5640 | 100 |
|  | 8-12 mers | 5412 | 96 |
|  | other | 228 | 4 |
| NADEOXY | all lengths | 5779 | 100 |
|  | 8-12 mers | 5562 | 96 |
|  | other | 217 | 4 |

---

### Annotation Results

NetMHCpan eluted ligand predictions made for all peptides between 8 & 12 mers, inclusive.
Percent rank cutoffs for strong and weak binders: 0.5 and 2.0.

| Allele | Sample | Total peptides | Strong binders | Weak binders |
| --- | --- | --- | --- | --- |
| HLA-A02:01 | CHAPS | 5412 | 1994 (36.8%) | 304 (5.6%) |
|  | NADEOXY | 5562 | 2029 (36.5%) | 289 (5.2%) |
| HLA-B07:02 | CHAPS | 5412 | 2525 (46.7%) | 400 (7.4%) |
|  | NADEOXY | 5562 | 2701 (48.6%) | 411 (7.4%) |

**Binding Affinities**

---

### Binding Heatmaps

---

### Sequence Motifs

Clustering performed with all peptides between 8 & 12 mers, inclusive.

Percentages represent the percentage of peptides in a given group predicted to strongly bind the indicated allele.

Polar

Neutral

Basic

Acidic

Hydrophobic

- Unsupervised GibbsCluster
- Allele-specific GibbsCluster

**CHAPS**

Peptides in group: 2855

HLA-A02:01: 0%,

**HLA-B07:02: 88%**

Peptides in group: 2375

**HLA-A02:01: 83%,** 

HLA-B07:02: 0%

**NADEOXY**

Peptides in group: 3009

HLA-A02:01: 0%,

**HLA-B07:02: 89%**

Peptides in group: 2386

**HLA-A02:01: 84%,** 

HLA-B07:02: 1%

**CHAPS sequence motif(s)**

**HLA-A02:01**

Peptides: 2277

**HLA-B07:02**

Peptides: 2889

**Non-binders group 1**

Peptides: 138

**Non-binders group 2**

Peptides: 222

**NADEOXY sequence motif(s)**

**HLA-A02:01**

Peptides: 2300

**HLA-B07:02**

Peptides: 3082

**Non-binders group 1**

Peptides: 162

**Non-binders group 2**

Peptides: 74

**Non-binders group 3**

Peptides: 65

**Non-binders group 4**

Peptides: 67
