## Supplementary material for "Immunopeptidomics for Dummies: Detailed Experimental Protocols and Rapid, User-Friendly Visualization of MHC I and II Ligand Datasets with MhcVizPipe": All supplementary data and tables: S6_MVP report Effect of beads JY TFA 1%.htm

MhcVizPipe Report


# M

### hc

# V

### iz

# P

### ipe

### - Analysis report

---

**Date:** 2020-09-02

**Submitted by:** Anonymous

**Analysis type:** Class I

**Description of experiment:**

Effect of beads JY

**Alleles:** HLA-A02:01, HLA-B07:02

**Samples:**

MAGA
CNBR
SEPG

**Species:** Human JY cellls

**# of cells:** 100 millions

**Lysis buffer:** chaps

**Type of beads:** variable

**Antibody:** W6/32

**Incubation time:** OVERNIGHT

**MHC-ligand complex elution buffer:** 1% TFA

**Peptide elution buffer:** 28% ACN

**Type of MS/MS:** FUSION

**Peptide identification software:** PEAKS

**Peptide FDR:** 1%

---

### Sample Overview

**UpSet Plot**

Only intersections > 0.5% are displayed

**Peptide Length Distribution** (maximum of 30 mers)

| Sample | Peptide length | Total peptides | % |
| --- | --- | --- | --- |
| MAGA | all lengths | 5640 | 100 |
|  | 8-12 mers | 5412 | 96 |
|  | other | 228 | 4 |
| CNBR | all lengths | 4657 | 100 |
|  | 8-12 mers | 4329 | 93 |
|  | other | 328 | 7 |
| SEPG | all lengths | 4233 | 100 |
|  | 8-12 mers | 4072 | 96 |
|  | other | 161 | 4 |

---

### Annotation Results

NetMHCpan eluted ligand predictions made for all peptides between 8 & 12 mers, inclusive.
Percent rank cutoffs for strong and weak binders: 0.5 and 2.0.

| Allele | Sample | Total peptides | Strong binders | Weak binders |
| --- | --- | --- | --- | --- |
| HLA-A02:01 | MAGA | 5412 | 1994 (36.8%) | 304 (5.6%) |
|  | CNBR | 4329 | 1996 (46.1%) | 234 (5.4%) |
|  | SEPG | 4072 | 1721 (42.3%) | 226 (5.6%) |
| HLA-B07:02 | MAGA | 5412 | 2525 (46.7%) | 400 (7.4%) |
|  | CNBR | 4329 | 1776 (41.0%) | 292 (6.7%) |
|  | SEPG | 4072 | 1750 (43.0%) | 277 (6.8%) |

**Binding Affinities**

---

### Binding Heatmaps

---

### Sequence Motifs

Clustering performed with all peptides between 8 & 12 mers, inclusive.

Percentages represent the percentage of peptides in a given group predicted to strongly bind the indicated allele.

Polar

Neutral

Basic

Acidic

Hydrophobic

- Unsupervised GibbsCluster
- Allele-specific GibbsCluster

**MAGA**

Peptides in group: 2855

HLA-A02:01: 0%,

**HLA-B07:02: 88%**

Peptides in group: 2375

**HLA-A02:01: 83%,** 

HLA-B07:02: 0%

**CNBR**

Peptides in group: 1958

HLA-A02:01: 0%,

**HLA-B07:02: 90%**

Peptides in group: 2270

**HLA-A02:01: 87%,** 

HLA-B07:02: 0%

**SEPG**

Peptides in group: 1952

HLA-A02:01: 0%,

**HLA-B07:02: 89%**

Peptides in group: 1993

**HLA-A02:01: 86%,** 

HLA-B07:02: 0%

**MAGA sequence motif(s)**

**HLA-A02:01**

Peptides: 2277

**HLA-B07:02**

Peptides: 2889

**Non-binders group 1**

Peptides: 138

**Non-binders group 2**

Peptides: 222

**CNBR sequence motif(s)**

**HLA-A02:01**

Peptides: 2218

**HLA-B07:02**

Peptides: 2053

**Non-binders group 1**

Peptides: 52

**Non-binders group 2**

Peptides: 105

**Non-binders group 3**

Peptides: 22

**Non-binders group 4**

Peptides: 15

**Non-binders group 5**

Peptides: 33

**SEPG sequence motif(s)**

**HLA-A02:01**

Peptides: 1925

**HLA-B07:02**

Peptides: 2006

**Non-binders group 1**

Peptides: 65

**Non-binders group 2**

Peptides: 37

**Non-binders group 3**

Peptides: 48

**Non-binders group 4**

Peptides: 102
