## Supplementary material for "Immunopeptidomics for Dummies: Detailed Experimental Protocols and Rapid, User-Friendly Visualization of MHC I and II Ligand Datasets with MhcVizPipe": All supplementary data and tables: S7_MVP report Effect of incubation time JY TFA 1%.htm

MhcVizPipe Report


# M

### hc

# V

### iz

# P

### ipe

### - Analysis report

---

**Date:** 2020-09-02

**Submitted by:** Anonymous

**Analysis type:** Class I

**Description of experiment:**

None provided

**Alleles:** HLA-A02:01, HLA-B07:02

**Samples:**

90\_min\_R.T.\_R1
90\_min\_R.T.\_R2
90\_min\_R.T.\_R3
ON\_4degrees\_R1
ON\_4\_degrees\_R2
ON\_4\_degrees\_R3

**Species:** Human JY

**# of cells:** 100 MILLIONS

**Lysis buffer:** CHAPS

**Type of beads:** CNBR

**Antibody:** W6/32

**Incubation time:** VARIABLE

**MHC-ligand complex elution buffer:** 1% TFA

**Peptide elution buffer:** 28% ACN

**Type of MS/MS:** FUSION

**Peptide identification software:** PEAKS

**Peptide FDR:** 1%

---

### Sample Overview

**UpSet Plot**

Only intersections > 0.5% are displayed

**Peptide Length Distribution** (maximum of 30 mers)

| Sample | Peptide length | Total peptides | % |
| --- | --- | --- | --- |
| 90\_min\_R.T.\_R1 | all lengths | 3730 | 100 |
|  | 8-12 mers | 3448 | 92 |
|  | other | 282 | 8 |
| 90\_min\_R.T.\_R2 | all lengths | 3445 | 100 |
|  | 8-12 mers | 2981 | 87 |
|  | other | 464 | 13 |
| 90\_min\_R.T.\_R3 | all lengths | 3871 | 100 |
|  | 8-12 mers | 3624 | 94 |
|  | other | 247 | 6 |
| ON\_4degrees\_R1 | all lengths | 4135 | 100 |
|  | 8-12 mers | 3858 | 93 |
|  | other | 277 | 7 |
| ON\_4\_degrees\_R2 | all lengths | 3943 | 100 |
|  | 8-12 mers | 3598 | 91 |
|  | other | 345 | 9 |
| ON\_4\_degrees\_R3 | all lengths | 3349 | 100 |
|  | 8-12 mers | 3036 | 91 |
|  | other | 313 | 9 |

---

### Annotation Results

NetMHCpan eluted ligand predictions made for all peptides between 8 & 12 mers, inclusive.
Percent rank cutoffs for strong and weak binders: 0.5 and 2.0.

| Allele | Sample | Total peptides | Strong binders | Weak binders |
| --- | --- | --- | --- | --- |
| HLA-A02:01 | 90\_min\_R.T.\_R1 | 3448 | 1578 (45.8%) | 195 (5.7%) |
|  | 90\_min\_R.T.\_R2 | 2981 | 1341 (45.0%) | 155 (5.2%) |
|  | 90\_min\_R.T.\_R3 | 3624 | 1630 (45.0%) | 194 (5.4%) |
|  | ON\_4degrees\_R1 | 3858 | 1647 (42.7%) | 207 (5.4%) |
|  | ON\_4\_degrees\_R2 | 3598 | 1549 (43.1%) | 182 (5.1%) |
|  | ON\_4\_degrees\_R3 | 3036 | 1392 (45.8%) | 148 (4.9%) |
| HLA-B07:02 | 90\_min\_R.T.\_R1 | 3448 | 1362 (39.5%) | 250 (7.3%) |
|  | 90\_min\_R.T.\_R2 | 2981 | 1199 (40.2%) | 209 (7.0%) |
|  | 90\_min\_R.T.\_R3 | 3624 | 1439 (39.7%) | 275 (7.6%) |
|  | ON\_4degrees\_R1 | 3858 | 1671 (43.3%) | 281 (7.3%) |
|  | ON\_4\_degrees\_R2 | 3598 | 1527 (42.4%) | 259 (7.2%) |
|  | ON\_4\_degrees\_R3 | 3036 | 1230 (40.5%) | 212 (7.0%) |

**Binding Affinities**

---

### Binding Heatmaps

---

### Sequence Motifs

Clustering performed with all peptides between 8 & 12 mers, inclusive.

Percentages represent the percentage of peptides in a given group predicted to strongly bind the indicated allele.

Polar

Neutral

Basic

Acidic

Hydrophobic

- Unsupervised GibbsCluster
- Allele-specific GibbsCluster

**90\_min\_R.T.\_R1**

Peptides in group: 1796

**HLA-A02:01: 87%,** 

HLA-B07:02: 0%

Peptides in group: 1558

HLA-A02:01: 0%,

**HLA-B07:02: 87%**

**90\_min\_R.T.\_R2**

Peptides in group: 1537

**HLA-A02:01: 87%,** 

HLA-B07:02: 0%

Peptides in group: 1364

HLA-A02:01: 0%,

**HLA-B07:02: 87%**

**90\_min\_R.T.\_R3**

Peptides in group: 1857

**HLA-A02:01: 87%,** 

HLA-B07:02: 0%

Peptides in group: 1670

HLA-A02:01: 0%,

**HLA-B07:02: 86%**

**ON\_4degrees\_R1**

Peptides in group: 1899

**HLA-A02:01: 86%,** 

HLA-B07:02: 0%

Peptides in group: 1866

HLA-A02:01: 0%,

**HLA-B07:02: 89%**

**ON\_4\_degrees\_R2**

Peptides in group: 1765

**HLA-A02:01: 87%,** 

HLA-B07:02: 0%

Peptides in group: 1718

HLA-A02:01: 0%,

**HLA-B07:02: 88%**

**ON\_4\_degrees\_R3**

Peptides in group: 1574

**HLA-A02:01: 88%,** 

HLA-B07:02: 0%

Peptides in group: 1391

HLA-A02:01: 0%,

**HLA-B07:02: 88%**

**90\_min\_R.T.\_R1 sequence motif(s)**

**HLA-A02:01**

Peptides: 1752

**HLA-B07:02**

Peptides: 1598

**Non-binders group 1**

Peptides: 18

**Non-binders group 2**

Peptides: 24

**Non-binders group 3**

Peptides: 51

**Non-binders group 4**

Peptides: 94

**Non-binders group 5**

Peptides: 26

**90\_min\_R.T.\_R2 sequence motif(s)**

**HLA-A02:01**

Peptides: 1482

**HLA-B07:02**

Peptides: 1395

**Non-binders group 1**

Peptides: 46

**Non-binders group 2**

Peptides: 22

**Non-binders group 3**

Peptides: 23

**Non-binders group 4**

Peptides: 28

**Non-binders group 5**

Peptides: 80

**90\_min\_R.T.\_R3 sequence motif(s)**

**HLA-A02:01**

Peptides: 1806

**HLA-B07:02**

Peptides: 1699

**Non-binders group 1**

Peptides: 34

**Non-binders group 2**

Peptides: 20

**Non-binders group 3**

Peptides: 87

**Non-binders group 4**

Peptides: 63

**Non-binders group 5**

Peptides: 31

**ON\_4degrees\_R1 sequence motif(s)**

**HLA-A02:01**

Peptides: 1834

**HLA-B07:02**

Peptides: 1930

**Non-binders group 1**

Peptides: 43

**Non-binders group 2**

Peptides: 49

**Non-binders group 3**

Peptides: 99

**Non-binders group 4**

Peptides: 30

**ON\_4\_degrees\_R2 sequence motif(s)**

**HLA-A02:01**

Peptides: 1711

**HLA-B07:02**

Peptides: 1761

**Non-binders group 1**

Peptides: 121

**Non-binders group 2**

Peptides: 89

**ON\_4\_degrees\_R3 sequence motif(s)**

**HLA-A02:01**

Peptides: 1527

**HLA-B07:02**

Peptides: 1429

**Non-binders group 1**

Peptides: 63

**Non-binders group 2**

Peptides: 34

**Non-binders group 3**

Peptides: 74
