## Supplementary material for "Immunopeptidomics for Dummies: Detailed Experimental Protocols and Rapid, User-Friendly Visualization of MHC I and II Ligand Datasets with MhcVizPipe": All supplementary data and tables: S8_MVP Report Effect of beads EL4.htm

MhcVizPipe Report


# M

### hc

# V

### iz

# P

### ipe

### - Analysis report

---

**Date:** 2020-10-06

**Submitted by:** Anonymous

**Analysis type:** Class I

**Description of experiment:**

Figure 3 - Effect of Beads Y3 antibody EL4 cells

**Alleles:** H2-Kb

**Samples:**

CNBR
SEPG
SEPA
MAGG

**Species:** Mouse

**# of cells:** 100 millions

**Lysis buffer:** 0.5% chaps

**Type of beads:** Variable

**Antibody:** Y3 (H2Kb)

**Incubation time:** ON

**MHC-ligand complex elution buffer:** 1% TFA

**Peptide elution buffer:** 28% ACN/0.1% TFA

**Type of MS/MS:** Fusion

**Peptide identification software:** Peaks

**Peptide FDR:** 1%

---

### Sample Overview

**UpSet Plot**

Only intersections > 0.5% are displayed

**Peptide Length Distribution** (maximum of 30 mers)

| Sample | Peptide length | Total peptides | % |
| --- | --- | --- | --- |
| CNBR | all lengths | 1288 | 100 |
|  | 8-12 mers | 1164 | 90 |
|  | other | 124 | 10 |
| SEPG | all lengths | 1132 | 100 |
|  | 8-12 mers | 1027 | 91 |
|  | other | 105 | 9 |
| SEPA | all lengths | 568 | 100 |
|  | 8-12 mers | 507 | 89 |
|  | other | 61 | 11 |
| MAGG | all lengths | 542 | 100 |
|  | 8-12 mers | 448 | 83 |
|  | other | 94 | 17 |

---

### Annotation Results

NetMHCpan eluted ligand predictions made for all peptides between 8 & 12 mers, inclusive.
Percent rank cutoffs for strong and weak binders: 0.5 and 2.0.

| Allele | Sample | Total peptides | Strong binders | Weak binders | Non-binders |
| --- | --- | --- | --- | --- | --- |
| H2-Kb | CNBR | 1164 | 950 (81.6%) | 114 (9.8%) | 100 (8.6%) |
|  | SEPG | 1027 | 861 (83.8%) | 72 (7.0%) | 94 (9.2%) |
|  | SEPA | 507 | 419 (82.6%) | 35 (6.9%) | 53 (10.5%) |
|  | MAGG | 448 | 311 (69.4%) | 39 (8.7%) | 98 (21.9%) |

**Binding Affinities**

---

### Binding Heatmaps

---

### Sequence Motifs

Clustering performed with all peptides between 8 & 12 mers, inclusive.

Percentages represent the percentage of peptides in a given group predicted to strongly bind the indicated allele.

Polar

Neutral

Basic

Acidic

Hydrophobic

- Unsupervised GibbsCluster
- Allele-specific GibbsCluster

**CNBR**

Peptides in group: 1117

**H2-Kb: 85%**

**SEPG**

Peptides in group: 968

**H2-Kb: 89%**

**SEPA**

Peptides in group: 470

**H2-Kb: 89%**

**MAGG**

Peptides in group: 400

**H2-Kb: 77%**

**CNBR sequence motif(s)**

**H2-Kb**

Peptides: 1052

**Non-binders group 1**

Peptides: 20

**Non-binders group 3**

Peptides: 57

**Non-binders group 2**

Peptides: 4

**Non-binders group 4**

Peptides: 11

**SEPG sequence motif(s)**

**H2-Kb**

Peptides: 924

**Non-binders group 1**

Peptides: 15

**Non-binders group 3**

Peptides: 13

**Non-binders group 2**

Peptides: 25

**Non-binders group 4**

Peptides: 35

**SEPA sequence motif(s)**

**H2-Kb**

Peptides: 450

**Non-binders group 2**

Peptides: 14

**Non-binders group 3**

Peptides: 11

**Non-binders group 1**

Peptides: 21

**MAGG sequence motif(s)**

**H2-Kb**

Peptides: 349

**Non-binders group 2**

Peptides: 32

**Non-binders group 3**

Peptides: 26

**Non-binders group 1**

Peptides: 30
