## Supplementary material for "Immunopeptidomics for Dummies: Detailed Experimental Protocols and Rapid, User-Friendly Visualization of MHC I and II Ligand Datasets with MhcVizPipe": All supplementary data and tables: S9_MVPReport Effect of acidic buffer EL4 sep G.htm

MhcVizPipe Report


# M

### hc

# V

### iz

# P

### ipe

### - Analysis report

---

**Date:** 2020-09-02

**Submitted by:** Anonymous

**Analysis type:** Class I

**Description of experiment:**

Effect of elution buffer

**Alleles:** H2-Kb

**Samples:**

0.2%\_Formic\_acid
10%\_acetic\_acid
1%\_TFA
10%\_TFA

**Species:** Mouse EL4

**# of cells:** 100 millions

**Lysis buffer:** chaps

**Type of beads:** sepharose G

**Antibody:** H2Kb (Y3)

**Incubation time:** overnight

**MHC-ligand complex elution buffer:** variable

**Peptide elution buffer:** 28% ACN

**Type of MS/MS:** Fusion

**Peptide identification software:** peaks

**Peptide FDR:** 1%

---

### Sample Overview

**UpSet Plot**

Only intersections > 0.5% are displayed

**Peptide Length Distribution** (maximum of 30 mers)

| Sample | Peptide length | Total peptides | % |
| --- | --- | --- | --- |
| 0.2%\_Formic\_acid | all lengths | 626 | 100 |
|  | 8-12 mers | 510 | 81 |
|  | other | 116 | 19 |
| 10%\_acetic\_acid | all lengths | 159 | 100 |
|  | 8-12 mers | 141 | 89 |
|  | other | 18 | 11 |
| 1%\_TFA | all lengths | 1057 | 100 |
|  | 8-12 mers | 963 | 91 |
|  | other | 94 | 9 |
| 10%\_TFA | all lengths | 967 | 100 |
|  | 8-12 mers | 747 | 77 |
|  | other | 220 | 23 |

---

### Annotation Results

NetMHCpan eluted ligand predictions made for all peptides between 8 & 12 mers, inclusive.
Percent rank cutoffs for strong and weak binders: 0.5 and 2.0.

| Allele | Sample | Total peptides | Strong binders | Weak binders |
| --- | --- | --- | --- | --- |
| H2-Kb | 0.2%\_Formic\_acid | 510 | 398 (78.0%) | 29 (5.7%) |
|  | 10%\_acetic\_acid | 141 | 130 (92.2%) | 5 (3.5%) |
|  | 1%\_TFA | 963 | 801 (83.2%) | 69 (7.2%) |
|  | 10%\_TFA | 747 | 542 (72.6%) | 42 (5.6%) |

**Binding Affinities**

---

### Binding Heatmaps

---

### Sequence Motifs

Clustering performed with all peptides between 8 & 12 mers, inclusive.

Percentages represent the percentage of peptides in a given group predicted to strongly bind the indicated allele.

Polar

Neutral

Basic

Acidic

Hydrophobic

- Unsupervised GibbsCluster
- Allele-specific GibbsCluster

**10%\_acetic\_acid**

Peptides in group: 2

H2-Kb: 0%

Peptides in group: 136

**H2-Kb: 96%**

**0.2%\_Formic\_acid**

Peptides in group: 468

**H2-Kb: 85%**

**1%\_TFA**

Peptides in group: 908

**H2-Kb: 88%**

**10%\_TFA**

Peptides in group: 626

**H2-Kb: 86%**

**10%\_acetic\_acid sequence motif(s)**

**H2-Kb**

Peptides: 135

**0.2%\_Formic\_acid sequence motif(s)**

**H2-Kb**

Peptides: 425

**Non-binders group 1**

Peptides: 19

**Non-binders group 2**

Peptides: 15

**Non-binders group 3**

Peptides: 11

**Non-binders group 4**

Peptides: 33

**1%\_TFA sequence motif(s)**

**H2-Kb**

Peptides: 862

**Non-binders group 1**

Peptides: 4

**Non-binders group 2**

Peptides: 19

**Non-binders group 3**

Peptides: 20

**Non-binders group 4**

Peptides: 42

**10%\_TFA sequence motif(s)**

**H2-Kb**

Peptides: 581

**Non-binders group 1**

Peptides: 28

**Non-binders group 2**

Peptides: 36

**Non-binders group 3**

Peptides: 16

**Non-binders group 4**

Peptides: 35

**Non-binders group 5**

Peptides: 38
