## Supplementary material for "Immunopeptidomics for Dummies: Detailed Experimental Protocols and Rapid, User-Friendly Visualization of MHC I and II Ligand Datasets with MhcVizPipe": All supplementary data and tables: S10_MVP report effect of acidic buffer on MHC class II mouse A20.htm

MhcVizPipe Report


# M

### hc

# V

### iz

# P

### ipe

### - Analysis report

---

**Date:** 2020-09-02

**Submitted by:** Anonymous

**Analysis type:** Class II

**Description of experiment:**

Effect of elution buffer

**Alleles:** H-2-IAd, H-2-IEd

**Samples:**

Formic\_Acid
Acetic\_Acid
1%\_TFA
10%\_TFA

**Species:** Mouse, A20 cell line

**# of cells:** 100 millions

**Lysis buffer:** chaps

**Type of beads:** Sepharose G

**Antibody:** M5

**Incubation time:** Overnight

**MHC-ligand complex elution buffer:** variable

**Peptide elution buffer:** ACN 28%

**Type of MS/MS:** Fusion

**Peptide identification software:** Peaks

**Peptide FDR:** 1%

---

### Sample Overview

**UpSet Plot**

Only intersections > 0.5% are displayed

**Peptide Length Distribution** (maximum of 30 mers)

| Sample | Peptide length | Total peptides | % |
| --- | --- | --- | --- |
| Formic\_Acid | all lengths | 563 | 100 |
|  | 9-22 mers | 474 | 84 |
|  | other | 89 | 16 |
| Acetic\_Acid | all lengths | 315 | 100 |
|  | 9-22 mers | 284 | 90 |
|  | other | 31 | 10 |
| 1%\_TFA | all lengths | 2019 | 100 |
|  | 9-22 mers | 1825 | 90 |
|  | other | 194 | 10 |
| 10%\_TFA | all lengths | 2139 | 100 |
|  | 9-22 mers | 1457 | 68 |
|  | other | 682 | 32 |

---

### Annotation Results

NetMHCIIpan eluted ligand predictions made for all peptides between 9 & 22 mers, inclusive.
Percent rank cutoffs for strong and weak binders: 2.0 and 10.0.

| Allele | Sample | Total peptides | Strong binders | Weak binders |
| --- | --- | --- | --- | --- |
| H-2-IAd | Formic\_Acid | 474 | 139 (29.3%) | 64 (13.5%) |
|  | Acetic\_Acid | 284 | 143 (50.4%) | 60 (21.1%) |
|  | 1%\_TFA | 1825 | 790 (43.3%) | 401 (22.0%) |
|  | 10%\_TFA | 1457 | 381 (26.1%) | 151 (10.4%) |
| H-2-IEd | Formic\_Acid | 474 | 50 (10.5%) | 89 (18.8%) |
|  | Acetic\_Acid | 284 | 19 (6.7%) | 55 (19.4%) |
|  | 1%\_TFA | 1825 | 243 (13.3%) | 354 (19.4%) |
|  | 10%\_TFA | 1457 | 186 (12.8%) | 210 (14.4%) |

**Binding Affinities**

---

### Binding Heatmaps

---

### Sequence Motifs

Clustering performed with all peptides between 9 & 22 mers, inclusive.

Percentages represent the percentage of peptides in a given group predicted to strongly bind the indicated allele.

Polar

Neutral

Basic

Acidic

Hydrophobic

- Unsupervised GibbsCluster
- Allele-specific GibbsCluster

**Acetic\_Acid**

Peptides in group: 53

H-2-IAd: 8%,

**H-2-IEd: 23%**

Peptides in group: 224

**H-2-IAd: 62%,** 

H-2-IEd: 3%

**1%\_TFA**

Peptides in group: 449

H-2-IAd: 2%,

**H-2-IEd: 44%**

Peptides in group: 1339

**H-2-IAd: 58%,** 

H-2-IEd: 3%

**10%\_TFA**

Peptides in group: 561

H-2-IAd: 2%,

**H-2-IEd: 23%**

Peptides in group: 692

**H-2-IAd: 53%,** 

H-2-IEd: 8%

**Formic\_Acid**

Peptides in group: 365

**H-2-IAd: 37%,** 

H-2-IEd: 13%

**Acetic\_Acid sequence motif(s)**

**H-2-IAd**

Peptides: 203

**H-2-IEd**

Peptides: 74

**Non-binders group 1**

Peptides: 30

**Non-binders group 2**

Peptides: 17

**1%\_TFA sequence motif(s)**

**H-2-IAd**

Peptides: 1185

**H-2-IEd**

Peptides: 570

**Non-binders group 1**

Peptides: 100

**Non-binders group 2**

Peptides: 206

**10%\_TFA sequence motif(s)**

**H-2-IAd**

Peptides: 528

**H-2-IEd**

Peptides: 389

**Non-binders group 1**

Peptides: 259

**Non-binders group 2**

Peptides: 320

**Formic\_Acid sequence motif(s)**

**H-2-IAd**

Peptides: 201

**H-2-IEd**

Peptides: 137

**Non-binders group 1**

Peptides: 84

**Non-binders group 2**

Peptides: 39

**Non-binders group 3**

Peptides: 54
