## Supplementary material for "Immunopeptidomics for Dummies: Detailed Experimental Protocols and Rapid, User-Friendly Visualization of MHC I and II Ligand Datasets with MhcVizPipe": All supplementary data and tables: S11_MVP Report H2-Db_canonical.html

MhcVizPipe Report


# M

### hc

# V

### iz

# P

### ipe

### - Analysis report

---

**Date:** 2020-09-03

**Submitted by:** Anonymous

**Analysis type:** Class I

**Description of experiment:**

None provided

**Alleles:** H-2-Db

**Samples:**

AdrenalGland.txt: AdrenalGland.txt
Bladder.txt: Bladder.txt
BoneMarrow.txt: BoneMarrow.txt
Brain.txt: Brain.txt
Colon.txt: Colon.txt
Heart.txt: Heart.txt
Kidney.txt: Kidney.txt
Liver.txt: Liver.txt
Lung.txt: Lung.txt
Ovary.txt: Ovary.txt
Pancreas.txt: Pancreas.txt
Skin.txt: Skin.txt
SpinalCord.txt: SpinalCord.txt
Spleen.txt: Spleen.txt
Stomach.txt: Stomach.txt
Testis.txt: Testis.txt
Thymus.txt: Thymus.txt
Uterus.txt: Uterus.txt
Intestine.txt: Intestine.txt

---

### Sample Overview

**UpSet Plot**

Only intersections > 0.5% are displayed

**Peptide Length Distribution** (maximum of 30 mers)

| Sample | Peptide length | Total peptides | % |
| --- | --- | --- | --- |
| AdrenalGland.txt | all lengths | 758 | 100 |
|  | 8-12 mers | 728 | 96 |
|  | other | 30 | 4 |
| Bladder.txt | all lengths | 968 | 100 |
|  | 8-12 mers | 867 | 90 |
|  | other | 101 | 10 |
| BoneMarrow.txt | all lengths | 1492 | 100 |
|  | 8-12 mers | 1438 | 96 |
|  | other | 54 | 4 |
| Brain.txt | all lengths | 1098 | 100 |
|  | 8-12 mers | 942 | 86 |
|  | other | 156 | 14 |
| Colon.txt | all lengths | 2998 | 100 |
|  | 8-12 mers | 2585 | 86 |
|  | other | 413 | 14 |
| Heart.txt | all lengths | 778 | 100 |
|  | 8-12 mers | 693 | 89 |
|  | other | 85 | 11 |
| Kidney.txt | all lengths | 3821 | 100 |
|  | 8-12 mers | 3218 | 84 |
|  | other | 603 | 16 |
| Liver.txt | all lengths | 3561 | 100 |
|  | 8-12 mers | 3246 | 91 |
|  | other | 315 | 9 |
| Lung.txt | all lengths | 2547 | 100 |
|  | 8-12 mers | 2399 | 94 |
|  | other | 148 | 6 |
| Ovary.txt | all lengths | 635 | 100 |
|  | 8-12 mers | 575 | 91 |
|  | other | 60 | 9 |
| Pancreas.txt | all lengths | 382 | 100 |
|  | 8-12 mers | 341 | 89 |
|  | other | 41 | 11 |
| Skin.txt | all lengths | 1255 | 100 |
|  | 8-12 mers | 1208 | 96 |
|  | other | 47 | 4 |
| SpinalCord.txt | all lengths | 512 | 100 |
|  | 8-12 mers | 433 | 85 |
|  | other | 79 | 15 |
| Spleen.txt | all lengths | 3219 | 100 |
|  | 8-12 mers | 3058 | 95 |
|  | other | 161 | 5 |
| Stomach.txt | all lengths | 5625 | 100 |
|  | 8-12 mers | 4169 | 74 |
|  | other | 1456 | 26 |
| Testis.txt | all lengths | 1045 | 100 |
|  | 8-12 mers | 971 | 93 |
|  | other | 74 | 7 |
| Thymus.txt | all lengths | 2362 | 100 |
|  | 8-12 mers | 2220 | 94 |
|  | other | 142 | 6 |
| Uterus.txt | all lengths | 1486 | 100 |
|  | 8-12 mers | 1437 | 97 |
|  | other | 49 | 3 |
| Intestine.txt | all lengths | 6203 | 100 |
|  | 8-12 mers | 5117 | 82 |
|  | other | 1086 | 18 |

---

### Annotation Results

NetMHCpan eluted ligand predictions made for all peptides between 8 & 12 mers, inclusive.
Percent rank cutoffs for strong and weak binders: 0.5 and 2.0.

| Allele | Sample | Total peptides | Strong binders | Weak binders |
| --- | --- | --- | --- | --- |
| H-2-Db | AdrenalGland.txt | 728 | 501 (68.8%) | 29 (4.0%) |
|  | Bladder.txt | 867 | 504 (58.1%) | 36 (4.2%) |
|  | BoneMarrow.txt | 1438 | 1006 (70.0%) | 99 (6.9%) |
|  | Brain.txt | 942 | 427 (45.3%) | 38 (4.0%) |
|  | Colon.txt | 2585 | 1260 (48.7%) | 143 (5.5%) |
|  | Heart.txt | 693 | 432 (62.3%) | 17 (2.5%) |
|  | Kidney.txt | 3218 | 1301 (40.4%) | 163 (5.1%) |
|  | Liver.txt | 3246 | 1758 (54.2%) | 269 (8.3%) |
|  | Lung.txt | 2399 | 1534 (63.9%) | 169 (7.0%) |
|  | Ovary.txt | 575 | 364 (63.3%) | 19 (3.3%) |
|  | Pancreas.txt | 341 | 217 (63.6%) | 8 (2.3%) |
|  | Skin.txt | 1208 | 914 (75.7%) | 89 (7.4%) |
|  | SpinalCord.txt | 433 | 141 (32.6%) | 15 (3.5%) |
|  | Spleen.txt | 3058 | 2044 (66.8%) | 241 (7.9%) |
|  | Stomach.txt | 4169 | 347 (8.3%) | 142 (3.4%) |
|  | Testis.txt | 971 | 660 (68.0%) | 42 (4.3%) |
|  | Thymus.txt | 2220 | 1452 (65.4%) | 140 (6.3%) |
|  | Uterus.txt | 1437 | 1078 (75.0%) | 76 (5.3%) |
|  | Intestine.txt | 5117 | 1673 (32.7%) | 270 (5.3%) |

**Binding Affinities**

---

### Binding Heatmaps

---

### Sequence Motifs

Clustering performed with all peptides between 8 & 12 mers, inclusive.

Percentages represent the percentage of peptides in a given group predicted to strongly bind the indicated allele.

Polar

Neutral

Basic

Acidic

Hydrophobic

- Unsupervised GibbsCluster
- Allele-specific GibbsCluster

**AdrenalGland.txt**

Peptides in group: 600

**H-2-Db: 83%**

**Bladder.txt**

Peptides in group: 667

**H-2-Db: 75%**

**BoneMarrow.txt**

Peptides in group: 1275

**H-2-Db: 79%**

**Brain.txt**

Peptides in group: 668

**H-2-Db: 64%**

**Colon.txt**

Peptides in group: 1880

**H-2-Db: 67%**

**Heart.txt**

Peptides in group: 538

**H-2-Db: 80%**

**Kidney.txt**

Peptides in group: 2204

**H-2-Db: 59%**

**Liver.txt**

Peptides in group: 2547

**H-2-Db: 69%**

**Lung.txt**

Peptides in group: 1936

**H-2-Db: 79%**

**Ovary.txt**

Peptides in group: 437

**H-2-Db: 83%**

**Pancreas.txt**

Peptides in group: 284

**H-2-Db: 76%**

**Skin.txt**

Peptides in group: 1088

**H-2-Db: 84%**

**SpinalCord.txt**

Peptides in group: 173

**H-2-Db: 81%**

Peptides in group: 69

H-2-Db: 0%

Peptides in group: 46

H-2-Db: 0%

Peptides in group: 71

**H-2-Db: 1%**

Peptides in group: 54

H-2-Db: 0%

**Spleen.txt**

Peptides in group: 2615

**H-2-Db: 78%**

**Stomach.txt**

Peptides in group: 692

**H-2-Db: 1%**

Peptides in group: 906

H-2-Db: 0%

Peptides in group: 727

**H-2-Db: 2%**

Peptides in group: 818

**H-2-Db: 39%**

Peptides in group: 736

H-2-Db: 0%

**Testis.txt**

Peptides in group: 797

**H-2-Db: 83%**

**Thymus.txt**

Peptides in group: 1790

**H-2-Db: 81%**

**Uterus.txt**

Peptides in group: 1249

**H-2-Db: 86%**

**Intestine.txt**

Peptides in group: 2289

**H-2-Db: 72%**

Peptides in group: 1068

**H-2-Db: 1%**

Peptides in group: 1215

H-2-Db: 0%

**AdrenalGland.txt sequence motif(s)**

**H-2-Db**

Peptides: 519

**Non-binders group 3**

Peptides: 23

**Non-binders group 1**

Peptides: 25

**Non-binders group 5**

Peptides: 47

**Non-binders group 2**

Peptides: 44

**Non-binders group 4**

Peptides: 45

**Bladder.txt sequence motif(s)**

**H-2-Db**

Peptides: 524

**Non-binders group 3**

Peptides: 51

**Non-binders group 1**

Peptides: 73

**Non-binders group 5**

Peptides: 66

**Non-binders group 2**

Peptides: 57

**Non-binders group 4**

Peptides: 63

**BoneMarrow.txt sequence motif(s)**

**H-2-Db**

Peptides: 1089

**Non-binders group 2**

Peptides: 104

**Non-binders group 3**

Peptides: 88

**Non-binders group 4**

Peptides: 50

**Non-binders group 1**

Peptides: 75

**Brain.txt sequence motif(s)**

**H-2-Db**

Peptides: 455

**Non-binders group 3**

Peptides: 80

**Non-binders group 1**

Peptides: 87

**Non-binders group 5**

Peptides: 101

**Non-binders group 2**

Peptides: 96

**Non-binders group 4**

Peptides: 82

**Colon.txt sequence motif(s)**

**H-2-Db**

Peptides: 1371

**Non-binders group 3**

Peptides: 212

**Non-binders group 1**

Peptides: 218

**Non-binders group 5**

Peptides: 193

**Non-binders group 2**

Peptides: 246

**Non-binders group 4**

Peptides: 248

**Heart.txt sequence motif(s)**

**H-2-Db**

Peptides: 443

**Non-binders group 3**

Peptides: 35

**Non-binders group 1**

Peptides: 48

**Non-binders group 5**

Peptides: 64

**Non-binders group 2**

Peptides: 31

**Non-binders group 4**

Peptides: 47

**Kidney.txt sequence motif(s)**

**H-2-Db**

Peptides: 1423

**Non-binders group 3**

Peptides: 305

**Non-binders group 1**

Peptides: 301

**Non-binders group 5**

Peptides: 372

**Non-binders group 2**

Peptides: 314

**Non-binders group 4**

Peptides: 330

**Liver.txt sequence motif(s)**

**H-2-Db**

Peptides: 1967

**Non-binders group 3**

Peptides: 240

**Non-binders group 1**

Peptides: 227

**Non-binders group 5**

Peptides: 274

**Non-binders group 2**

Peptides: 211

**Non-binders group 4**

Peptides: 174

**Lung.txt sequence motif(s)**

**H-2-Db**

Peptides: 1655

**Non-binders group 2**

Peptides: 182

**Non-binders group 3**

Peptides: 119

**Non-binders group 4**

Peptides: 148

**Non-binders group 1**

Peptides: 172

**Ovary.txt sequence motif(s)**

**H-2-Db**

Peptides: 379

**Non-binders group 3**

Peptides: 30

**Non-binders group 1**

Peptides: 40

**Non-binders group 5**

Peptides: 52

**Non-binders group 2**

Peptides: 30

**Non-binders group 4**

Peptides: 33

**Pancreas.txt sequence motif(s)**

**H-2-Db**

Peptides: 223

**Non-binders group 3**

Peptides: 20

**Non-binders group 1**

Peptides: 13

**Non-binders group 5**

Peptides: 23

**Non-binders group 2**

Peptides: 23

**Non-binders group 4**

Peptides: 30

**Skin.txt sequence motif(s)**

**H-2-Db**

Peptides: 987

**Non-binders group 2**

Peptides: 50

**Non-binders group 1**

Peptides: 52

**Non-binders group 3**

Peptides: 75

**SpinalCord.txt sequence motif(s)**

**H-2-Db**

Peptides: 152

**Non-binders group 3**

Peptides: 47

**Non-binders group 1**

Peptides: 53

**Non-binders group 5**

Peptides: 39

**Non-binders group 2**

Peptides: 44

**Non-binders group 4**

Peptides: 81

**Spleen.txt sequence motif(s)**

**H-2-Db**

Peptides: 2210

**Non-binders group 3**

Peptides: 103

**Non-binders group 1**

Peptides: 114

**Non-binders group 5**

Peptides: 221

**Non-binders group 2**

Peptides: 113

**Non-binders group 4**

Peptides: 179

**Stomach.txt sequence motif(s)**

**H-2-Db**

Peptides: 468

**Non-binders group 3**

Peptides: 685

**Non-binders group 1**

Peptides: 685

**Non-binders group 5**

Peptides: 692

**Non-binders group 2**

Peptides: 667

**Non-binders group 4**

Peptides: 698

**Testis.txt sequence motif(s)**

**H-2-Db**

Peptides: 692

**Non-binders group 3**

Peptides: 53

**Non-binders group 1**

Peptides: 60

**Non-binders group 5**

Peptides: 48

**Non-binders group 2**

Peptides: 46

**Non-binders group 4**

Peptides: 44

**Thymus.txt sequence motif(s)**

**H-2-Db**

Peptides: 1550

**Non-binders group 2**

Peptides: 148

**Non-binders group 3**

Peptides: 115

**Non-binders group 4**

Peptides: 152

**Non-binders group 1**

Peptides: 151

**Uterus.txt sequence motif(s)**

**H-2-Db**

Peptides: 1135

**Non-binders group 3**

Peptides: 71

**Non-binders group 1**

Peptides: 53

**Non-binders group 5**

Peptides: 41

**Non-binders group 2**

Peptides: 62

**Non-binders group 4**

Peptides: 43

**Intestine.txt sequence motif(s)**

**H-2-Db**

Peptides: 1871

**Non-binders group 3**

Peptides: 564

**Non-binders group 1**

Peptides: 522

**Non-binders group 5**

Peptides: 572

**Non-binders group 2**

Peptides: 667

**Non-binders group 4**

Peptides: 587
