## Supplementary material for "Immunopeptidomics for Dummies: Detailed Experimental Protocols and Rapid, User-Friendly Visualization of MHC I and II Ligand Datasets with MhcVizPipe": All supplementary data and tables: S12_MVP report H2Db_PSCP.html

MhcVizPipe Report


# M

### hc

# V

### iz

# P

### ipe

### - Analysis report

---

**Date:** 2020-09-02

**Submitted by:** Anonymous

**Analysis type:** Class I

**Description of experiment:**

Spliced H2-Db

**Alleles:** H-2-Db

**Samples:**

ADRENAL.txt: ADRENAL.txt
BLADDER.txt: BLADDER.txt
BONE\_MARROW.txt: BONE\_MARROW.txt
BRAIN.txt: BRAIN.txt
COLON.txt: COLON.txt
heart.txt: heart.txt
kidney.txt: kidney.txt
Liver.txt: Liver.txt
lung.txt: lung.txt
ovary.txt: ovary.txt
pancreas.txt: pancreas.txt
skin.txt: skin.txt
small\_intestine.txt: small\_intestine.txt
spinal\_cord.txt: spinal\_cord.txt
spleen.txt: spleen.txt
stomach.txt: stomach.txt
testis.txt: testis.txt
Thymus.txt: Thymus.txt
uterus.txt: uterus.txt

---

### Sample Overview

**UpSet Plot**

Only intersections > 0.5% are displayed

**Peptide Length Distribution** (maximum of 30 mers)

| Sample | Peptide length | Total peptides | % |
| --- | --- | --- | --- |
| ADRENAL.txt | all lengths | 93 | 100 |
|  | 8-12 mers | 93 | 100 |
|  | other | 0 | 0 |
| BLADDER.txt | all lengths | 57 | 100 |
|  | 8-12 mers | 57 | 100 |
|  | other | 0 | 0 |
| BONE\_MARROW.txt | all lengths | 154 | 100 |
|  | 8-12 mers | 154 | 100 |
|  | other | 0 | 0 |
| BRAIN.txt | all lengths | 136 | 100 |
|  | 8-12 mers | 136 | 100 |
|  | other | 0 | 0 |
| COLON.txt | all lengths | 271 | 100 |
|  | 8-12 mers | 271 | 100 |
|  | other | 0 | 0 |
| heart.txt | all lengths | 39 | 100 |
|  | 8-12 mers | 39 | 100 |
|  | other | 0 | 0 |
| kidney.txt | all lengths | 314 | 100 |
|  | 8-12 mers | 314 | 100 |
|  | other | 0 | 0 |
| Liver.txt | all lengths | 349 | 100 |
|  | 8-12 mers | 349 | 100 |
|  | other | 0 | 0 |
| lung.txt | all lengths | 442 | 100 |
|  | 8-12 mers | 442 | 100 |
|  | other | 0 | 0 |
| ovary.txt | all lengths | 33 | 100 |
|  | 8-12 mers | 33 | 100 |
|  | other | 0 | 0 |
| pancreas.txt | all lengths | 25 | 100 |
|  | 8-12 mers | 25 | 100 |
|  | other | 0 | 0 |
| skin.txt | all lengths | 134 | 100 |
|  | 8-12 mers | 134 | 100 |
|  | other | 0 | 0 |
| small\_intestine.txt | all lengths | 362 | 100 |
|  | 8-12 mers | 362 | 100 |
|  | other | 0 | 0 |
| spinal\_cord.txt | all lengths | 34 | 100 |
|  | 8-12 mers | 34 | 100 |
|  | other | 0 | 0 |
| spleen.txt | all lengths | 337 | 100 |
|  | 8-12 mers | 337 | 100 |
|  | other | 0 | 0 |
| stomach.txt | all lengths | 74 | 100 |
|  | 8-12 mers | 74 | 100 |
|  | other | 0 | 0 |
| testis.txt | all lengths | 72 | 100 |
|  | 8-12 mers | 72 | 100 |
|  | other | 0 | 0 |
| Thymus.txt | all lengths | 231 | 100 |
|  | 8-12 mers | 231 | 100 |
|  | other | 0 | 0 |
| uterus.txt | all lengths | 175 | 100 |
|  | 8-12 mers | 175 | 100 |
|  | other | 0 | 0 |

---

### Annotation Results

NetMHCpan eluted ligand predictions made for all peptides between 8 & 12 mers, inclusive.
Percent rank cutoffs for strong and weak binders: 0.5 and 2.0.

| Allele | Sample | Total peptides | Strong binders | Weak binders |
| --- | --- | --- | --- | --- |
| H-2-Db | ADRENAL.txt | 93 | 20 (21.5%) | 4 (4.3%) |
|  | BLADDER.txt | 57 | 21 (36.8%) | 1 (1.8%) |
|  | BONE\_MARROW.txt | 154 | 70 (45.5%) | 14 (9.1%) |
|  | BRAIN.txt | 136 | 15 (11.0%) | 2 (1.5%) |
|  | COLON.txt | 271 | 91 (33.6%) | 6 (2.2%) |
|  | heart.txt | 39 | 15 (38.5%) | 0 (0.0%) |
|  | kidney.txt | 314 | 103 (32.8%) | 17 (5.4%) |
|  | Liver.txt | 349 | 214 (61.3%) | 15 (4.3%) |
|  | lung.txt | 442 | 211 (47.7%) | 39 (8.8%) |
|  | ovary.txt | 33 | 4 (12.1%) | 1 (3.0%) |
|  | pancreas.txt | 25 | 8 (32.0%) | 1 (4.0%) |
|  | skin.txt | 134 | 66 (49.3%) | 13 (9.7%) |
|  | small\_intestine.txt | 362 | 150 (41.4%) | 23 (6.4%) |
|  | spinal\_cord.txt | 34 | 2 (5.9%) | 0 (0.0%) |
|  | spleen.txt | 337 | 215 (63.8%) | 38 (11.3%) |
|  | stomach.txt | 74 | 6 (8.1%) | 0 (0.0%) |
|  | testis.txt | 72 | 17 (23.6%) | 1 (1.4%) |
|  | Thymus.txt | 231 | 137 (59.3%) | 18 (7.8%) |
|  | uterus.txt | 175 | 78 (44.6%) | 12 (6.9%) |

**Binding Affinities**

---

### Binding Heatmaps

---

### Sequence Motifs

Clustering performed with all peptides between 8 & 12 mers, inclusive.

Percentages represent the percentage of peptides in a given group predicted to strongly bind the indicated allele.

Polar

Neutral

Basic

Acidic

Hydrophobic

- Unsupervised GibbsCluster
- Allele-specific GibbsCluster

**COLON.txt**

Peptides in group: 129

**H-2-Db: 69%**

Peptides in group: 46

H-2-Db: 0%

Peptides in group: 11

H-2-Db: 0%

Peptides in group: 53

H-2-Db: 0%

Peptides in group: 32

**H-2-Db: 6%**

**lung.txt**

Peptides in group: 276

**H-2-Db: 76%**

Peptides in group: 71

H-2-Db: 0%

Peptides in group: 37

H-2-Db: 0%

Peptides in group: 15

H-2-Db: 0%

Peptides in group: 43

**H-2-Db: 2%**

**spleen.txt**

Peptides in group: 280

**H-2-Db: 75%**

Peptides in group: 32

H-2-Db: 0%

Peptides in group: 14

**H-2-Db: 36%**

Peptides in group: 9

H-2-Db: 0%

**testis.txt**

Peptides in group: 22

**H-2-Db: 68%**

Peptides in group: 19

H-2-Db: 0%

Peptides in group: 24

**H-2-Db: 4%**

Peptides in group: 3

**H-2-Db: 33%**

**ADRENAL.txt**

Peptides in group: 26

**H-2-Db: 73%**

Peptides in group: 60

H-2-Db: 0%

Peptides in group: 5

**H-2-Db: 20%**

**BONE\_MARROW.txt**

Peptides in group: 95

**H-2-Db: 69%**

Peptides in group: 36

H-2-Db: 0%

Peptides in group: 19

**H-2-Db: 21%**

**BRAIN.txt**

Peptides in group: 25

**H-2-Db: 28%**

Peptides in group: 54

**H-2-Db: 13%**

Peptides in group: 53

**H-2-Db: 2%**

**kidney.txt**

Peptides in group: 168

**H-2-Db: 61%**

Peptides in group: 65

H-2-Db: 0%

Peptides in group: 66

H-2-Db: 0%

**Liver.txt**

Peptides in group: 257

**H-2-Db: 83%**

Peptides in group: 40

H-2-Db: 0%

Peptides in group: 41

**H-2-Db: 2%**

**ovary.txt**

Peptides in group: 7

H-2-Db: 0%

Peptides in group: 4

**H-2-Db: 25%**

Peptides in group: 19

**H-2-Db: 16%**

**skin.txt**

Peptides in group: 91

**H-2-Db: 73%**

Peptides in group: 17

H-2-Db: 0%

Peptides in group: 24

H-2-Db: 0%

**small\_intestine.txt**

Peptides in group: 234

**H-2-Db: 64%**

Peptides in group: 44

H-2-Db: 0%

Peptides in group: 77

H-2-Db: 0%

**spinal\_cord.txt**

Peptides in group: 15

**H-2-Db: 7%**

Peptides in group: 10

**H-2-Db: 10%**

Peptides in group: 8

H-2-Db: 0%

**stomach.txt**

Peptides in group: 10

**H-2-Db: 40%**

Peptides in group: 19

H-2-Db: 0%

Peptides in group: 40

**H-2-Db: 2%**

**Thymus.txt**

Peptides in group: 170

**H-2-Db: 80%**

Peptides in group: 47

**H-2-Db: 2%**

Peptides in group: 10

H-2-Db: 0%

**uterus.txt**

Peptides in group: 101

**H-2-Db: 71%**

Peptides in group: 60

H-2-Db: 0%

Peptides in group: 13

**H-2-Db: 46%**

**BLADDER.txt**

Peptides in group: 32

**H-2-Db: 66%**

Peptides in group: 21

H-2-Db: 0%

**heart.txt**

Peptides in group: 17

**H-2-Db: 88%**

Peptides in group: 19

H-2-Db: 0%

**pancreas.txt**

Peptides in group: 24

**H-2-Db: 33%**

**COLON.txt sequence motif(s)**

**H-2-Db**

Peptides: 97

**Non-binders group 1**

Peptides: 37

**Non-binders group 2**

Peptides: 33

**Non-binders group 3**

Peptides: 54

**Non-binders group 4**

Peptides: 22

**Non-binders group 5**

Peptides: 27

**lung.txt sequence motif(s)**

**H-2-Db**

Peptides: 250

**Non-binders group 1**

Peptides: 64

**Non-binders group 2**

Peptides: 40

**Non-binders group 3**

Peptides: 6

**Non-binders group 4**

Peptides: 45

**Non-binders group 5**

Peptides: 34

**spleen.txt sequence motif(s)**

**H-2-Db**

Peptides: 253

**Non-binders group 1**

Peptides: 23

**Non-binders group 2**

Peptides: 15

**Non-binders group 3**

Peptides: 34

**Non-binders group 4**

Peptides: 12

**testis.txt sequence motif(s)**

**H-2-Db**

Too few peptides to cluster

**Non-binders group 1**

Peptides: 4

**Non-binders group 2**

Peptides: 4

**Non-binders group 3**

Peptides: 27

**ADRENAL.txt sequence motif(s)**

**H-2-Db**

Peptides: 24

**Non-binders group 1**

Peptides: 18

**Non-binders group 2**

Peptides: 51

**BONE\_MARROW.txt sequence motif(s)**

**H-2-Db**

Peptides: 84

**Non-binders group 1**

Peptides: 28

**Non-binders group 2**

Peptides: 10

**Non-binders group 3**

Peptides: 27

**BRAIN.txt sequence motif(s)**

**H-2-Db**

Too few peptides to cluster

**Non-binders group 1**

Peptides: 40

**Non-binders group 2**

Peptides: 17

**Non-binders group 3**

Peptides: 13

**Non-binders group 4**

Peptides: 6

**Non-binders group 5**

Peptides: 41

**kidney.txt sequence motif(s)**

**H-2-Db**

Peptides: 119

**Non-binders group 1**

Peptides: 33

**Non-binders group 2**

Peptides: 61

**Non-binders group 3**

Peptides: 67

**Non-binders group 4**

Peptides: 27

**Liver.txt sequence motif(s)**

**H-2-Db**

Peptides: 228

**Non-binders group 1**

Peptides: 10

**Non-binders group 2**

Peptides: 49

**Non-binders group 3**

Peptides: 41

**Non-binders group 4**

Peptides: 18

**ovary.txt sequence motif(s)**

**H-2-Db**

Too few peptides to cluster

**Non-binders group 1**

Peptides: 12

**skin.txt sequence motif(s)**

**H-2-Db**

Peptides: 79

**Non-binders group 1**

Peptides: 26

**Non-binders group 2**

Peptides: 10

**Non-binders group 3**

Peptides: 6

**Non-binders group 4**

Peptides: 12

**small\_intestine.txt sequence motif(s)**

**H-2-Db**

Peptides: 172

**Non-binders group 1**

Peptides: 63

**Non-binders group 2**

Peptides: 34

**Non-binders group 3**

Peptides: 41

**Non-binders group 4**

Peptides: 6

**Non-binders group 5**

Peptides: 43

**spinal\_cord.txt sequence motif(s)**

**H-2-Db**

Too few peptides to cluster

**Non-binders group 1**

Peptides: 13

**Non-binders group 2**

Peptides: 18

**stomach.txt sequence motif(s)**

**H-2-Db**

Too few peptides to cluster

**Non-binders group 1**

Peptides: 36

**Non-binders group 2**

Peptides: 25

**Thymus.txt sequence motif(s)**

**H-2-Db**

Peptides: 154

**Non-binders group 1**

Peptides: 38

**Non-binders group 2**

Peptides: 17

**Non-binders group 3**

Peptides: 14

**Non-binders group 4**

Peptides: 6

**uterus.txt sequence motif(s)**

**H-2-Db**

Peptides: 90

**Non-binders group 1**

Peptides: 13

**Non-binders group 2**

Peptides: 20

**Non-binders group 3**

Peptides: 7

**Non-binders group 4**

Peptides: 30

**Non-binders group 5**

Peptides: 13

**BLADDER.txt sequence motif(s)**

**H-2-Db**

Peptides: 22

**Non-binders group 1**

Peptides: 4

**Non-binders group 2**

Peptides: 9

**Non-binders group 3**

Peptides: 18

**heart.txt sequence motif(s)**

**H-2-Db**

Too few peptides to cluster

**Non-binders group 1**

Peptides: 19

**pancreas.txt sequence motif(s)**

**H-2-Db**

Too few peptides to cluster
