## Supplementary material for "Immunopeptidomics for Dummies: Detailed Experimental Protocols and Rapid, User-Friendly Visualization of MHC I and II Ligand Datasets with MhcVizPipe": All supplementary data and tables: S13_MVP report H2-Kb_canonical.html

MhcVizPipe Report


# M

### hc

# V

### iz

# P

### ipe

### - Analysis report

---

**Date:** 2020-09-03

**Submitted by:** Anonymous

**Analysis type:** Class I

**Description of experiment:**

Canonical H2-Kb

**Alleles:** H-2-Kb

**Samples:**

AdrenalGland.txt: AdrenalGland.txt
Bladder.txt: Bladder.txt
BoneMarrow.txt: BoneMarrow.txt
Brain.txt: Brain.txt
Colon.txt: Colon.txt
Heart.txt: Heart.txt
Kidney.txt: Kidney.txt
Liver.txt: Liver.txt
Lung.txt: Lung.txt
Ovary.txt: Ovary.txt
Pancreas.txt: Pancreas.txt
Skin.txt: Skin.txt
SpinalCord.txt: SpinalCord.txt
Spleen.txt: Spleen.txt
Testis.txt: Testis.txt
Thymus.txt: Thymus.txt
Uterus.txt: Uterus.txt
Intestine.txt: Intestine.txt
Stomach.txt: Stomach.txt

---

### Sample Overview

**UpSet Plot**

Only intersections > 0.5% are displayed

**Peptide Length Distribution** (maximum of 30 mers)

| Sample | Peptide length | Total peptides | % |
| --- | --- | --- | --- |
| AdrenalGland.txt | all lengths | 545 | 100 |
|  | 8-12 mers | 505 | 93 |
|  | other | 40 | 7 |
| Bladder.txt | all lengths | 783 | 100 |
|  | 8-12 mers | 704 | 90 |
|  | other | 79 | 10 |
| BoneMarrow.txt | all lengths | 1117 | 100 |
|  | 8-12 mers | 1051 | 94 |
|  | other | 66 | 6 |
| Brain.txt | all lengths | 1277 | 100 |
|  | 8-12 mers | 975 | 76 |
|  | other | 302 | 24 |
| Colon.txt | all lengths | 3854 | 100 |
|  | 8-12 mers | 3118 | 81 |
|  | other | 736 | 19 |
| Heart.txt | all lengths | 735 | 100 |
|  | 8-12 mers | 617 | 84 |
|  | other | 118 | 16 |
| Kidney.txt | all lengths | 4096 | 100 |
|  | 8-12 mers | 3295 | 80 |
|  | other | 801 | 20 |
| Liver.txt | all lengths | 3860 | 100 |
|  | 8-12 mers | 3429 | 89 |
|  | other | 431 | 11 |
| Lung.txt | all lengths | 2005 | 100 |
|  | 8-12 mers | 1819 | 91 |
|  | other | 186 | 9 |
| Ovary.txt | all lengths | 370 | 100 |
|  | 8-12 mers | 321 | 87 |
|  | other | 49 | 13 |
| Pancreas.txt | all lengths | 316 | 100 |
|  | 8-12 mers | 276 | 87 |
|  | other | 40 | 13 |
| Skin.txt | all lengths | 756 | 100 |
|  | 8-12 mers | 716 | 95 |
|  | other | 40 | 5 |
| SpinalCord.txt | all lengths | 114 | 100 |
|  | 8-12 mers | 84 | 74 |
|  | other | 30 | 26 |
| Spleen.txt | all lengths | 2430 | 100 |
|  | 8-12 mers | 2287 | 94 |
|  | other | 143 | 6 |
| Testis.txt | all lengths | 1092 | 100 |
|  | 8-12 mers | 953 | 87 |
|  | other | 139 | 13 |
| Thymus.txt | all lengths | 1694 | 100 |
|  | 8-12 mers | 1574 | 93 |
|  | other | 120 | 7 |
| Uterus.txt | all lengths | 1739 | 100 |
|  | 8-12 mers | 1613 | 93 |
|  | other | 126 | 7 |
| Intestine.txt | all lengths | 7532 | 100 |
|  | 8-12 mers | 5763 | 77 |
|  | other | 1769 | 23 |
| Stomach.txt | all lengths | 5944 | 100 |
|  | 8-12 mers | 4318 | 73 |
|  | other | 1626 | 27 |

---

### Annotation Results

NetMHCpan eluted ligand predictions made for all peptides between 8 & 12 mers, inclusive.
Percent rank cutoffs for strong and weak binders: 0.5 and 2.0.

| Allele | Sample | Total peptides | Strong binders | Weak binders |
| --- | --- | --- | --- | --- |
| H-2-Kb | AdrenalGland.txt | 505 | 342 (67.7%) | 19 (3.8%) |
|  | Bladder.txt | 704 | 396 (56.2%) | 39 (5.5%) |
|  | BoneMarrow.txt | 1051 | 755 (71.8%) | 80 (7.6%) |
|  | Brain.txt | 975 | 191 (19.6%) | 38 (3.9%) |
|  | Colon.txt | 3118 | 1218 (39.1%) | 176 (5.6%) |
|  | Heart.txt | 617 | 341 (55.3%) | 17 (2.8%) |
|  | Kidney.txt | 3295 | 986 (29.9%) | 138 (4.2%) |
|  | Liver.txt | 3429 | 1905 (55.6%) | 320 (9.3%) |
|  | Lung.txt | 1819 | 1091 (60.0%) | 111 (6.1%) |
|  | Ovary.txt | 321 | 136 (42.4%) | 17 (5.3%) |
|  | Pancreas.txt | 276 | 141 (51.1%) | 15 (5.4%) |
|  | Skin.txt | 716 | 526 (73.5%) | 44 (6.1%) |
|  | SpinalCord.txt | 84 | 11 (13.1%) | 4 (4.8%) |
|  | Spleen.txt | 2287 | 1651 (72.2%) | 165 (7.2%) |
|  | Testis.txt | 953 | 558 (58.6%) | 45 (4.7%) |
|  | Thymus.txt | 1574 | 1076 (68.4%) | 82 (5.2%) |
|  | Uterus.txt | 1613 | 1076 (66.7%) | 108 (6.7%) |
|  | Intestine.txt | 5763 | 1379 (23.9%) | 254 (4.4%) |
|  | Stomach.txt | 4318 | 260 (6.0%) | 176 (4.1%) |

**Binding Affinities**

---

### Binding Heatmaps

---

### Sequence Motifs

Clustering performed with all peptides between 8 & 12 mers, inclusive.

Percentages represent the percentage of peptides in a given group predicted to strongly bind the indicated allele.

Polar

Neutral

Basic

Acidic

Hydrophobic

- Unsupervised GibbsCluster
- Allele-specific GibbsCluster

**AdrenalGland.txt**

Peptides in group: 423

**H-2-Kb: 80%**

**Bladder.txt**

Peptides in group: 535

**H-2-Kb: 74%**

**BoneMarrow.txt**

Peptides in group: 912

**H-2-Kb: 82%**

**Brain.txt**

Peptides in group: 210

H-2-Kb: 0%

Peptides in group: 168

H-2-Kb: 0%

Peptides in group: 134

**H-2-Kb: 8%**

Peptides in group: 283

**H-2-Kb: 63%**

Peptides in group: 125

H-2-Kb: 0%

**Colon.txt**

Peptides in group: 634

H-2-Kb: 0%

Peptides in group: 1536

**H-2-Kb: 79%**

Peptides in group: 681

H-2-Kb: 0%

**Heart.txt**

Peptides in group: 141

H-2-Kb: 0%

Peptides in group: 410

**H-2-Kb: 83%**

**Kidney.txt**

Peptides in group: 588

**H-2-Kb: 6%**

Peptides in group: 1207

**H-2-Kb: 79%**

Peptides in group: 552

H-2-Kb: 0%

Peptides in group: 659

H-2-Kb: 0%

**Liver.txt**

Peptides in group: 2649

**H-2-Kb: 72%**

**Lung.txt**

Peptides in group: 1420

**H-2-Kb: 77%**

**Ovary.txt**

Peptides in group: 156

**H-2-Kb: 85%**

Peptides in group: 47

H-2-Kb: 0%

Peptides in group: 47

**H-2-Kb: 6%**

Peptides in group: 50

H-2-Kb: 0%

**Pancreas.txt**

Peptides in group: 28

H-2-Kb: 0%

Peptides in group: 21

**H-2-Kb: 14%**

Peptides in group: 31

**H-2-Kb: 3%**

Peptides in group: 167

**H-2-Kb: 82%**

Peptides in group: 23

H-2-Kb: 0%

**Skin.txt**

Peptides in group: 631

**H-2-Kb: 83%**

**SpinalCord.txt**

Peptides in group: 18

H-2-Kb: 0%

Peptides in group: 26

**H-2-Kb: 23%**

Peptides in group: 17

**H-2-Kb: 6%**

Peptides in group: 10

**H-2-Kb: 40%**

Peptides in group: 5

H-2-Kb: 0%

**Spleen.txt**

Peptides in group: 1956

**H-2-Kb: 84%**

**Testis.txt**

Peptides in group: 728

**H-2-Kb: 76%**

**Thymus.txt**

Peptides in group: 1262

**H-2-Kb: 85%**

**Uterus.txt**

Peptides in group: 1293

**H-2-Kb: 83%**

**Intestine.txt**

Peptides in group: 741

**H-2-Kb: 2%**

Peptides in group: 1880

**H-2-Kb: 72%**

Peptides in group: 948

**H-2-Kb: 1%**

Peptides in group: 951

H-2-Kb: 0%

Peptides in group: 904

H-2-Kb: 0%

**Stomach.txt**

Peptides in group: 862

**H-2-Kb: 1%**

Peptides in group: 881

**H-2-Kb: 26%**

Peptides in group: 749

**H-2-Kb: 1%**

Peptides in group: 810

**H-2-Kb: 1%**

Peptides in group: 739

**H-2-Kb: 1%**

**AdrenalGland.txt sequence motif(s)**

**H-2-Kb**

Peptides: 357

**Non-binders group 3**

Peptides: 28

**Non-binders group 1**

Peptides: 35

**Non-binders group 5**

Peptides: 40

**Non-binders group 2**

Peptides: 21

**Non-binders group 4**

Peptides: 12

**Bladder.txt sequence motif(s)**

**H-2-Kb**

Peptides: 430

**Non-binders group 2**

Peptides: 75

**Non-binders group 3**

Peptides: 53

**Non-binders group 4**

Peptides: 66

**Non-binders group 1**

Peptides: 50

**BoneMarrow.txt sequence motif(s)**

**H-2-Kb**

Peptides: 828

**Non-binders group 2**

Peptides: 50

**Non-binders group 3**

Peptides: 59

**Non-binders group 4**

Peptides: 41

**Non-binders group 1**

Peptides: 47

**Brain.txt sequence motif(s)**

**H-2-Kb**

Peptides: 223

**Non-binders group 3**

Peptides: 115

**Non-binders group 1**

Peptides: 168

**Non-binders group 5**

Peptides: 148

**Non-binders group 2**

Peptides: 142

**Non-binders group 4**

Peptides: 123

**Colon.txt sequence motif(s)**

**H-2-Kb**

Peptides: 1378

**Non-binders group 3**

Peptides: 334

**Non-binders group 1**

Peptides: 316

**Non-binders group 5**

Peptides: 263

**Non-binders group 2**

Peptides: 350

**Non-binders group 4**

Peptides: 356

**Heart.txt sequence motif(s)**

**H-2-Kb**

Peptides: 357

**Non-binders group 2**

Peptides: 65

**Non-binders group 1**

Peptides: 79

**Non-binders group 3**

Peptides: 79

**Kidney.txt sequence motif(s)**

**H-2-Kb**

Peptides: 1097

**Non-binders group 3**

Peptides: 357

**Non-binders group 1**

Peptides: 403

**Non-binders group 5**

Peptides: 417

**Non-binders group 2**

Peptides: 400

**Non-binders group 4**

Peptides: 438

**Liver.txt sequence motif(s)**

**H-2-Kb**

Peptides: 2176

**Non-binders group 3**

Peptides: 188

**Non-binders group 1**

Peptides: 201

**Non-binders group 5**

Peptides: 246

**Non-binders group 2**

Peptides: 222

**Non-binders group 4**

Peptides: 252

**Lung.txt sequence motif(s)**

**H-2-Kb**

Peptides: 1186

**Non-binders group 3**

Peptides: 139

**Non-binders group 1**

Peptides: 99

**Non-binders group 5**

Peptides: 146

**Non-binders group 2**

Peptides: 93

**Non-binders group 4**

Peptides: 100

**Ovary.txt sequence motif(s)**

**H-2-Kb**

Peptides: 152

**Non-binders group 2**

Peptides: 34

**Non-binders group 3**

Peptides: 41

**Non-binders group 4**

Peptides: 41

**Non-binders group 1**

Peptides: 37

**Pancreas.txt sequence motif(s)**

**H-2-Kb**

Peptides: 155

**Non-binders group 3**

Peptides: 13

**Non-binders group 1**

Peptides: 22

**Non-binders group 5**

Peptides: 42

**Non-binders group 2**

Peptides: 15

**Non-binders group 4**

Peptides: 20

**Skin.txt sequence motif(s)**

**H-2-Kb**

Peptides: 565

**Non-binders group 3**

Peptides: 19

**Non-binders group 1**

Peptides: 32

**Non-binders group 5**

Peptides: 12

**Non-binders group 2**

Peptides: 26

**Non-binders group 4**

Peptides: 46

**SpinalCord.txt sequence motif(s)**

**H-2-Kb**

Too few peptides to cluster

**Non-binders group 2**

Peptides: 11

**Non-binders group 3**

Peptides: 16

**Non-binders group 4**

Peptides: 10

**Non-binders group 1**

Peptides: 26

**Spleen.txt sequence motif(s)**

**H-2-Kb**

Peptides: 1786

**Non-binders group 3**

Peptides: 94

**Non-binders group 1**

Peptides: 86

**Non-binders group 5**

Peptides: 79

**Non-binders group 2**

Peptides: 74

**Non-binders group 4**

Peptides: 95

**Testis.txt sequence motif(s)**

**H-2-Kb**

Peptides: 597

**Non-binders group 2**

Peptides: 79

**Non-binders group 3**

Peptides: 79

**Non-binders group 4**

Peptides: 84

**Non-binders group 1**

Peptides: 80

**Thymus.txt sequence motif(s)**

**H-2-Kb**

Peptides: 1145

**Non-binders group 3**

Peptides: 78

**Non-binders group 1**

Peptides: 70

**Non-binders group 5**

Peptides: 62

**Non-binders group 2**

Peptides: 86

**Non-binders group 4**

Peptides: 85

**Uterus.txt sequence motif(s)**

**H-2-Kb**

Peptides: 1168

**Non-binders group 3**

Peptides: 72

**Non-binders group 1**

Peptides: 79

**Non-binders group 5**

Peptides: 92

**Non-binders group 2**

Peptides: 75

**Non-binders group 4**

Peptides: 82

**Intestine.txt sequence motif(s)**

**H-2-Kb**

Peptides: 1597

**Non-binders group 3**

Peptides: 725

**Non-binders group 1**

Peptides: 719

**Non-binders group 5**

Peptides: 840

**Non-binders group 2**

Peptides: 877

**Non-binders group 4**

Peptides: 678

**Stomach.txt sequence motif(s)**

**H-2-Kb**

Peptides: 423

**Non-binders group 3**

Peptides: 799

**Non-binders group 1**

Peptides: 741

**Non-binders group 5**

Peptides: 757

**Non-binders group 2**

Peptides: 613

**Non-binders group 4**

Peptides: 730
