## Supplementary material for "Immunopeptidomics for Dummies: Detailed Experimental Protocols and Rapid, User-Friendly Visualization of MHC I and II Ligand Datasets with MhcVizPipe": All supplementary data and tables: S14_MVP report H2Kb_PSCP.html

MhcVizPipe Report


# M

### hc

# V

### iz

# P

### ipe

### - Analysis report

---

**Date:** 2020-09-02

**Submitted by:** Anonymous

**Analysis type:** Class I

**Description of experiment:**

Spliced H2-Kb

**Alleles:** H-2-Kb

**Samples:**

ADRENAL.txt: ADRENAL.txt
BLADDER.txt: BLADDER.txt
BONE\_MARROW.txt: BONE\_MARROW.txt
BRAIN.txt: BRAIN.txt
COLON.txt: COLON.txt
heart.txt: heart.txt
kidney.txt: kidney.txt
Liver.txt: Liver.txt
lung.txt: lung.txt
ovary.txt: ovary.txt
pancreas.txt: pancreas.txt
skin.txt: skin.txt
small\_intestine.txt: small\_intestine.txt
spinal\_cord.txt: spinal\_cord.txt
spleen.txt: spleen.txt
stomach.txt: stomach.txt
testis.txt: testis.txt
Thymus.txt: Thymus.txt
uterus.txt: uterus.txt

---

### Sample Overview

**UpSet Plot**

Only intersections > 0.5% are displayed

**Peptide Length Distribution** (maximum of 30 mers)

| Sample | Peptide length | Total peptides | % |
| --- | --- | --- | --- |
| ADRENAL.txt | all lengths | 23 | 100 |
|  | 8-12 mers | 23 | 100 |
|  | other | 0 | 0 |
| BLADDER.txt | all lengths | 25 | 100 |
|  | 8-12 mers | 25 | 100 |
|  | other | 0 | 0 |
| BONE\_MARROW.txt | all lengths | 64 | 100 |
|  | 8-12 mers | 64 | 100 |
|  | other | 0 | 0 |
| BRAIN.txt | all lengths | 45 | 100 |
|  | 8-12 mers | 45 | 100 |
|  | other | 0 | 0 |
| COLON.txt | all lengths | 119 | 100 |
|  | 8-12 mers | 119 | 100 |
|  | other | 0 | 0 |
| heart.txt | all lengths | 25 | 100 |
|  | 8-12 mers | 25 | 100 |
|  | other | 0 | 0 |
| kidney.txt | all lengths | 152 | 100 |
|  | 8-12 mers | 152 | 100 |
|  | other | 0 | 0 |
| Liver.txt | all lengths | 338 | 100 |
|  | 8-12 mers | 338 | 100 |
|  | other | 0 | 0 |
| lung.txt | all lengths | 164 | 100 |
|  | 8-12 mers | 164 | 100 |
|  | other | 0 | 0 |
| ovary.txt | all lengths | 15 | 100 |
|  | 8-12 mers | 15 | 100 |
|  | other | 0 | 0 |
| pancreas.txt | all lengths | 9 | 100 |
|  | 8-12 mers | 9 | 100 |
|  | other | 0 | 0 |
| skin.txt | all lengths | 42 | 100 |
|  | 8-12 mers | 42 | 100 |
|  | other | 0 | 0 |
| small\_intestine.txt | all lengths | 217 | 100 |
|  | 8-12 mers | 217 | 100 |
|  | other | 0 | 0 |
| spinal\_cord.txt | all lengths | 4 | 100 |
|  | 8-12 mers | 4 | 100 |
|  | other | 0 | 0 |
| spleen.txt | all lengths | 188 | 100 |
|  | 8-12 mers | 188 | 100 |
|  | other | 0 | 0 |
| stomach.txt | all lengths | 41 | 100 |
|  | 8-12 mers | 41 | 100 |
|  | other | 0 | 0 |
| testis.txt | all lengths | 49 | 100 |
|  | 8-12 mers | 49 | 100 |
|  | other | 0 | 0 |
| Thymus.txt | all lengths | 69 | 100 |
|  | 8-12 mers | 69 | 100 |
|  | other | 0 | 0 |
| uterus.txt | all lengths | 79 | 100 |
|  | 8-12 mers | 79 | 100 |
|  | other | 0 | 0 |

---

### Annotation Results

NetMHCpan eluted ligand predictions made for all peptides between 8 & 12 mers, inclusive.
Percent rank cutoffs for strong and weak binders: 0.5 and 2.0.

| Allele | Sample | Total peptides | Strong binders | Weak binders |
| --- | --- | --- | --- | --- |
| H-2-Kb | ADRENAL.txt | 23 | 15 (65.2%) | 1 (4.3%) |
|  | BLADDER.txt | 25 | 17 (68.0%) | 1 (4.0%) |
|  | BONE\_MARROW.txt | 64 | 43 (67.2%) | 5 (7.8%) |
|  | BRAIN.txt | 45 | 11 (24.4%) | 4 (8.9%) |
|  | COLON.txt | 119 | 87 (73.1%) | 11 (9.2%) |
|  | heart.txt | 25 | 22 (88.0%) | 0 (0.0%) |
|  | kidney.txt | 152 | 82 (53.9%) | 13 (8.6%) |
|  | Liver.txt | 338 | 266 (78.7%) | 35 (10.4%) |
|  | lung.txt | 164 | 108 (65.9%) | 11 (6.7%) |
|  | ovary.txt | 15 | 5 (33.3%) | 1 (6.7%) |
|  | pancreas.txt | 9 | 5 (55.6%) | 0 (0.0%) |
|  | skin.txt | 42 | 34 (81.0%) | 2 (4.8%) |
|  | small\_intestine.txt | 217 | 130 (59.9%) | 26 (12.0%) |
|  | spinal\_cord.txt | 4 | 0 (0.0%) | 0 (0.0%) |
|  | spleen.txt | 188 | 153 (81.4%) | 18 (9.6%) |
|  | stomach.txt | 41 | 12 (29.3%) | 5 (12.2%) |
|  | testis.txt | 49 | 32 (65.3%) | 3 (6.1%) |
|  | Thymus.txt | 69 | 56 (81.2%) | 1 (1.4%) |
|  | uterus.txt | 79 | 59 (74.7%) | 7 (8.9%) |

**Binding Affinities**

---

### Binding Heatmaps

---

### Sequence Motifs

Clustering performed with all peptides between 8 & 12 mers, inclusive.

Percentages represent the percentage of peptides in a given group predicted to strongly bind the indicated allele.

Polar

Neutral

Basic

Acidic

Hydrophobic

- Unsupervised GibbsCluster
- Allele-specific GibbsCluster

**BRAIN.txt**

Peptides in group: 2

H-2-Kb: 0%

Peptides in group: 16

**H-2-Kb: 56%**

Peptides in group: 11

H-2-Kb: 0%

Peptides in group: 14

**H-2-Kb: 14%**

**kidney.txt**

Peptides in group: 20

H-2-Kb: 0%

Peptides in group: 34

**H-2-Kb: 18%**

Peptides in group: 2

H-2-Kb: 0%

Peptides in group: 90

**H-2-Kb: 84%**

**lung.txt**

Peptides in group: 4

H-2-Kb: 0%

Peptides in group: 121

**H-2-Kb: 84%**

Peptides in group: 9

H-2-Kb: 0%

Peptides in group: 29

**H-2-Kb: 21%**

**ADRENAL.txt**

Peptides in group: 17

**H-2-Kb: 88%**

Peptides in group: 4

H-2-Kb: 0%

**BLADDER.txt**

Peptides in group: 3

H-2-Kb: 0%

Peptides in group: 20

**H-2-Kb: 85%**

**BONE\_MARROW.txt**

Peptides in group: 2

H-2-Kb: 0%

Peptides in group: 55

**H-2-Kb: 78%**

**stomach.txt**

Peptides in group: 14

H-2-Kb: 0%

Peptides in group: 13

**H-2-Kb: 85%**

**Thymus.txt**

Peptides in group: 59

**H-2-Kb: 95%**

Peptides in group: 9

H-2-Kb: 0%

**COLON.txt**

Peptides in group: 109

**H-2-Kb: 80%**

**heart.txt**

Peptides in group: 24

**H-2-Kb: 92%**

**Liver.txt**

Peptides in group: 326

**H-2-Kb: 81%**

**ovary.txt**

Peptides in group: 9

**H-2-Kb: 56%**

**pancreas.txt**

Peptides in group: 5

**H-2-Kb: 100%**

**skin.txt**

Peptides in group: 40

**H-2-Kb: 85%**

**small\_intestine.txt**

Peptides in group: 184

**H-2-Kb: 71%**

**spinal\_cord.txt**

Peptides in group: 2

H-2-Kb: 0%

**spleen.txt**

Peptides in group: 185

**H-2-Kb: 83%**

**testis.txt**

Peptides in group: 35

**H-2-Kb: 91%**

**uterus.txt**

Peptides in group: 74

**H-2-Kb: 80%**

**BRAIN.txt sequence motif(s)**

**H-2-Kb**

Too few peptides to cluster

**Non-binders group 1**

Peptides: 12

**kidney.txt sequence motif(s)**

**H-2-Kb**

Peptides: 93

**Non-binders group 1**

Peptides: 33

**lung.txt sequence motif(s)**

**H-2-Kb**

Peptides: 118

**Non-binders group 1**

Peptides: 20

**Non-binders group 2**

Peptides: 21

**ADRENAL.txt sequence motif(s)**

**H-2-Kb**

Too few peptides to cluster

**BLADDER.txt sequence motif(s)**

**H-2-Kb**

Too few peptides to cluster

**BONE\_MARROW.txt sequence motif(s)**

**H-2-Kb**

Peptides: 48

**stomach.txt sequence motif(s)**

**H-2-Kb**

Too few peptides to cluster

**Non-binders group 1**

Peptides: 9

**Non-binders group 2**

Peptides: 2

**Thymus.txt sequence motif(s)**

**H-2-Kb**

Peptides: 57

**COLON.txt sequence motif(s)**

**H-2-Kb**

Peptides: 97

**Non-binders group 1**

Peptides: 7

**heart.txt sequence motif(s)**

**H-2-Kb**

Peptides: 22

**Liver.txt sequence motif(s)**

**H-2-Kb**

Peptides: 298

**Non-binders group 1**

Peptides: 15

**Non-binders group 2**

Peptides: 9

**Non-binders group 3**

Peptides: 6

**ovary.txt sequence motif(s)**

**H-2-Kb**

Too few peptides to cluster

**pancreas.txt sequence motif(s)**

**H-2-Kb**

Too few peptides to cluster

**skin.txt sequence motif(s)**

**H-2-Kb**

Peptides: 36

**small\_intestine.txt sequence motif(s)**

**H-2-Kb**

Peptides: 154

**Non-binders group 1**

Peptides: 9

**Non-binders group 2**

Peptides: 5

**Non-binders group 3**

Peptides: 14

**Non-binders group 4**

Peptides: 11

**Non-binders group 5**

Peptides: 19

**spinal\_cord.txt sequence motif(s)**

**H-2-Kb**

Too few peptides to cluster

**spleen.txt sequence motif(s)**

**H-2-Kb**

Peptides: 171

**testis.txt sequence motif(s)**

**H-2-Kb**

Peptides: 35

**uterus.txt sequence motif(s)**

**H-2-Kb**

Peptides: 66
