## Supplementary material for "Immunopeptidomics for Dummies: Detailed Experimental Protocols and Rapid, User-Friendly Visualization of MHC I and II Ligand Datasets with MhcVizPipe": All supplementary data and tables: S15_MVP Report canonical cancer cell line H2Db.htm

MhcVizPipe Report


# M

### hc

# V

### iz

# P

### ipe

### - Analysis report

---

**Date:** 2020-09-02

**Submitted by:** Datasets reprocessed from Caron et al. Sci data 2015

**Analysis type:** Class I

**Description of experiment:**

Canonical peptides Mouse cancer cell lines

**Alleles:** H2-Db

**Samples:**

Lymphoma\_canonical
Lung\_Lewis\_cancer\_Canonical
Malignant\_Glioma\_Canonical
Melanoma\_Canonical

---

### Sample Overview

**UpSet Plot**

Only intersections > 0.5% are displayed

**Peptide Length Distribution** (maximum of 30 mers)

| Sample | Peptide length | Total peptides | % |
| --- | --- | --- | --- |
| Lymphoma\_canonical | all lengths | 2728 | 100 |
|  | 8-12 mers | 2626 | 96 |
|  | other | 102 | 4 |
| Lung\_Lewis\_cancer\_Canonical | all lengths | 1900 | 100 |
|  | 8-12 mers | 1801 | 95 |
|  | other | 99 | 5 |
| Malignant\_Glioma\_Canonical | all lengths | 1105 | 100 |
|  | 8-12 mers | 1034 | 94 |
|  | other | 71 | 6 |
| Melanoma\_Canonical | all lengths | 1568 | 100 |
|  | 8-12 mers | 1498 | 96 |
|  | other | 70 | 4 |

---

### Annotation Results

NetMHCpan eluted ligand predictions made for all peptides between 8 & 12 mers, inclusive.
Percent rank cutoffs for strong and weak binders: 0.5 and 2.0.

| Allele | Sample | Total peptides | Strong binders | Weak binders |
| --- | --- | --- | --- | --- |
| H2-Db | Lymphoma\_canonical | 2626 | 1796 (68.4%) | 225 (8.6%) |
|  | Lung\_Lewis\_cancer\_Canonical | 1801 | 1072 (59.5%) | 190 (10.5%) |
|  | Malignant\_Glioma\_Canonical | 1034 | 631 (61.0%) | 79 (7.6%) |
|  | Melanoma\_Canonical | 1498 | 934 (62.3%) | 133 (8.9%) |

**Binding Affinities**

---

### Binding Heatmaps

---

### Sequence Motifs

Clustering performed with all peptides between 8 & 12 mers, inclusive.

Percentages represent the percentage of peptides in a given group predicted to strongly bind the indicated allele.

Polar

Neutral

Basic

Acidic

Hydrophobic

- Unsupervised GibbsCluster
- Allele-specific GibbsCluster

**Lymphoma\_canonical**

Peptides in group: 2279

**H2-Db: 79%**

**Lung\_Lewis\_cancer\_Canonical**

Peptides in group: 1508

**H2-Db: 71%**

**Malignant\_Glioma\_Canonical**

Peptides in group: 847

**H2-Db: 74%**

**Melanoma\_Canonical**

Peptides in group: 1238

**H2-Db: 75%**

**Lymphoma\_canonical sequence motif(s)**

**H2-Db**

Peptides: 1977

**Non-binders group 1**

Peptides: 178

**Non-binders group 2**

Peptides: 123

**Non-binders group 3**

Peptides: 136

**Non-binders group 4**

Peptides: 117

**Lung\_Lewis\_cancer\_Canonical sequence motif(s)**

**H2-Db**

Peptides: 1228

**Non-binders group 1**

Peptides: 83

**Non-binders group 2**

Peptides: 86

**Non-binders group 3**

Peptides: 106

**Non-binders group 4**

Peptides: 94

**Non-binders group 5**

Peptides: 139

**Malignant\_Glioma\_Canonical sequence motif(s)**

**H2-Db**

Peptides: 695

**Non-binders group 1**

Peptides: 55

**Non-binders group 2**

Peptides: 51

**Non-binders group 3**

Peptides: 77

**Non-binders group 4**

Peptides: 45

**Non-binders group 5**

Peptides: 74

**Melanoma\_Canonical sequence motif(s)**

**H2-Db**

Peptides: 1036

**Non-binders group 1**

Peptides: 78

**Non-binders group 2**

Peptides: 49

**Non-binders group 3**

Peptides: 87

**Non-binders group 4**

Peptides: 77

**Non-binders group 5**

Peptides: 114
