## Supplementary material for "Immunopeptidomics for Dummies: Detailed Experimental Protocols and Rapid, User-Friendly Visualization of MHC I and II Ligand Datasets with MhcVizPipe": All supplementary data and tables: S16_MVP Report PSCP 4 cancer cell lines H2Db.htm

MhcVizPipe Report


# M

### hc

# V

### iz

# P

### ipe

### - Analysis report

---

**Date:** 2020-09-02

**Submitted by:** Anonymous

**Analysis type:** Class I

**Description of experiment:**

PSCP in cancer cell lines

**Alleles:** H2-Db

**Samples:**

Lewis\_lung\_cancer
Lymphoma\_\_EL4\_
Malignant\_Glioma
Melanoma

**Species:** Mouse

---

### Sample Overview

**UpSet Plot**

Only intersections > 0.5% are displayed

**Peptide Length Distribution** (maximum of 30 mers)

| Sample | Peptide length | Total peptides | % |
| --- | --- | --- | --- |
| Lewis\_lung\_cancer | all lengths | 172 | 100 |
|  | 8-12 mers | 172 | 100 |
|  | other | 0 | 0 |
| Lymphoma\_\_EL4\_ | all lengths | 327 | 100 |
|  | 8-12 mers | 327 | 100 |
|  | other | 0 | 0 |
| Malignant\_Glioma | all lengths | 166 | 100 |
|  | 8-12 mers | 166 | 100 |
|  | other | 0 | 0 |
| Melanoma | all lengths | 231 | 100 |
|  | 8-12 mers | 231 | 100 |
|  | other | 0 | 0 |

---

### Annotation Results

NetMHCpan eluted ligand predictions made for all peptides between 8 & 12 mers, inclusive.
Percent rank cutoffs for strong and weak binders: 0.5 and 2.0.

| Allele | Sample | Total peptides | Strong binders | Weak binders |
| --- | --- | --- | --- | --- |
| H2-Db | Lewis\_lung\_cancer | 172 | 60 (34.9%) | 9 (5.2%) |
|  | Lymphoma\_\_EL4\_ | 327 | 200 (61.2%) | 34 (10.4%) |
|  | Malignant\_Glioma | 166 | 56 (33.7%) | 17 (10.2%) |
|  | Melanoma | 231 | 66 (28.6%) | 16 (6.9%) |

**Binding Affinities**

---

### Binding Heatmaps

---

### Sequence Motifs

Clustering performed with all peptides between 8 & 12 mers, inclusive.

Percentages represent the percentage of peptides in a given group predicted to strongly bind the indicated allele.

Polar

Neutral

Basic

Acidic

Hydrophobic

- Unsupervised GibbsCluster
- Allele-specific GibbsCluster

**Lewis\_lung\_cancer**

Peptides in group: 3

H2-Db: 0%

Peptides in group: 22

H2-Db: 0%

Peptides in group: 69

**H2-Db: 12%**

Peptides in group: 76

**H2-Db: 68%**

**Lymphoma\_\_EL4\_**

Peptides in group: 7

H2-Db: 0%

Peptides in group: 29

**H2-Db: 7%**

Peptides in group: 33

**H2-Db: 18%**

Peptides in group: 255

**H2-Db: 75%**

**Malignant\_Glioma**

Peptides in group: 22

**H2-Db: 14%**

Peptides in group: 46

H2-Db: 0%

Peptides in group: 25

**H2-Db: 32%**

Peptides in group: 72

**H2-Db: 62%**

**Melanoma**

Peptides in group: 4

H2-Db: 0%

Peptides in group: 93

H2-Db: 0%

Peptides in group: 42

**H2-Db: 10%**

Peptides in group: 92

**H2-Db: 67%**

**Lewis\_lung\_cancer sequence motif(s)**

**H2-Db**

Peptides: 69

**Non-binders group 1**

Peptides: 3

**Non-binders group 2**

Peptides: 14

**Non-binders group 3**

Peptides: 52

**Non-binders group 4**

Peptides: 2

**Non-binders group 5**

Peptides: 32

**Lymphoma\_\_EL4\_ sequence motif(s)**

**H2-Db**

Peptides: 234

**Non-binders group 1**

Peptides: 29

**Non-binders group 2**

Peptides: 15

**Non-binders group 3**

Peptides: 25

**Non-binders group 4**

Peptides: 20

**Malignant\_Glioma sequence motif(s)**

**H2-Db**

Peptides: 71

**Non-binders group 1**

Peptides: 2

**Non-binders group 2**

Peptides: 26

**Non-binders group 3**

Peptides: 17

**Non-binders group 4**

Peptides: 27

**Non-binders group 5**

Peptides: 19

**Melanoma sequence motif(s)**

**H2-Db**

Peptides: 81

**Non-binders group 1**

Peptides: 42

**Non-binders group 2**

Peptides: 99
