## Supplementary material for "Immunopeptidomics for Dummies: Detailed Experimental Protocols and Rapid, User-Friendly Visualization of MHC I and II Ligand Datasets with MhcVizPipe": All supplementary data and tables: S17_MVP Report canonical cancer cell lines H2Kb.htm

MhcVizPipe Report


# M

### hc

# V

### iz

# P

### ipe

### - Analysis report

---

**Date:** 2020-09-02

**Submitted by:** Datasets rerpocessed from Caron et al. 2015 Sci data

**Analysis type:** Class I

**Description of experiment:**

Canonical Mouse cancer cell lines H2Kb

**Alleles:** H2-Kb

**Samples:**

Lung\_Lewis\_Cancer\_Canonical
Lymphoma\_EL4\_Canonical
Malignant\_Glioma\_Canonical
Melanoma\_Canonical

---

### Sample Overview

**UpSet Plot**

Only intersections > 0.5% are displayed

**Peptide Length Distribution** (maximum of 30 mers)

| Sample | Peptide length | Total peptides | % |
| --- | --- | --- | --- |
| Lung\_Lewis\_Cancer\_Canonical | all lengths | 1103 | 100 |
|  | 8-12 mers | 893 | 81 |
|  | other | 210 | 19 |
| Lymphoma\_EL4\_Canonical | all lengths | 2127 | 100 |
|  | 8-12 mers | 2031 | 95 |
|  | other | 96 | 5 |
| Malignant\_Glioma\_Canonical | all lengths | 841 | 100 |
|  | 8-12 mers | 769 | 91 |
|  | other | 72 | 9 |
| Melanoma\_Canonical | all lengths | 1172 | 100 |
|  | 8-12 mers | 1124 | 96 |
|  | other | 48 | 4 |

---

### Annotation Results

NetMHCpan eluted ligand predictions made for all peptides between 8 & 12 mers, inclusive.
Percent rank cutoffs for strong and weak binders: 0.5 and 2.0.

| Allele | Sample | Total peptides | Strong binders | Weak binders |
| --- | --- | --- | --- | --- |
| H2-Kb | Lung\_Lewis\_Cancer\_Canonical | 893 | 133 (14.9%) | 65 (7.3%) |
|  | Lymphoma\_EL4\_Canonical | 2031 | 1383 (68.1%) | 248 (12.2%) |
|  | Malignant\_Glioma\_Canonical | 769 | 488 (63.5%) | 62 (8.1%) |
|  | Melanoma\_Canonical | 1124 | 735 (65.4%) | 85 (7.6%) |

**Binding Affinities**

---

### Binding Heatmaps

---

### Sequence Motifs

Clustering performed with all peptides between 8 & 12 mers, inclusive.

Percentages represent the percentage of peptides in a given group predicted to strongly bind the indicated allele.

Polar

Neutral

Basic

Acidic

Hydrophobic

- Unsupervised GibbsCluster
- Allele-specific GibbsCluster

**Lung\_Lewis\_Cancer\_Canonical**

Peptides in group: 172

**H2-Kb: 12%**

Peptides in group: 153

**H2-Kb: 3%**

Peptides in group: 95

H2-Kb: 0%

Peptides in group: 202

**H2-Kb: 53%**

Peptides in group: 218

H2-Kb: 0%

**Lymphoma\_EL4\_Canonical**

Peptides in group: 1776

**H2-Kb: 78%**

**Malignant\_Glioma\_Canonical**

Peptides in group: 612

**H2-Kb: 79%**

**Melanoma\_Canonical**

Peptides in group: 927

**H2-Kb: 79%**

**Lung\_Lewis\_Cancer\_Canonical sequence motif(s)**

**H2-Kb**

Peptides: 197

**Non-binders group 1**

Peptides: 150

**Non-binders group 2**

Peptides: 148

**Non-binders group 3**

Peptides: 105

**Non-binders group 4**

Peptides: 110

**Non-binders group 5**

Peptides: 135

**Lymphoma\_EL4\_Canonical sequence motif(s)**

**H2-Kb**

Peptides: 1603

**Non-binders group 1**

Peptides: 73

**Non-binders group 2**

Peptides: 42

**Non-binders group 3**

Peptides: 77

**Non-binders group 4**

Peptides: 113

**Non-binders group 5**

Peptides: 74

**Malignant\_Glioma\_Canonical sequence motif(s)**

**H2-Kb**

Peptides: 546

**Non-binders group 1**

Peptides: 40

**Non-binders group 2**

Peptides: 57

**Non-binders group 3**

Peptides: 36

**Non-binders group 4**

Peptides: 19

**Non-binders group 5**

Peptides: 52

**Melanoma\_Canonical sequence motif(s)**

**H2-Kb**

Peptides: 807

**Non-binders group 1**

Peptides: 58

**Non-binders group 2**

Peptides: 61

**Non-binders group 3**

Peptides: 66

**Non-binders group 4**

Peptides: 67

**Non-binders group 5**

Peptides: 34
