## Supplementary material for "Immunopeptidomics for Dummies: Detailed Experimental Protocols and Rapid, User-Friendly Visualization of MHC I and II Ligand Datasets with MhcVizPipe": All supplementary data and tables: S18_MVP Report PSCP 4 cancer cell lines H2Kb.htm

MhcVizPipe Report


# M

### hc

# V

### iz

# P

### ipe

### - Analysis report

---

**Date:** 2020-09-02

**Submitted by:** Anonymous

**Analysis type:** Class I

**Description of experiment:**

PSCP in mouse cancer cell line

**Alleles:** H2-Kb

**Samples:**

Lewis\_Lung\_Cancer
Lymphoma\_EL4
Malignant\_Glioma
Melanoma

**Species:** Mouse

---

### Sample Overview

**UpSet Plot**

Only intersections > 0.5% are displayed

**Peptide Length Distribution** (maximum of 30 mers)

| Sample | Peptide length | Total peptides | % |
| --- | --- | --- | --- |
| Lewis\_Lung\_Cancer | all lengths | 22 | 100 |
|  | 8-12 mers | 22 | 100 |
|  | other | 0 | 0 |
| Lymphoma\_EL4 | all lengths | 133 | 100 |
|  | 8-12 mers | 133 | 100 |
|  | other | 0 | 0 |
| Malignant\_Glioma | all lengths | 35 | 100 |
|  | 8-12 mers | 35 | 100 |
|  | other | 0 | 0 |
| Melanoma | all lengths | 76 | 100 |
|  | 8-12 mers | 76 | 100 |
|  | other | 0 | 0 |

---

### Annotation Results

NetMHCpan eluted ligand predictions made for all peptides between 8 & 12 mers, inclusive.
Percent rank cutoffs for strong and weak binders: 0.5 and 2.0.

| Allele | Sample | Total peptides | Strong binders | Weak binders |
| --- | --- | --- | --- | --- |
| H2-Kb | Lewis\_Lung\_Cancer | 22 | 1 (4.5%) | 1 (4.5%) |
|  | Lymphoma\_EL4 | 133 | 95 (71.4%) | 18 (13.5%) |
|  | Malignant\_Glioma | 35 | 28 (80.0%) | 1 (2.9%) |
|  | Melanoma | 76 | 48 (63.2%) | 8 (10.5%) |

**Binding Affinities**

---

### Binding Heatmaps

---

### Sequence Motifs

Clustering performed with all peptides between 8 & 12 mers, inclusive.

Percentages represent the percentage of peptides in a given group predicted to strongly bind the indicated allele.

Polar

Neutral

Basic

Acidic

Hydrophobic

- Unsupervised GibbsCluster
- Allele-specific GibbsCluster

**Lewis\_Lung\_Cancer**

Peptides in group: 8

H2-Kb: 0%

Peptides in group: 5

**H2-Kb: 20%**

Peptides in group: 2

H2-Kb: 0%

**Melanoma**

Peptides in group: 10

**H2-Kb: 10%**

Peptides in group: 60

**H2-Kb: 78%**

**Lymphoma\_EL4**

Peptides in group: 125

**H2-Kb: 76%**

**Malignant\_Glioma**

Peptides in group: 30

**H2-Kb: 93%**

**Lewis\_Lung\_Cancer sequence motif(s)**

**H2-Kb**

Too few peptides to cluster

**Non-binders group 1**

Peptides: 8

**Melanoma sequence motif(s)**

**H2-Kb**

Peptides: 56

**Non-binders group 1**

Peptides: 8

**Lymphoma\_EL4 sequence motif(s)**

**H2-Kb**

Peptides: 113

**Non-binders group 1**

Peptides: 6

**Non-binders group 2**

Peptides: 8

**Malignant\_Glioma sequence motif(s)**

**H2-Kb**

Peptides: 29
